# An isogenic single-cell atlas of familial Parkinson’s disease mutations reveals convergent changes in dopamine neurons

**DOI:** 10.64898/2026.05.04.722458

**Authors:** Khaja Mohieddin Syed, Jesse Dunnack, Sarah Paatz, Krishna Ajjarapu, Atehsa Sahagun, Donald C. Rio, Frank Soldner, Helen S. Bateup, Dirk Hockemeyer

## Abstract

Familial Parkinson’s disease (PD) is caused by mutations in more than twenty genes that affect diverse cellular pathways, including mitochondrial quality control, lysosomal function, and vesicular trafficking. A central question is how mutations impacting these distinct pathways converge to cause the selective degeneration of dopamine neurons. Human pluripotent stem cell (hPSC)-based disease models provide a valuable system to study this; however, systematic comparison of the pathogenic effects of different mutations has been limited by genetic background variability. To address this, we generated an isogenic single-cell transcriptomic atlas of fourteen familial PD mutations comprising more than 200,000 hPSC-derived midbrain specified cells. Integrated analysis revealed mutation-specific transcriptional signatures alongside shared dysregulated genes and modules that converge on mitochondrial homeostasis, endolysosomal degradation, and iron/ferroptosis pathways. Differentially expressed genes were significantly enriched for PD GWAS-implicated genes in dopamine neurons, bridging monogenic and sporadic PD genetic risk and highlighting a shared downstream state across multiple mutations. Finally, cells with a *DNAJC6* mutation, which is associated with juvenile-onset parkinsonism, exhibited alteration of neurodevelopmental and psychiatric disorder risk genes, providing a transcriptional correlate for neurodevelopmental features observed in early-onset PD. Together, this resource enables molecular stratification of familial PD mutations and provides a foundational benchmarking data set.

## Introduction

Parkinson’s disease (PD) is the second most common neurodegenerative disorder and is defined by the progressive loss of midbrain dopamine neurons. The genetic architecture of PD is heterogeneous, with familial mutations accounting for approximately 10% of cases, while the remaining sporadic forms arise from complex gene-environment interactions^1^. Although these familial mutations have identified key pathways implicated in PD, including mitochondrial function, proteostasis, and lysosomal biology, how these well-defined genetic lesions ultimately cause dopamine neuron degeneration remains incompletely understood^2^. A major challenge is the pronounced heterogeneity in clinical presentation and molecular phenotypes, even among individuals carrying the same mutation, which obscures mutation-specific effects and complicates mechanistic interpretation.

Genetically engineered isogenic human pluripotent stem cell (hPSC) models provide a powerful approach to disentangle mutation-specific effects from genetic background by enabling direct comparison of disease-causing variants in an otherwise identical genome^3^. Differentiation of hPSCs into midbrain dopamine neurons, however, is inherently heterogeneous, yielding mixtures of neuronal subtypes and variable maturation states even under optimized protocols^4,5^. As a result, bulk transcriptional profiling obscures cell-type-specific and state-dependent effects of PD mutations, making single-cell resolution essential for accurate mechanistic interpretation. Recent single-cell transcriptomic studies (scRNA-seq) using hPSC-derived dopamine neurons have revealed distinct mutation-specific transcriptional programs in PD^6^. ScRNA-seq of neurons harboring the *SNCA* A53T mutation versus isogenic controls demonstrated metabolic and proteostatic perturbations, including altered glycolytic and cholesterol metabolic programs and disrupted ubiquitin-proteasomal degradation^7^. Analysis of patient-derived *GBA* N370S dopamine neurons identified ER stress with HDAC4 nuclear mislocalization as an upstream driver^8^. *LRRK2* G2019S dopamine neuron models have demonstrated altered differentiation dynamics via NR2F1^9^. While these studies highlight the power of combining isogenic hPSC models with single-cell analysis, they examined individual mutations in isolation. Consequently, we still lack a systematic and scalable approach for comparing multiple familial PD mutations side by side to define shared versus mutation-specific pathways and to resolve mechanistic relationships across genetic lesions.

Direct analysis of PD brain tissue represents the gold standard. However, such analysis reflects late-stage disease^10,11,12^, and does not resolve early pathogenic events or separate primary effects from secondary responses. A comprehensive reference map that integrates multiple PD mutations within an isogenic, developmentally controlled system is therefore needed to define convergent biological signatures and mutation-specific vulnerabilities. To address this need, we generated a large-scale single-cell transcriptomic atlas of dopamine neurons carrying one of fourteen familial PD mutations engineered into a single parental Human embryonic stem cells (hESC) background from the iSCORE-PD collection^13^. This design enables direct comparison of mutation-associated transcriptional programs while controlling for genetic background and capturing differentiation-state heterogeneity at single-cell resolution. Using supervised cell-type prioritization, curated pathway/module scoring, and integration with nominated PD GWAS risk genes, we find both convergent molecular states shared across mutations, and divergent genotype-specific responses. This atlas and accompanying analysis provide a cell type resolved reference resource for interpreting how genetically distinct forms of familial PD signatures converge on shared features in dopamine neurons, and for bridging familial and sporadic disease.

## Results

### Single-cell RNA sequencing of dopamine neurons from the iSCORE-PD collection

To systematically compare transcriptional programs across mutations associated with familial PD, we generated a scRNA-seq atlas of hESC-derived dopamine neurons carrying 14 distinct PD-associated mutations alongside isogenic controls (Fig. 1A). We leveraged the iSCORE-PD (Isogenic Stem Cell Collection to Research PD^13^) resource, composed of cell lines engineered from a single, fully characterized female hESC line, enabling direct comparison of disease-causing mutations in a common genetic background.

**Figure 1:**
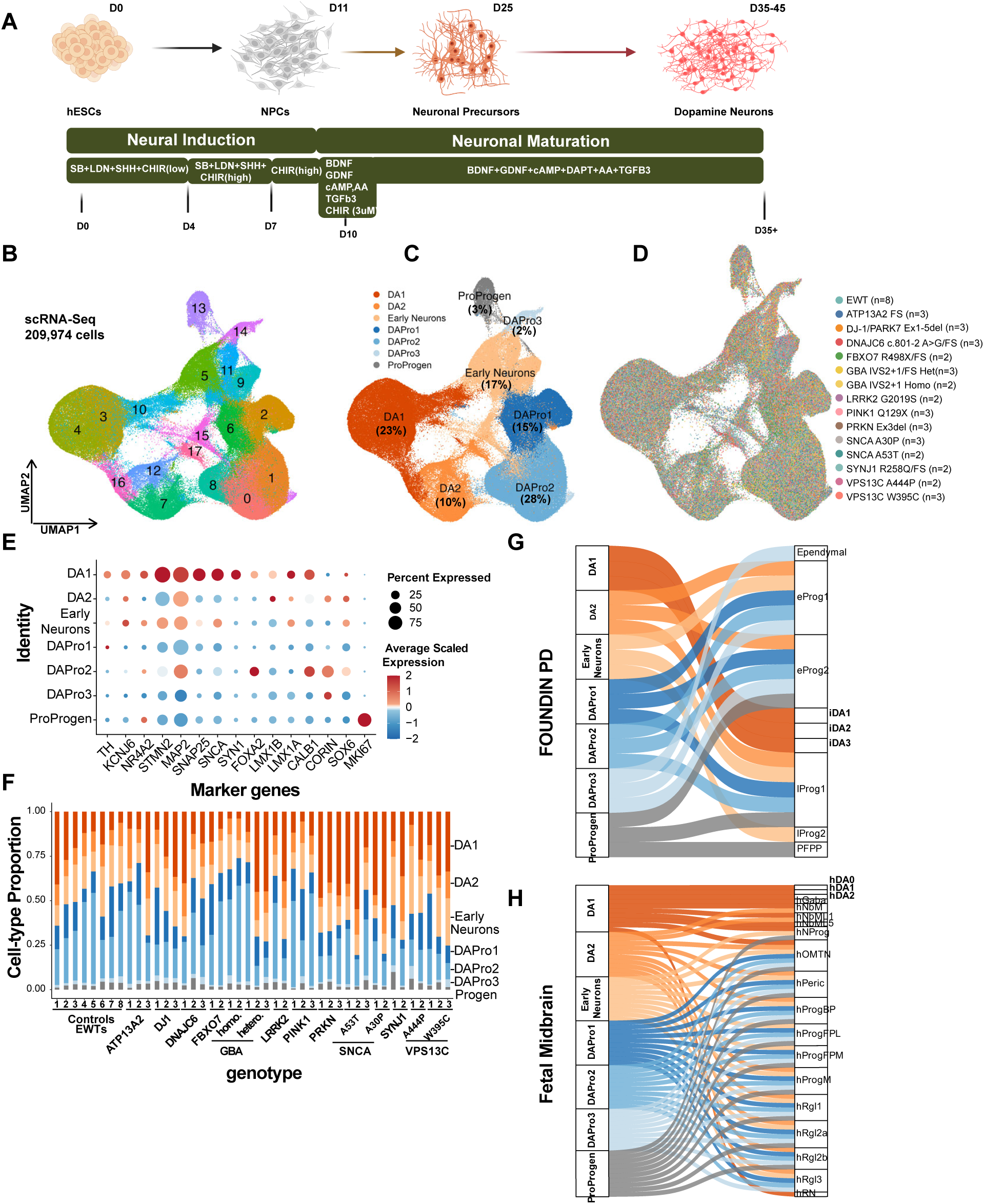
Familial PD single cell transcriptomics profiling and molecular features of pluripotent stem cell derived dopamine neurons. **A)** Schematic of the directed differentiation protocol used to generate ventral midbrain dopamine neurons from human pluripotent stem cells. **(B-D)** Uniform Manifold Approximation and Projection (UMAP) of the integrated dataset comprising 209,974 high quality single cells from cells harvested between days 35 and 45. Cells are colored by (B) unsupervised cluster identity, (C) annotated cell types (DA1, DA2, Early Neurons, DAPro1, DAPro2, DAPro3, and ProProgen; percentage of total cells per type indicated), and (D) genotype (14 familial PD mutations and 8 isogenic Edited Wild-Type (EWT) control subclones; n=number of subclones per genotype indicated) **E)** Dot plot showing average expression of dopamine neuron and progenitor markers for each cell type. Dot color indicates scaled mean expression (blue, low; red, high), and dot size represents the percentage of cells expressing the gene within that cell type. **F)** Cell-type proportions distribution plot showing the robustness of the differentiation platform across lines and mutations. Each color represents the cell type as shown in the annotated UMAP and each bar represents a different cell line used. The plots represent a total of 44 subclones encompassing 14 PD mutations along with control (n=8, EWT control subclones). Colors indicate cell types as in C: Bright orange, DA1; mid bright Orange, DA2; low bright orange, Early Neurons; Bright blue, DAPro1; mid bright blue, DAPro2; low bright blue, DAPro3. Grey, ProProgen. **G-H)** Spearman correlation of our dataset with the *in-vivo* fetal ventral midbrain dopamine neuron single-cell dataset (G; top 10 reference cell types per iSCORE-PD scAtlas population) and *in-vitro* stem-cell derived dopamine neurons from FOUNDIN PD dataset (H; top 3 reference cell types per iSCORE-PD scAtlas population), showing higher correlation of gene expression to our cell types across datasets.

In total, 44 independently derived hESC subclones, comprising 14 mutations across 11 genes and 8 wild-type controls were differentiated using a floor plate-based midbrain protocol^14,15^ and profiled between days 35 and 45 of maturation (Fig. 1A). The controls went through the gene editing pipeline but remained wild-type at the targeted locus (referred to as edited wild-type, hereafter “EWT”). Using the 10x Genomics platform, we obtained 209,974 high-quality single cells with a median of ∼3000 genes detected per cell (and a median of ∼9000 UMIs per cell) across subclones spanning seven differentiation batches following stringent quality control filtering (Fig. S1A-D, TableS1). Sequencing depth and gene detection rates were highly correlated across genotypes and batches (Fig. S1E-F), demonstrating technical consistency across the dataset. Unsupervised clustering and marker-based annotation identified seven major cell populations spanning midbrain progenitors, cycling progenitors, and post-mitotic dopamine neurons (Fig. 1B-D, TableS2). Three dopamine neuron clusters (DA1, DA2, Early Neurons), defined by expression of *TH*, *NR4A2*, and *KCNJ6*, comprised 33.8% of all cells. Three progenitor clusters (DAPro1, DAPro2, DAPro3) were defined by robust expression of FOXA2, a midbrain floor-plate transcription factor and *CORIN,* a canonical floor-plate progenitor marker (Fig. 1E, S2A-B). Minimal expression of forebrain or hindbrain markers confirmed appropriate regional specification (Fig. S2C-D). Cellular composition and differentiation efficiency were generally consistent across the mutant lines and EWT controls (Fig. 1F). No systematic genotype- or batch-driven shifts in cluster representation were observed (Fig, S3A-D). To further confirm our differentiation protocol, we quantified TH⁺/FOXA2⁺ neurons by immunohistochemistry across most subclones and batches (Fig. S4A-C). The percentage of TH⁺ neurons determined by immunostaining showed a positive correlation with the percentage of TH-expressing cells determined by scRNA-seq (Pearson r=0.74) (Fig.S4C). Key neuronal markers (*TH, MAP2, SNAP25, SYN1*) further confirmed consistent neuronal identity across the subclones (Fig. S1G). To benchmark differentiation fidelity, we compared our dataset with published single-cell profiles of midbrain dopamine neurons^16,17,18^. Pairwise correlation analysis demonstrated strong transcriptional concordance between our dopamine neuron populations and previously published in vitro-derived dopamine neurons, human fetal midbrain single cell datasets (Fig. 1G-H, Fig. S3E-F), and adult in vivo human post-mortem midbrain dopamine neurons from the Substantia Nigra (SNc) (Fig. S3G). These results confirm that the iSCORE-PD differentiation platform captures key molecular features of midbrain dopamine neuron identity.

### Cell-type specific transcriptional vulnerability across familial PD mutations

Familial PD risk genes show different cell type expression patterns; some are broadly expressed, while others are known to be expressed in specific cell types^12, 19^. We examined this in our dataset and found that familial PD genes had variable expression patterns across cell populations in control cells, suggesting that mutation-associated effects may vary by cell type (Fig. S5A). This heterogeneity in the baseline expression suggested that the transcriptional impact of each mutation may differ across all cell types, motivating a systematic approach to identify which cell populations are most responsive to each genetic perturbation. To this end, we applied Augur^20^, a machine learning framework that quantifies how well mutant cells can be distinguished from isogenic controls within each cell type, to identify the cell populations exhibiting the strongest transcriptional shifts for each mutation.

We first examined overall perturbation by averaging Augur scores across neuronal cell types (DA1, DA2, Early neurons) and found that *VPS13C*-W395C and *DJ-1/PARK7*-Ex1-5del showed the strongest mean Augur perturbation scores, whereas *SNCA* A53T, *GBA* IVS2+1 homozygous, and *PRKN* Ex3del exhibited more modest scores. At the individual cell type level, *ATP13A2* mutation induced the most pronounced changes in DA1 neurons, while *VPS13C*-W395C and *DJ-1/PARK7* showed stronger perturbations in both early neurons and DA1 cell populations consistent with their known enriched expression in human SNc dopamine neurons^12^.Notably, mutations in genes encoding endolysosomal and vesicular proteins (*VPS13C*, *ATP13A2*, and *LRRK2*) had the highest overall perturbation scores, with *DJ-1/PARK7* likewise also showing high perturbation scores, whereas mutations associated with mitochondrial dysfunction (*PINK1*, *PRKN*) or *SNCA* variants generally showed lower transcriptional perturbation scores (Fig. 2A-S6A). In terms of cell-type-specific effects, *DNAJC6* and *SYNJ1* mutations showed notably higher perturbation score in progenitor populations (DAPro1) than in DA1 neurons, whereas *ATP13A2* and *GBA* IVS2+1 heterozygous induced the largest changes in DA1 neurons (Fig. S6B). From this analysis we conclude that the transcriptional changes caused by familial PD mutations differ in both magnitude and cell type specificity of their transcriptional impact.

**Figure 2:**
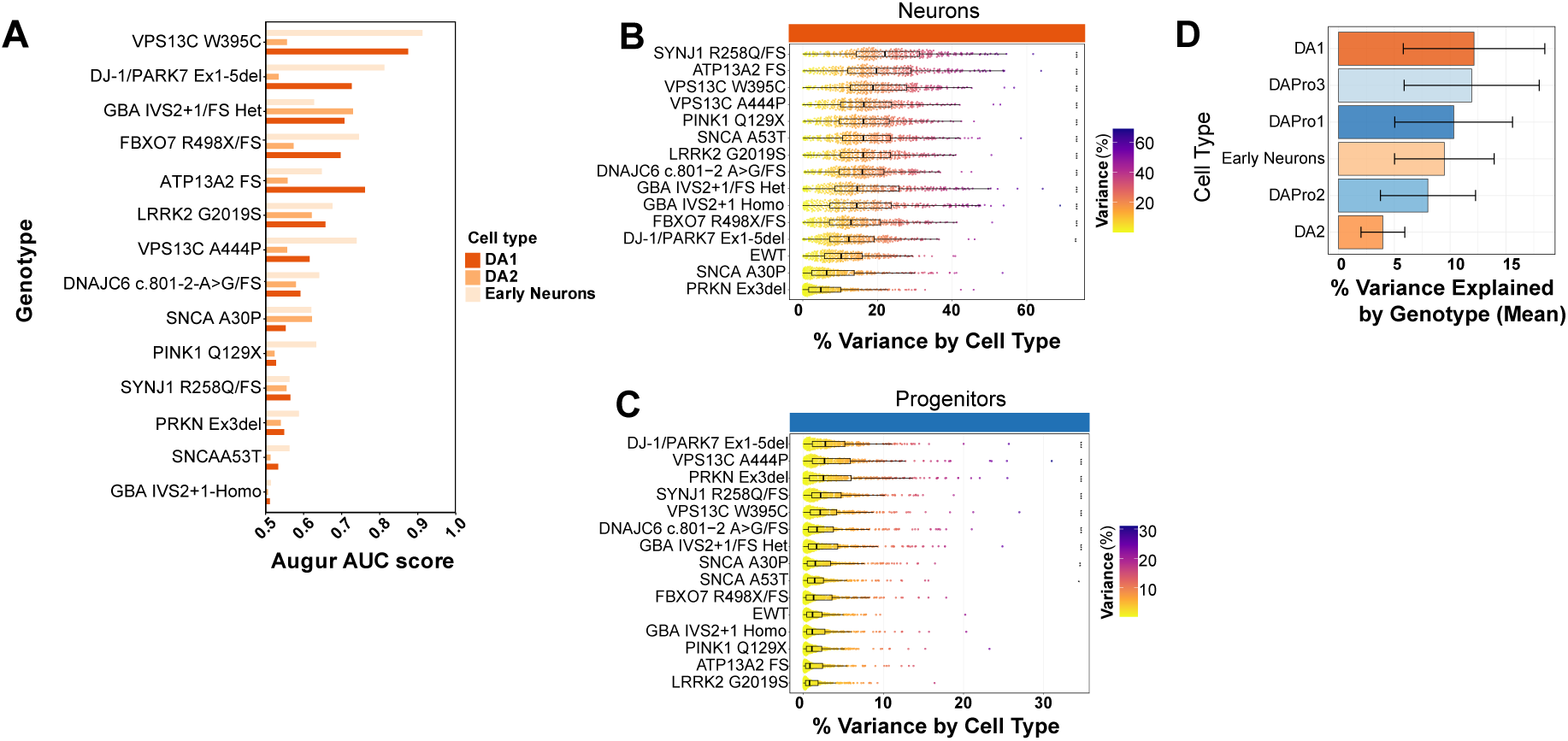
Systematic characterization of mutation-induced transcriptional perturbations in dopaminergic populations. **A)** Cell-type-level mutation responsiveness quantified by Augur analysis. Augur Area Under the Curve (AUC) scores quantifying transcriptional perturbation for DA1, DA2 and Early Neurons populations across all familial PD mutations. Each bar represents the AUC score from batch-stratified random forest classification distinguishing mutation cells from batch-matched controls (EWT) within each cell type. Higher scores indicate stronger mutation-induced disruption. Genotypes are ordered by average AUC scores across the three cell populations. (top to bottom: highest to lowest perturbation). **B-C)** Familial PD mutations amplify cell-type-specific TF regulatory divergence in neurons but not progenitors. ANOVA-based variance decomposition quantifying how much cell type identity explains TF activity variance (%) within each genotype, analyzed separately for (B) DA neurons (DA1, DA2, Early Neurons) and (C) DA progenitors (DAPro1, DAPro2, DAPro3). For each genotype, one-way ANOVA was computed per TF with cell type as the grouping variable. Each dot in the beeswarm plot represents one TF’s variance (up to 298 TF’s per genotype), colored by magnitude (viridis scale). Genotypes are ordered by median variance(ascending). Overlaid boxplots display interquartile range (box) and whiskers; black lines indicate median variance. Asterisks denote significant increases in variance relative to EWT control (one-sided Wilcoxon rank-sum test, ***p<0.001, **p<0.01, *p<0.05). Higher values indicate greater mutation induced divergence between cell subtypes within that compartment. Note the contrast between panels: DA neurons (B) show substantially higher variance and greater mutation-induced shifts than DA progenitors (C) indicating developmental-stage dependent vulnerability. **D)** Cell-type-level TF activity variance attributable to genetic background. Horizontal bar plot showing mean percentage of TF activity variance explained by genotype for each cell type. For each of 6 cell types (ProProgen excluded), one-way ANOVA was performed per TF with genotype as the grouping variable. Bars represent the mean variance per cell type, ordered by magnitude. Error bars indicate SD across TFs. Higher values indicate that TF activity in that cell type is more dependent on genetic background. Note that genotype includes EWT controls; variance therefore reflects total genotype-dependent variation rather than mutation effects alone.

To examine the transcriptional regulatory mechanisms underlying these perturbation patterns, we inferred transcription factor (TF) activity across all mutations in a cell type-specific manner using DoRothEA/VIPER regulon analysis^21,22^(Fig. S7A, Supplemental data Table S3). If familial PD mutations disrupt the core regulatory networks, we reasoned that mutant lines should exhibit greater variation in TF activity across cell types compared to isogenic controls. Indeed, TF activity variance analysis across neuronal populations (DA1, DA2, Early neurons) confirmed that *SYNJ1* and *ATP13A2* caused the highest variance in TF regulatory programs and *SNCA* A30P and *PRKN* exhibiting variance closest to EWT (Fig. 2B). In contrast, progenitor populations showed substantially lower variance across genotypes (Fig. 2C, Supplemental data Table S4), among neuronal cell types was driven primarily by DA1 neurons (Fig. 2D, Supplemental data Table S4). The top 10 upregulated and downregulated TFs per genotype across the three neuronal cell types are shown in Fig. S7B-D. This pattern indicates that familial PD mutations preferentially reshape transcriptional regulatory networks in dopamine neurons populations rather than in progenitors.

### Cell type specific pathway and biological processes enrichment analysis reveals transcriptional signatures across iSCORE-PD genotypes

To characterize the transcriptional changes induced by familial PD mutations, we performed unbiased differential expression (DE) testing using MAST^23^ comparing each mutant genotype to its batch-matched isogenic control within each cell type (Supplemental data Table S5). Transcriptional changes showed a broad spectrum of genotype specific responses, the majority of which were modest in magnitude (log2FC typically <0.5), indicating that the mutations induced subtle shifts in cellular state rather than overt transcriptional rewiring. The observed changes were frequently context dependent, varying across neurons vs progenitors and genotypes, and were influenced by differentiation batch, which can introduce correlated transcriptional shifts despite computational integration. To account for this, DE analysis was performed against batch-matched EWT control cells present in each differentiation batch.

To gain a broader view of the biological pathways underlying these gene expression changes, we performed gene set enrichment analysis (GSEA) on differentially expressed (DE) genes across all 14 mutations (Supplemental data Table S6). We highlight *DJ-1/PARK7* (Ex1-5del) as a representative genotype because of its well-established role in mitochondrial homeostasis and oxidative stress sensing^24^. Differential expression analysis of *DJ-1/PARK7* (Ex1-5del) cells identified robust transcriptional responses in DA1 and early neuron cell populations, whereas DA2 neurons showed a comparatively weak signature, consistent with their lower Augur perturbation scores (see Fig. 3A). KEGG pathway analysis revealed oxidative phosphorylation as the most significantly enriched pathway in both DA1 and early neurons harboring the *DJ-1/PARK7* (Ex1-5del) mutation (Fig. 3B). Proteasome, lysosome, pyruvate metabolism, fatty acid metabolism, and tricarboxylic cycle were consistently upregulated across both cell types, whereas ribosome was consistently downregulated. GO Biological Process analysis extended these findings (Fig. 3C). In *DJ-1/PARK7* DA1 neurons, significant enrichment was observed for mitochondrial electron transport from NADH to ubiquinone, oxidative phosphorylation, proton motive force-driven ATP synthesis, and ATP synthesis-coupled electron transport (Fig. 3C). Upregulation of these mitochondrial respiratory programs is consistent with an oxidative stress-associated transcriptional response to mitochondrial dysfunction following *DJ-1* loss. Additional upregulated terms included chaperone-mediated protein folding, mitochondrial translation and synaptic vesicle lumen acidification, whereas ribosome biogenesis and rRNA processing were downregulated. Equivalent GSEA analyses for the other 13 familial PD mutations are provided in Supplementary Fig. S8-S20.

**Figure 3:**
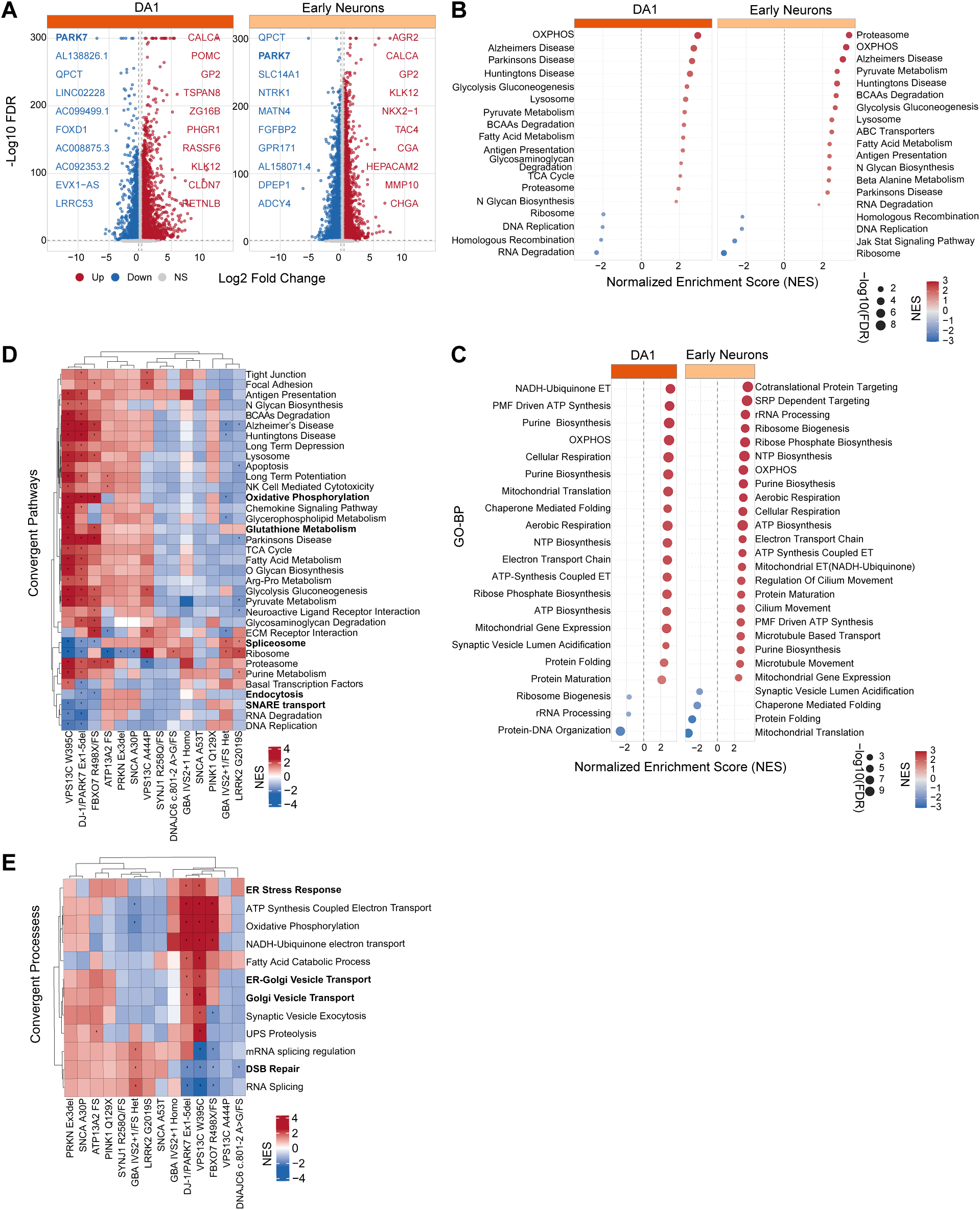
Cell-type-specific differential expression and pathway enrichment across Familial PD mutations. **A)** Volcano plots showing DE results for DA1 and Early Neuron cell type for DJ-1/PARK7 Ex1-5del genotype. X-axis shows average log2 fold change (log2FC); Y-axis shows -log10 (FDR-adjusted p-value). Upregulated genes are in red, downregulated in blue. The top 10 genes per direction per cell type ranked by log2FC are labeled. **B-C)** Dot plots showing Gene Set Enrichment Analysis (GSEA) results in Neuronal populations (DA1, Early Neurons; DA2 is omitted when no gene set passes the false discovery rate (FDR) adjusted p-value <0.05 threshold. Genes were ranked using p-value and log2FC combined, capturing both effect direction and significance. Dot size indicates the number of genes and color indicates the normalized enrichment score. **D-E)** Convergent KEGG pathway and GO-BP enrichment across familial PD mutations in DA1 neurons shown. Heatmaps displaying normalized enrichment score (NES) from GSEA for KEGG pathways and GO biological process terms significantly enriched (Benjamini-Hochberg (BH)-adjusted p<0.05 from adaptive multilevel permutation testing) in at least two genotypes in DA1 neurons. Asterisks indicate significant terms. Rows (pathways) and columns (genotypes) are hierarchically clustered by Pearson correlation distance.

To systematically characterize to what extent the transcriptional signatures for each of the 14 mutation converge in similar pathway-level changes, we examined KEGG and Gene Ontology biological process (GO-BP) enrichment patterns across all mutations in DA1 neurons and identified significantly enriched pathways and terms (FDR <0.05) in atleast two mutations (Fig. 3D-E; Table S6). This analysis showed that *VPS13C*-W395C, *DJ-1/PARK7* and *FBXO7* showed the broadest pathway disruption with the strongest positive enrichment scores across the largest number of pathways (Fig. 3D). PD-relevant KEGG pathways including proteasome and oxidative phosphorylation were dysregulated, showing predominantly upregulation across genotypes. SNARE interactions in vesicular transport and endocytosis were consistently downregulated in *DJ-1/PARK7* and *FBXO7* genotypes. Spliceosome showed bidirectional dysregulation across the genotypes, suggesting context-dependent RNA processing dysregulation. GO-BP analysis identified several dysregulated processes across the 14 PD genotypes, including oxidative phosphorylation, ATP synthesis, and double stranded DNA damage (Fig. 3E). Together, these analyses reveal a complex set of transcriptional changes that extend beyond curated PD pathways. This dataset provides a comprehensive reference atlas that can be interrogated to nominate candidate genes and biological processes for further mechanistic or functional validation. These resource data are provided in Supplemental data Table S1, are also available as raw sequencing files through the GEO repository and processed scRNA data sets can be accessed through the Aligning Parkinson’s Across Research CRN cloud data portal.

### Systematic perturbation profiling reveals convergent transcriptional mechanisms in familial Parkinson’s disease

To systematically identify convergent and divergent transcriptional changes across familial PD mutations, we implemented a directional single-gene AUC ROC framework (see Methods) that classifies genes as preferentially upregulated (AUC >0.5) or downregulated (AUC <0.5) in mutant relative to batch-matched isogenic control cells. We first applied this approach to familial PD genes themselves as validation. As expected, loss-of-function mutations served as internal positive controls: *PINK1*, *PARK7*, *ATP13A2*, and *DNAJC6* mutant cells were accurately classified by reduced transcript abundance of the mutated gene itself in DA1 neurons (Fig. 4A). Beyond the mutated gene, coordinated downstream signatures emerged. *LRRK2*-G2019S cells were distinguished by inverse *SNCA*, *VPS13C*, and *DNAJC6* expression with increased *UCHL1*^25,26^, *VPS13C*-W395C cells showed a complex signature affecting several genes such as *PINK1, ATP13A2 and TMEM230*. These cross-mutation expression patterns suggest that a set of familial PD genes are functionally interconnected, such that perturbation of one gene product can alter the transcriptional state of others, pointing to shared regulatory networks that may underlie the convergence observed across genetically distinct forms of PD.

**Figure 4:**
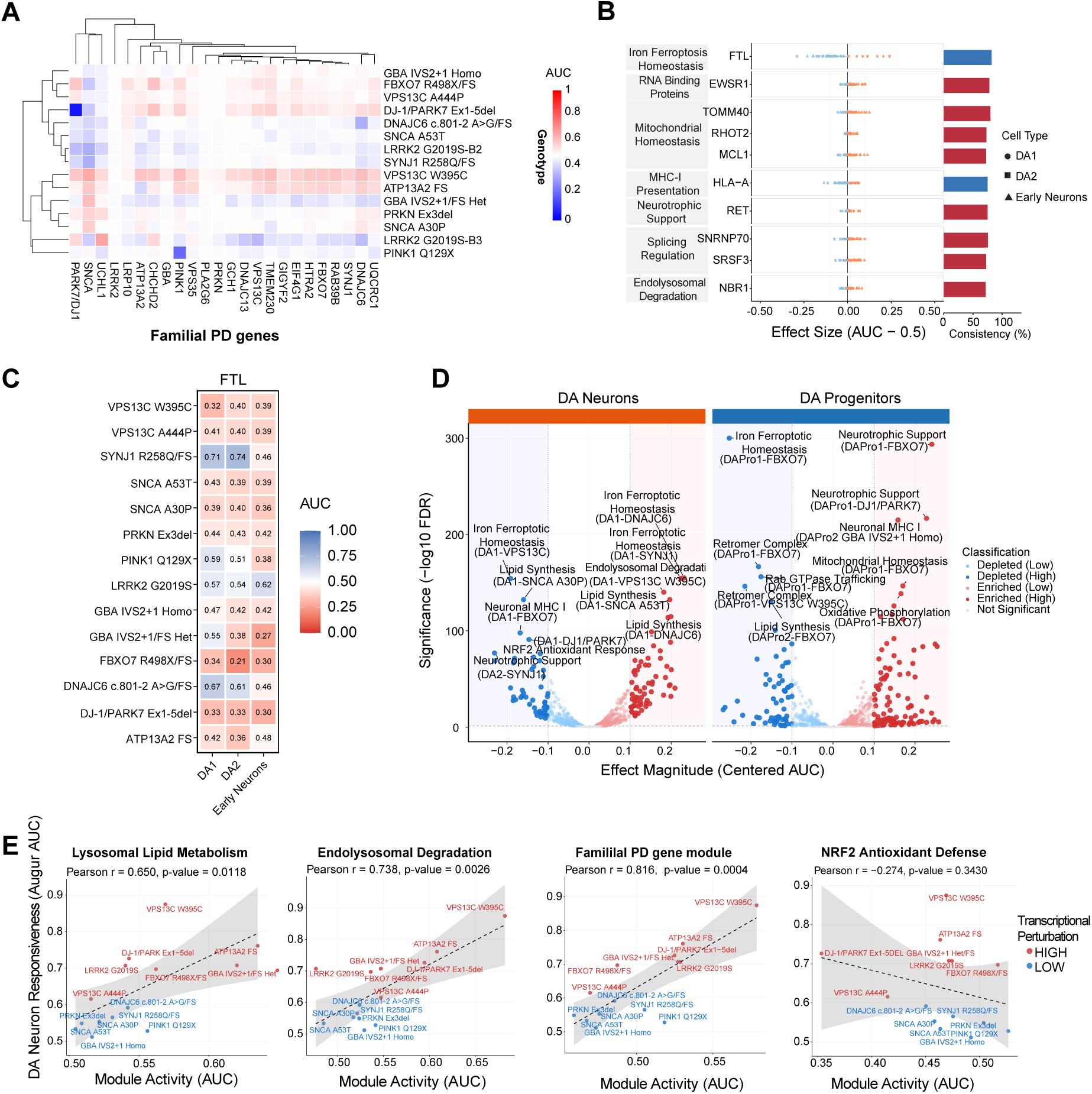
Gene-and module level AUC classification reveals convergent dysregulation across familial PD mutations linked to DA1 transcriptional response. **A)** Single-gene AUC classification reveals mutation-sensitive familial Parkinson’s disease genes in DA1 neurons. Heatmap showing AUC scores for familial PD (FPD) genes across mutations. Each row represents mutation and each column represents genes from the FPD gene set. Blue-white-red scale centered at AUC =0.5 (no change). Red (AUC>0.5) indicates gene upregulation in mutations. Blue (AUC<0.5) indicates gene downregulation, the gene has lower expression in mutant cells. Color intensity reflects magnitude, dark red/ dark blue is strong consistent up/down regulation across all mutant cells respectively; pale colors represent weak or variable effects. **B-C)** Up and down-regulation consistency analysis identifies convergent dysregulation targets across familial PD mutations. Dot plot showing effect sizes (centered AUC, defined as AUC – 0.5, positive values indicate upregulation in mutants and negative values indicate downregulation) for genes meeting at least 70% directional consistency (up or down regulation) across familial PD mutations and three neuronal cell populations (DA1, DA2, Early Neurons). Each dot represents one gene-mutation-cell-type combination, with color indication direction of dysregulation (red: upregulated, AUC>0.5; blue: downregulated, AUC<0.5) and shape indicating cell type (circle: DA1, square: DA2; triangle: Early Neurons). Gray horizontal bars show the mean effect size across all comparisons for each gene. Bars on the right indicate directional consistency (%), defined as the fraction of mutation x cell-type combinations dysregulated in the predominant direction; bar color indicated the predominant direction (red: upregulated; blue: downregulated). Genes are grouped by functional module (labels on the left) and ordered by consistency within each module. Input genes were drawn from a curated panel of >300 PD relevant genes; genes with negligible effect size (SD of AUC<0.01 and mean distance from 0.5<0.02) were excluded prior to ranking. **D)** Volcano plots showing module-level perturbations for PD relevant gene modules. Each point represents one module-genotype-cell type combination. X-axis; effect magnitude (Centered AUC =AUC - 0.5; positive values indicate module enrichment; negative values indicate depletion in mutation cells). Y-axis: statistical significance (-log10 FDR corrected p-value). Points are classified as enriched high (dark red; FDR <0.05, centered AUC >0.1), enriched low (light red; FDR <0.05, centered AUC =<0.1), similarly for depletion on the opposite side. Labels indicate the top 5 modules by significance or effect magnitude. P-values were corrected for multiple testing using Benjamini-Hochberg method across all tests. **E)** Pathway perturbation magnitude scales with cell type responses in DA1 neurons. Scatter plots showing the relationship between pathway module activity (AUC score, x axis) and Augur-derived cell type response score (Y-axis) for the top three correlated pathways in DA1 neurons across 14 familial PD mutations. Each point represents one genotype, colored by response classification: HIGH (red, mean DA neuron Augur AUC above median of 0.63) or LOW (blue, below median). Genotype labels are shown adjacent to each point. Linear regression fit (solid line) with 95% confidence interval (shaded band) is displayed for each pathway. Pearson correlation coefficient (r), p-value. Familial PD gene module (FPD genes) shows the strongest positive correlation with transcriptional response (r=0.816, p=0.0004), Endolysosomal Degradation module positively correlated (r=0.738, p=0.0026), lysosomal lipid metabolism pathways show positive correlation (r=0.65, p=0.0118). NRF2 Antioxidant response shows no association with Augur scores (r=-0.274, p=0.3430), confirming that the observed response associations are selective to specific biological pathways rather than a general feature of transcriptional response.

Having validated this AUC approach using familial PD genes, we applied it over 300 PD associated genes curated into 25 biologically defined modules/gene sets spanning mitochondrial function, endolysosomal degradation, protein quality control, membrane trafficking, cellular stress responses (Supplemental data Table S7), to identify directional convergence across mutations (Fig. S21). Among these genes, *FTL and FTH1*, which encodes the intracellular iron storage protein ferritin light chain, along with select mitochondrial regulators (*TOMM40, COX4I1, COX6C*) were predictive and convergently downregulated across multiple mutations (Fig. 4B; S22, Supplemental data Table S8) ^27,28^. Similarly, TF activity analysis revealed convergent downregulation of RXR-family nuclear receptors across multiple mutations. *RXRB* was suppressed in *ATP13A2, DJ-1/PARK7, and VPS13C* W395C, and *RXRG* in *DJ-1/PARK7*, *FBXO7*, and *VPS13C* W395C in neuronal populations (Fig. S7E-G, S23A-C, Supplemental data Table S9). Given prior evidence linking DJ-1 to ferroptosis suppression and reports that RXR signaling can support neuronal stress resistance^29,30^, the recurrent reduction of RXR receptors TF activity could suggest attenuation of protective lipid and redox regulatory transcriptional programs, potentially including ferroptosis relevant pathways, across genetically distinct familial PD mutations.

Co-suppression of iron storage (*FTL/FTH1*) and lipid protective nuclear receptors (RXRB/RXRG) in the genotypes with the highest Augur perturbation scores (*ATP13A2*, *DJ-1/PARK7*, *FBXO7*, *VPS13C* W395C) prompted us to directly interrogation of the ferroptosis pathway. Core ferroptosis genes (WikiPathways), including *HMOX1*, *NFE2L2*, and *GPX4* were significantly but heterogeneously dysregulated across mutations. TF regulon mapping identified BACH1, ATF4, and SREBF1 as differentially active regulators of these components (Fig. S24A-C). *VPS13C*-W395C and *FBXO7* mutant neurons additionally showed selective upregulation of glutathione metabolism (Fig. 3D-E). Together with convergent FTL downregulation (Fig. 4B-C) and concurrent suppression of RXR nuclear receptor activity (Fig. S23A-B), these findings point to disrupted iron homeostasis as a shared ferroptosis-associated vulnerability across genetically distinct PD mutations.

Having examined individual gene-level changes through the AUC approach, we next asked whether our curated gene sets were enriched or depleted across mutations. Using *Addmodulescore*^31^(see methods), we computed per-cell module scores for each gene set across genotypes and cell types (Supplemental Table S10), excluding the mutated gene from its corresponding gene set to avoid circularity (example, *DJ-1/PARK7* gene was removed from all gene sets when considering DJ-1/PARK7 genotype). In differentiated dopamine neurons, the iron ferroptosis module was the most prominently dysregulated program (Fig. 4D), consistent with the gene-level FTL dysregulation identified in Fig 4B-C. *ATP13A2 and VPS13C* mutations showed enrichment of endolysosomal degradation and lipid biosynthesis modules. The top depleted modules in neurons were related to ferroptosis, lipid biosynthesis, and NRF2 antioxidant support (Fig. 4E, S24D-F). Progenitor populations displayed distinct responses to some of the mutations, with *FBXO7* loss associated with enrichment of neurotrophic support modules and suppression of iron metabolism, suggesting heightened sensitivity to iron-related stress during early differentiation.

To determine which biological processes track with the overall strength of mutation-induced transcriptional disruption, we tested whether module activity correlated with Augur scores across the 14 familial PD mutations in dopamine neurons. Module activity was defined as enrichment or depletion of curated gene sets. Correlating all 25 modules with Augur scores (Fig. 4E) identified strong positive associations for lysosomal lipid metabolism (r=0.650, p=0.0118), familial PD gene expression (r=0.816, p=0.0004), and endolysosomal degradation pathways (r=0.738, p=0.0026), and a significant negative association with the iron ferroptosis homeostasis module (r=-0.56, p=0.036) (Fig. S25A-C). This was highly specific, as all other modules, such as the NRF2 antioxidant defense program, showed no significant correlation with Augur scores (Fig. 4E and S25A-C). Thus, mutations causing the strongest global transcriptional perturbation (*VPS13C*, *ATP13A2*, *DJ-1/PARK7, FBXO7*) also show reduced ferroptosis-protective gene expression, highlighting a shared vulnerability across genetically diverse familial PD mutations.

### Cell-type-specific convergence of familial and sporadic PD risk

To determine whether transcriptional changes induced by familial PD mutations converge on genes linked to sporadic PD, we tested for enrichment of PD GWAS-nominated genes^32,33^(selected from *Open Targets Platform*, with global score >0.1, Supplemental Table S11) within the DE genes identified in our single-cell data sets. Meta-analysis across genotypes and cell types revealed cell-type-specific enrichment of PD GWAS-nominated genes among DE genes, with pronounced heterogeneity across cell populations (Fig. 5A, Supplemental data Table S12). Notably, mature DA1 neurons exhibited the strongest overlap, with significant GWAS-gene enrichment for 10 of 14 mutations (71%; median OR = 1.66), establishing DA1 neurons as a key cell type intersection for monogenic and polygenic PD risk. This aligns with human postmortem SNc analyses demonstrating that the most vulnerable DA neuron subtype is specifically enriched for heritable PD risk^10^, with robust PD GWAS gene expression in iPSC-derived DA neurons^7^ and with the FOUNDIN-PD atlas establishing iPSC-derived DA neurons as a relevant cellular context for modeling PD genetic risk^18^. In contrast, progenitor populations showed progressively weaker enrichment. DAPro1 and DAPro2 displayed moderate convergence (median OR = 1.43 and 1.40; 29% and 43% significant), whereas DAPro3 showed no significant enrichment (0/14 significant), suggesting that the contribution of GWAS-nominated genes to PD risk is most likely mediated through mature dopamine neurons rather than early progenitor states.

**Figure 5:**
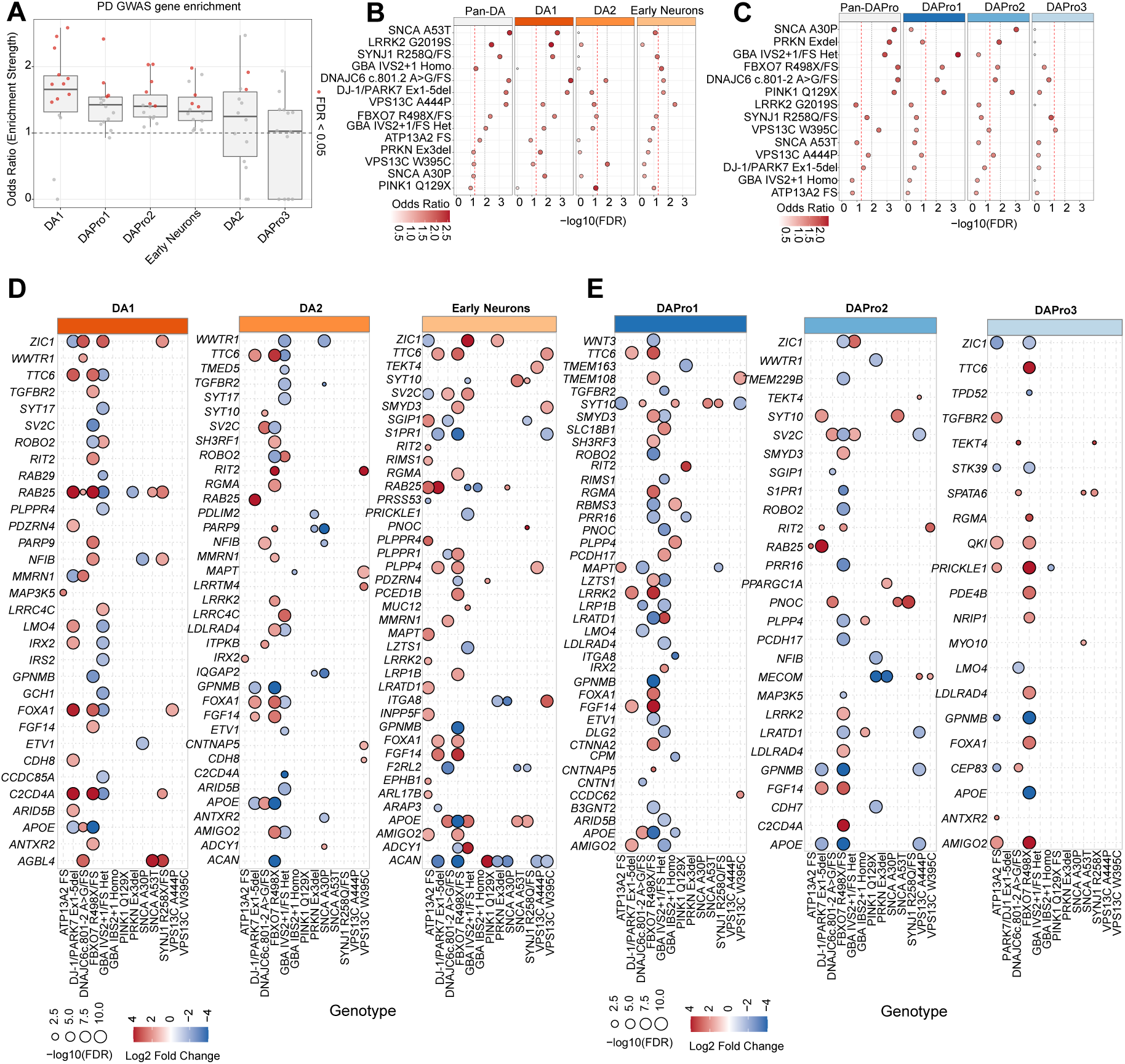
PD GWAS associated genes are significantly enriched among differentially expressed genes across familial PD mutations and cell types. **A)** Cell-type specific enrichment of PD enrichment of PD GWAS risk genes among differentially expressed genes in familial PD mutations. Box plot showing the distribution of Fisher’s exact test odds ratios (Y-axis) across 14 familial PD genotypes for each cell type (x-axis), testing enrichment of PD GWAS-associated genes among differentially expressed genes. PD GWAS genes (n=267) were curated from the Open Targets Platform (v25.09) by requiring GWAS credible set evidence and a global association score >0.1. DE genes were defined with cut off log2FC 0.25 and FDR adjusted p-value < 0.05, calculated with batch matched EWT control. Each dot represents genotype for that cell type; dot color indicates individual test significance (-red= FDR<0.05, grey=NS; BH correction applied globally across all tests). The median line indicates the median odds ratio per cell type. Horizontal dashed line at OR=1 indicates no enrichment (null expectation). Cell types are ordered by median odds ratio (descending order). For LRRK2/G2019S, which was profiled across two batches (B2 and B3), DE genes were defined as those significantly differentially expressed in both batches with the same up or down-regulation change (concordant log2FC directionality). Significance of cell-type-level enrichment was assessed by one-sided Wilcoxon signed-rank test against the null OR=1 across all 14 genotypes. P-values were adjusted using the BH method. **(B-C)** Genotype stratified PD GWAS enrichment **B)** Dot plot showing GWAS enrichment for each of 14 familial PD genotypes across DA1, DA2, Early Neurons, and Pan-DA (Union of DE genes across all three cell types). For each genotype/cell-type combination, a one-sided Fisher’s exact test was assessed for the enrichment of PD GWAS genes. The background universe was defined as all genes tested in each specific comparison. P-values were corrected globally (BH) across all tests in both neuronal and progenitor populations. Genotypes are ordered by Pan-DA odds ratio. **C)** Dot plot showing PD GWAS genes enrichment for DAPro1, DAPro2, DAPro3, and Pan-DAPro (progenitor populations). Note the reduced enrichment compared to Fig. 5B, with fewer genotype/cell-type combinations significant, which is consistent with the developmental stage-dependent vulnerability observed in the TF regulatory analysis (Fig. 2C) **D-E)** Dot plots showing PD GWAS-nominated risk genes that are significantly DE in each mutation across (D) DA neuron and (E) progenitor populations. Only genes meeting the significance thresholds (FDR adjusted p-value <0.05, log2FC cutoff 1.5) are shown.

Genotype-specific analysis in DA1 neurons (Fig. 5B) identified cells with mutations in *DNAJC6*, *SYNJ1*, *LRRK2*, *DJ-1/PARK7*, *SNCA A53T* and *FBXO7* among the strongest enrichments for PD GWAS-associated genes (one-sided Fisher’s exact test, Benjamini-Hochberg (BH)-adjusted p<0.05). This suggests familial PD mutations dysregulate broader networks of sporadic PD risk genes, providing a transcriptional link between monogenic and polygenic disease mechanisms, consistent with the established roles of *SNCA*, *LRRK2*, *DNAJC6* and *SYNJ1* in sporadic PD genetic risk^32,34^. Pan-neuronal analysis (DA1, DA2, and early neurons combined) confirmed *SNCA/A53T*, *DNAJC6* and *DJ-1/PARK7* as the genotypes whose transcriptional changes most significantly overlapped with PD GWAS associated genes (BH adjusted p<0.05), indicating that enrichment rankings are primarily driven by mature neuron effects than the progenitor’s populations (Fig. 5C). We next identified the specific PD GWAS risk genes driving these enrichments across genotypes and cell types (Fig. 5D-E). Among the most consistently dysregulated genes, *APOE* was upregulated across multiple genotypes in neuronal populations. *RAB25*, *FOXA1*, *GPNMB* and *TTC6* were also recurrently dysregulated across genotypes and cell types, identifying them as candidate points of convergence between familial and sporadic PD. In progenitor cells, *APOE, AMIGO2, FGF14, GPNMB and FOXA1* were dysregulated, demonstrating that some GWAS-nominated genes exhibited alterations across cell types and cell developmental stages. To test whether GWAS implicated genes are shared across mutations, we performed pairwise overlap analyses across familial PD mutations. This identified several significant pairs (Fig. S26A-E), among the strongest are the overlaps linking the two lysosomal related mutations *VPS13C*-W395C and *ATP13A2* (85 genes, 73% concordance) (Fig. S26A-B, Supplemental data Table S13). Together these findings suggest that GWAS-implicated genes represent regulatory nodes preferentially engaged by mechanistically distinct familial PD mutations, highlighting a shared vulnerability across genetically diverse familial forms of PD. We next asked whether familial PD mutations altered the expression of high-priority therapeutic target genes nominated by the Micheal J. Fox Foundation (MJFF) Targets to therapies (T2T) initiative. Across 59 T2T genes, several genotypes showed mutation and cell-type dysregulation (Fig.S27A-B). *DNAJC6* and *LRRK2* genotypes showed strong directional concordance in DA1 neurons, with differentially expressed T2T genes shifting in the expected disease associated direction (Fig. S27C). Individual targets, *CDK5* and *KANSL1*, matched T2T predicted expression patterns (Fig. S27D). Together with the GWAS enrichment analysis, these results nominate genotype and cell-type-specific targets associated with PD transcriptional states.

### Familial PD mutations dysregulate neuropsychiatric risk genes across dopamine subpopulations

Several familial PD mutations are associated with juvenile or very early onset parkinsonism^35,36, 37,38^.These genes include *DNAJC6*, *SYNJ1* and *FBXO7*. For individuals carrying mutations in *DNAJC6* and *SYNJ1*, symptom onset often occurs in the first or second decade of life and can include not only classical motor symptoms of PD but also developmental delay or intellectual disability, seizures, and other neuropsychiatric features^35,39^. Likewise, mutations in *FBXO7* cause juvenile/early-onset parkinsonism, typically with prominent parkinsonian and pyramidal motor features, although cognitive or other non-motor neurologic manifestations have also been reported^40,41^. To test whether these juvenile onset forms of PD were associated with distinct gene expression changes, we assessed enrichment of DE genes against established neuropsychiatric gene sets, including two autism spectrum disorder (ASD) datasets from the SFARI database representing non-syndromic and syndromic autism^42^, and schizophrenia (SCZ) and bipolar disorder (BP) risk genes from DisGeNET^43^. Significance of enrichment was evaluated using one-sided Fisher’s exact tests, applied separately to upregulated and downregulated gene sets (Fig. 6A; S28, Supplemental data Table S14). Across these analyses, cells harboring the *DNAJC6* mutation were significantly enriched for neuropsychiatric risk genes, including those associated with ASD (non-syndromic), SCZ, and BP specifically among downregulated genes. Together, these results indicate that familial PD mutations intersect with broad neuropsychiatric risk gene programs, with the strongest and most consistent suppressive effects observed in *DNAJC6*.

**Figure 6:**
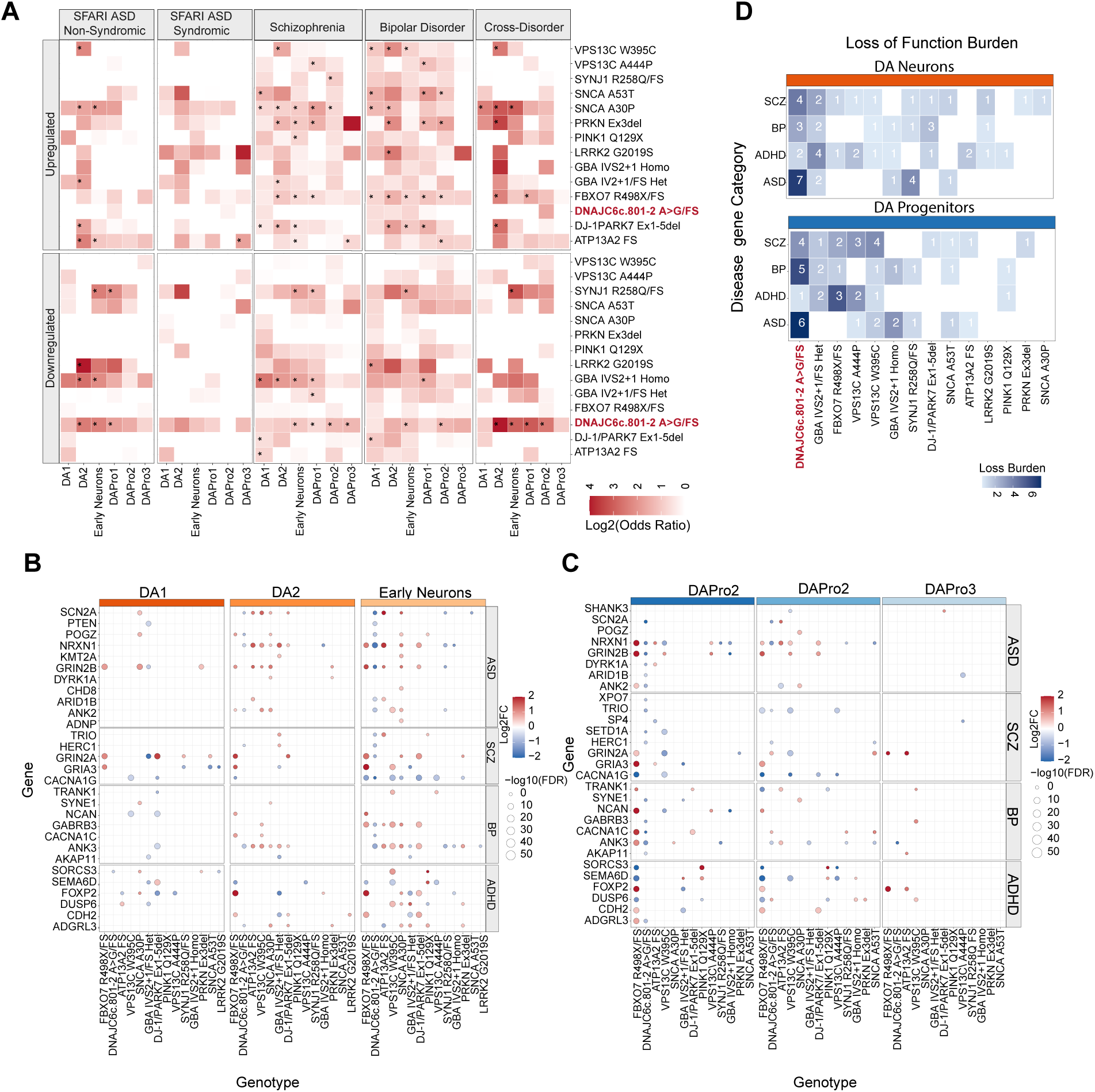
Neurodevelopment and neuropsychiatric risk genes enrichment in DNAJC6 links familial PD mutations to broader brain disorders. **A)** Over-representation of neuropsychiatric risk genes among differentially expressed genes across genotypes and cell type. Heatmap showing Fisher’s exact test odds ratios for enrichment of curated risk gene sets among significantly upregulated and downregulated genes. Each row presents one gene set. SFARI non-syndromic autism spectrum disorder (ASD) genes (n=123), and SFARI syndromic ASD genes (n=121), schizophrenia (SCZ) risk genes (n=497, DisGeNET GDA score >=0.6), and bipolar disorder (BD) risk genes (n=204, DisGeNET GDA score >=0.6), and a 37-gene curated cross-disorder risk gene set spanning ASD, attention deficit/hyperactivity disorder (ADHD), BD, and SCZ. Color indicates log2 odds ratio: red shading indicates over-representation of the gene set among DE genes, with darker red corresponding to stronger enrichment; white indicates no over-representation (log2 OR=<1). Significant enrichment is marked with an asterisk (BH-adjusted p < 0.05). DE genes were defined as those with adjusted p < 0.05 and absolute log2FC of at least 0.5. **B-C)** Dot plots showing differential expression of a curated set of 37 representative high-confidence neuropsychiatric risk genes across all PD mutations, shown for neuronal populations (DA1, DA2, Early Neurons) and progenitors’ populations (DAPro1, DAPro2, DAPro3). Genes are organized by disease category on the y-axis (ASD, SCZ, Bipolar, ADHD). **C)** Heatmap summarizing the cumulative downregulation loss-of-function burden of a curated 37 representative neuropsychiatric risk genes tested across four disease categories: ASD (SFARI category 1), ADHD (Demontis et al., 2023), BD (PGC3/BipEx), and SCZ (SCHEMA Tier 1, Singh et al., 2022). Only a subset of these genes is significant (FDR adjusted p-value<0.05, log2FC cutoff 0.5). Empty tiles indicate no significant downregulation of genes from that specific category in each genotype. Annotated numbers indicate the count of downregulated genes. Genotypes are ordered on the x-axis by decreasing total loss burden across all disease categories and cell populations.

We next curated a set of 37 neuropsychiatric risk genes across four disorders. ASD genes were drawn from SFARI category 1, representing genes with strong de novo or rare variant evidence^42^. SCZ genes were drawn from the SCHEMA rare variant burden analysis (SCHEMA Tier 1^44,45^). BP and attention-deficit/hyperactivity disorder (ADHD) genes were selected from genome wide significant loci in published GWAS studies^46,47,48,49^(Fig. 6B-C). 33 out of 37 genes were dysregulated by atleast one PD mutation across both neuronal and progenitor populations (Fig. 6B-C), indicating overlap between familial PD mutation transcriptional signatures and neurodevelopmental and neuropsychiatric genes. *DNAJC6* mutations showed the largest dysregulation of neuropsychiatric risk genes (22 of 37 genes), with a strong suppressive bias (21 genes downregulated). Some of the most strongly disrupted genes included *SCN2A*, *CACNA1G*, *GRIN2B*, and *NRXN1,* which encode voltage-gated sodium and calcium channels, an NMDA receptor subunit, and a synaptic adhesion molecule^50,51,52,53^. To quantify the magnitude of neuropsychiatric risk gene dysregulation across mutations, we computed a loss-of-function burden score for each genotype by summing the mean (log2FC) of downregulated risk genes from four disorders (ASD, SCZ, BP, ADHD) within DA neurons and progenitor populations (Fig. 6D, Supplemental data Table S14). DNAJC6 showed the highest burden across all four disease modules, driven predominantly by suppression of seven ASD and four SCZ risk genes in DA neurons and a comparable burden in progenitors. GBA heterozygous and FBXO7 followed, reinforcing the genotype-specific nature of neurodevelopmental gene impact. Collectively, this demonstrates disruption of genes controlling synaptic function and neuronal excitability in response to loss of *DNAJC6*. Interestingly, while the *DNAJC6* mutation predominantly caused transcriptional suppression of these genes, *SNCA*-A30P, *ATP13A2*, and *PRKN* mutations induced expression of this gene set. Broader ASD, schizophrenia, and bipolar disorder gene-set loss of burden analyses and gene expression are shown in Supplementary Fig. S28-S29. In summary, these analyses reveal that out of all PD risk genes analyzed, *DNAJC6* mutation causes the largest alterations in risk genes known to cause neuropsychiatric disorders, in particular ASD and schizophrenia. This is consistent with the fact that *DNAJC6* mutations are associated with earlier onset disease than typical familial or sporadic PD and may reflect the known dopamine component associated with ASD^54^ and SCZ ^55,56^.

## Discussion

Here we generated a comprehensive single cell transcriptomic atlas (>200,000 cells) of hPSC-derived dopamine neurons harboring one of 14 familial PD mutations across 11 genes from the iSCORE-PD resource^13^. Our analysis revealed that transcriptional perturbations are strongly cell type dependent, with dopamine neurons (DA1) exhibiting the greatest transcriptional susceptibility across genotypes. By integrating cell type prioritization with pathway and regulatory network analyses, we define transcriptional signatures associated with specific PD mutations and identify molecular programs that are perturbed across the dopamine lineage.

Our work complements and extends previous single-cell studies of PD using hPSC-derived neurons^7,18,57^. Prior studies typically examined one or a small number of mutations, often on different genetic backgrounds, which complicates systematic comparison of mutation effects and can confound genotype-specific transcriptional changes with background genetic variation. The iSCORE-PD platform addresses this limitation by engineering all mutations into a common parental hESC line, enabling direct comparison of transcriptional consequences across mutations in an isogenic context. Correlation with publicly available datasets, including FOUNDIN-PD^18^ and fetal midbrain atlases^16^, confirmed that the differentiated populations capture key molecular features of midbrain dopamine neuron identity and developmental progression. These validations support the biological relevance of the mutation-driven transcriptional signatures observed in this study.

To facilitate community use of this resource, the complete cell type resolved datasets generated here are publicly available, including differential expression gene lists, pathway annotations, and primary sequencing data through GEO and associated repositories. We anticipate that this dataset will serve as a benchmark resource for the PD research community, enabling comparisons across studies and guiding the prioritization of candidate genes and pathways for functional investigation.

The transcriptional signatures identified here point to iron homeostasis and ferroptosis-related pathways as a convergent vulnerability axis across multiple PD mutations. Iron accumulation and lipid peroxidation are well-established features of PD in the SNc. Ferroptosis, an iron-dependent regulated form of cell death driven by phospholipid peroxidation, could plausibly contribute to dopamine neuron loss in PD^58^. In our dataset, the downregulation of *FTL* across multiple genotypes, alongside the suppression of the nuclear receptors *RXRB* and *RXRG*, defines a transcriptional pre-ferroptotic state that precedes overt cytotoxicity. These results are particularly significant given recent evidence that nuclear receptor signaling can suppress ferroptosis defense factors including GPX4, FSP1, PPARalpha, SCD1 and ASCL3, and FXR’s anti-ferroptotic activity involves cooperation with RXR^59^. These findings extend prior observations that lipid peroxidation and iron homeostasis dysregulation represent interacting pathological mechanisms in PD^60^ and highlight the value of cross-comparison approaches for uncovering convergent vulnerability programs.

A fundamental question in PD genetics is whether rare monogenic mutations and common polygenic risk converge on shared molecular mechanisms. By integrating differential expression datasets with PD GWAS-nominated risk genes, we demonstrate that familial PD mutations dysregulate genes implicated in sporadic disease risk. This enrichment of GWAS-nominated genes was strongest in DA1 neurons, consistent with the selective vulnerability of dopamine neurons in PD. Importantly, convergence analysis restricted to mutation pairs processed in independent differentiation batches revealed reproducible overlaps of GWAS associated genes with concordant directionality, providing strong evidence that these signals reflect genuine biological convergence rather than technical artifacts. These findings support a model in which a subset of genetically diverse familial mutations converge on overlapping transcriptional responses, some of which intersect with the polygenic risk architecture of sporadic PD.

*DNAJC6* encodes auxilin, a key regulator of clathrin-mediated synaptic vesicle recycling, and dopamine homeostasis^35^, and its loss impairs neurodevelopmental synaptic programs in iPSC-derived dopamine neurons^61^. A patient-derived iPSC model of *DNAJC6* parkinsonism recently demonstrated neurodevelopmental dysregulation affecting ventral midbrain patterning and neuronal maturation, together with synaptic vesicle recycling defects that were partially rescued by lentiviral gene transfer^61^. Among the PD mutations analyzed, *DNAJC6* produced a distinct transcriptional signature characterized by suppression of several neuropsychiatric disorder risk genes, particularly those associated with ASD and SCZ risk. Notably, those genes encode proteins that are important for synaptic transmission or neuronal excitability. Further research shows that DNAJC6-dependent endocytic defects disrupt WNT-LMX1A signaling during dopamine neuron development^62^. These findings provide a molecular underpinning for the developmental delay and psychiatric manifestations observed in patients with biallelic *DNAJC6* mutations.

Several limitations should be considered when interpreting our results. While the hPSC differentiation system enables scalable and controlled comparison of mutations, it cannot fully capture key *in vivo* features of PD, including aging-related stressors, interactions with other cell types including microglia, and neuronal circuit activity. Consequently, the transcriptional perturbations observed here are generally modest in magnitude and likely reflect early mutation-driven changes in cellular state rather than late neurodegenerative processes. Because these transcriptional effects are often subtle, some mutations did not produce signatures that clearly exceeded the background variance inherent to *in vitro* differentiation-based systems. To minimize confounding effects from differentiation variability, all comparisons were performed against batch-matched wild-type controls differentiated in parallel. In addition, the analyses were conducted during relatively early stages of neuronal maturation and under baseline culture conditions without exogenous stressors, meaning that some transcriptional signatures may represent developmental states rather than disease processes that emerge over decades in patients. Finally, the WIBR3 parental line used throughout the iSCORE-PD collection is female hESC line, and few isogenic comparisons show differential expression of *XIST* and X-linked genes. In primed female hPSCs, prolonged culture leads to loss of *XIST* coating and erosion of X-chromosome inactivation (XCI), resulting in partial reactivation of the inactive X^63,64,65^. This eroded state persists during neuronal differentiation^66,67,68^. Because each iSCORE-PD line was independently edited and clonally expanded, even isogenic lines can acquire distinct XCI states, causing apparent differences in *XIST* and X-linked gene expression that are unrelated to the underlying PD mutation.

Beyond the specific insights highlighted here, the scale and isogenic design of this dataset provide a framework to identify convergent and divergent transcriptional alterations across PD genotypes. Our analyses point to one such axis, with convergent transcriptional signatures implicating ferroptosis-related pathways as a potential shared vulnerability across multiple PD mutations. More broadly, the convergence of familial mutations on GWAS-linked sporadic risk genes, together with mutation-specific programs such as the neurodevelopmental axis associated with *DNAJC6*, illustrates how large-scale isogenic stem cell modeling can reveal shared and genotype-specific mechanisms underlying dopamine neuron vulnerability in PD.

## Resource Availability

### Lead contact

Further information and requests for resources and reagents should be directed to and will be fulfilled by the lead contact, Dr. Dirk Hockemeyer.

### Materials availability

Isogenic hESC lines used in this study were generated and characterized as part of the iSCORE-PD collection^13^and are available from the lead contact upon request and from WiCell repository.

### Data and code availability

- All single-cell-sequencing data generated in this study have been deposited at the Gene Expression Omnibus (GEO, GSE329807) and will be publicly available as of the date of publication. (pending)
- Codes for the single-cell RNA-seq analysis will be available the GitHub repository. (pending)

## ACKNOWLEDGMENT

we thank all members of the Bateup, Hockemeyer, Soldner, Rio laboratories for helpful discussions and comments on the manuscript. This work was funded by Aligning Science Across Parkinson’s (ASAP-024409, ASAP-000486, MJFF-027664) through the Micheal J. Fox Foundation for Parkinson’s Research (MJFF) to the Bateup, Hockemeyer, Soldner, Rio laboratories. This work utilized the computational resources of the Savio high-performance computing cluster at the University of California, Berkeley. Single-cell library preparation and sequencing were performed at the QB3 Genomics core facility, University of California, Berkeley. For the purpose of open access, the author has applied a CC-BY public copyright license to the Author Accepted Manuscript version arising from this submission.

## AUTHOR CONTRIBUTIONS

KMS, DCR, FS, DH, HB designed the study and experiments. KMS and JD generated dopamine neurons and performed single-cell RNA sequencing experiments. KMS conducted the computational analysis. SP generated the GBA single-cell data. KA and AS conducted immunostaining and quantification experiments. KMS, DH and HB wrote the paper with inputs from all the authors.

## DECLARATION OF GENERATIVE AI AND AI-ASSISTED TECHNOLOGIES

The authors used AI assistance (Ex: ChatGPT) to streamline the analysis code used in paper.

## METHODS

### KEY RESOURCES TABLE

**Table S1.**
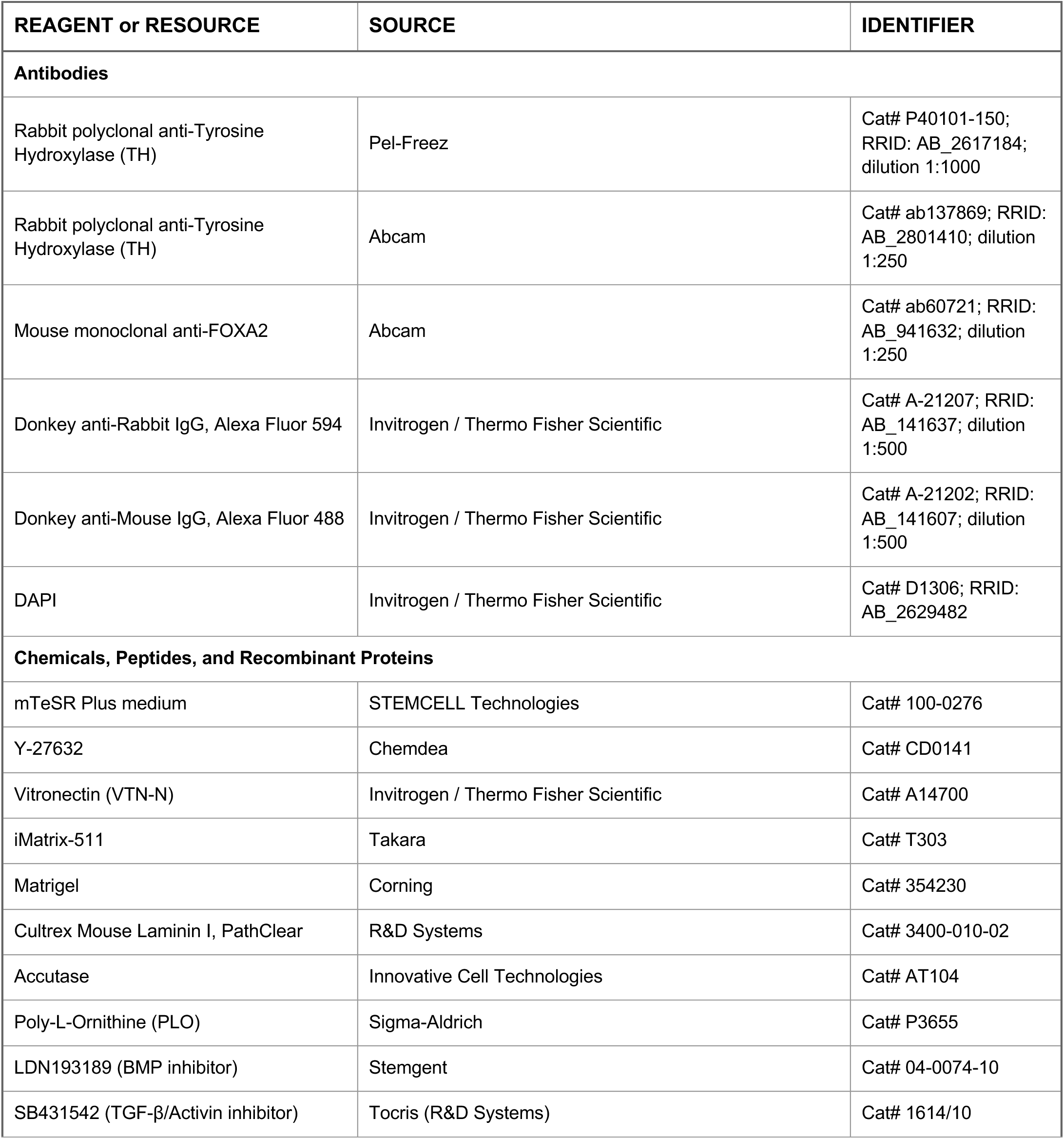
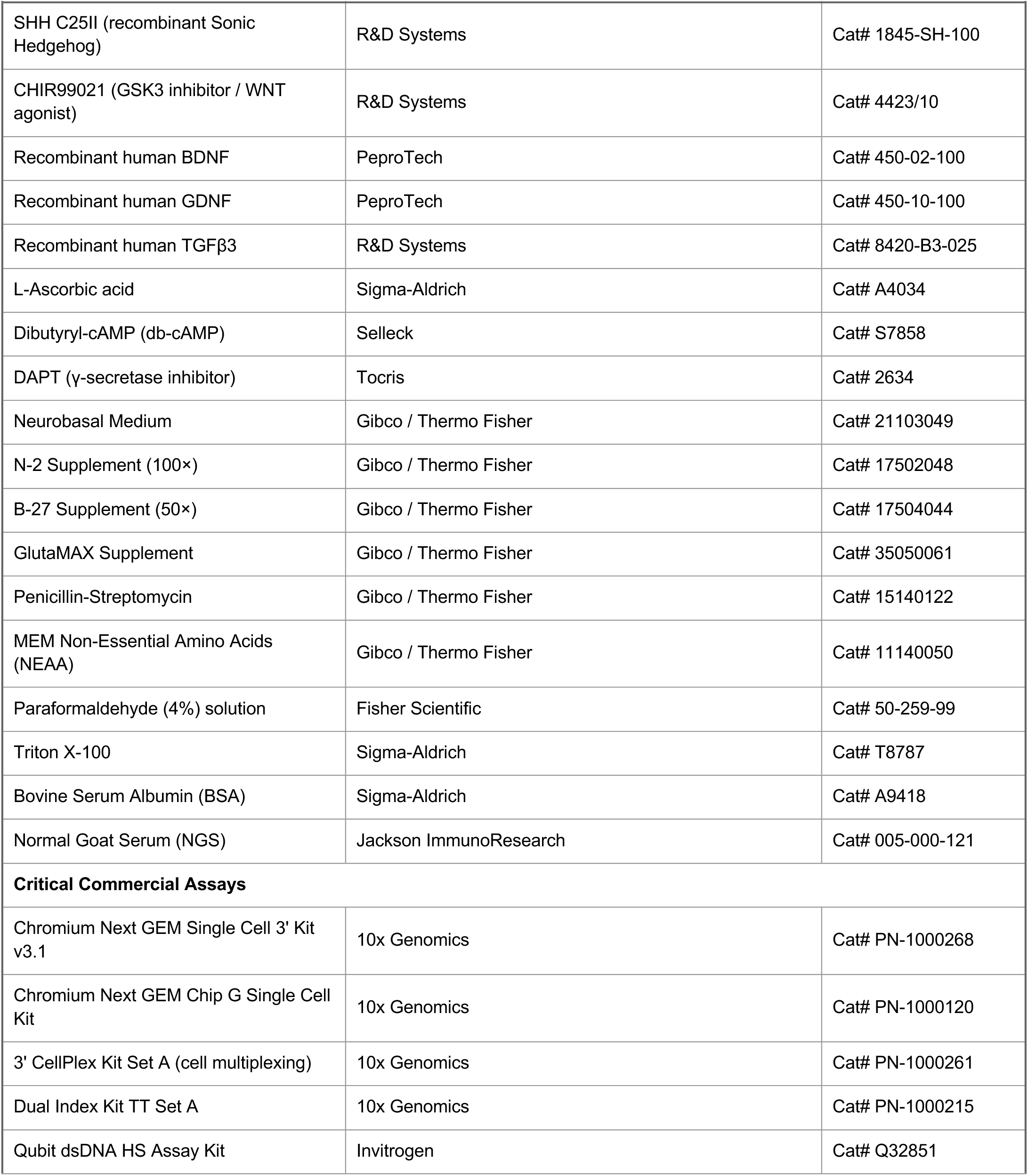
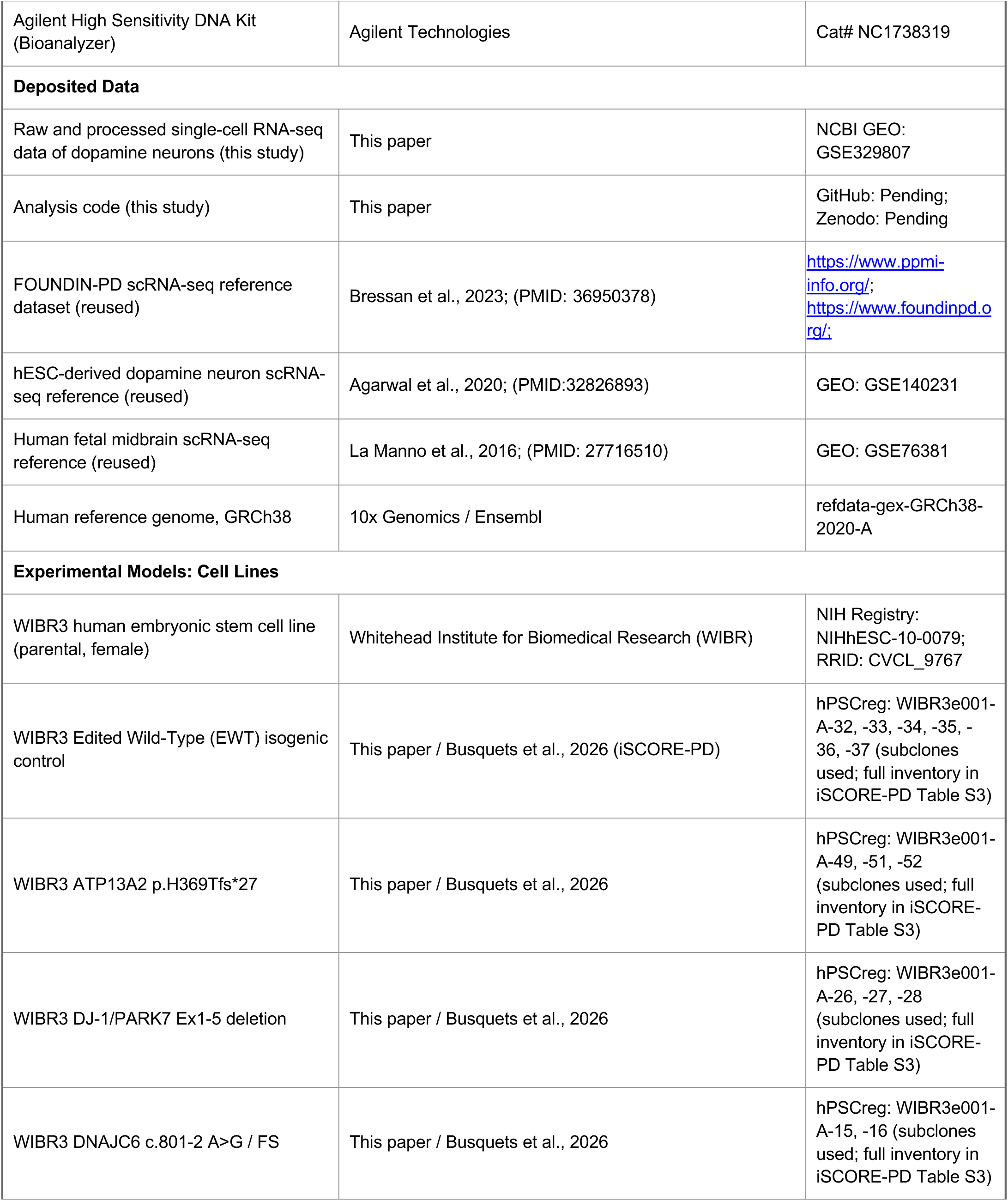
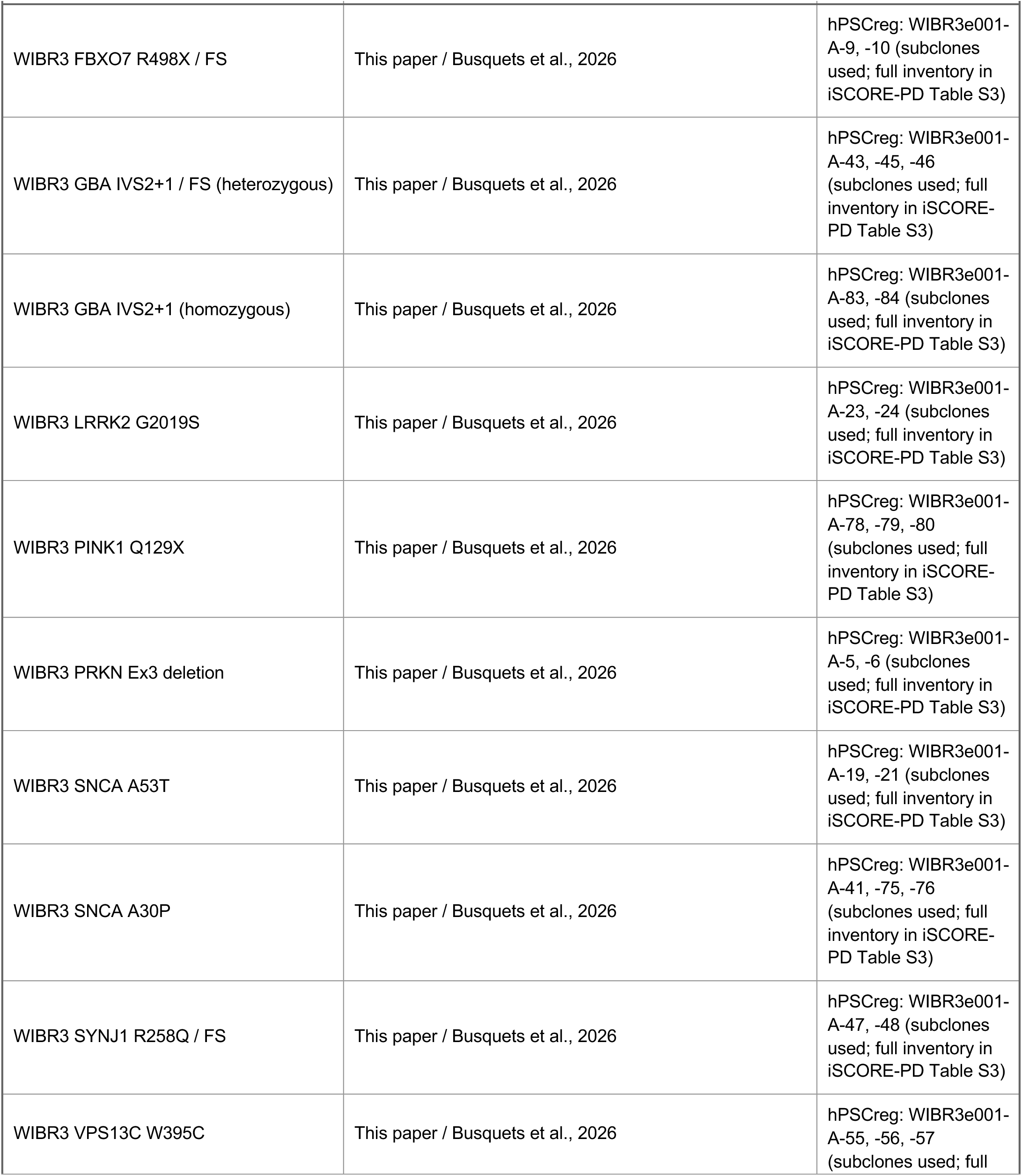
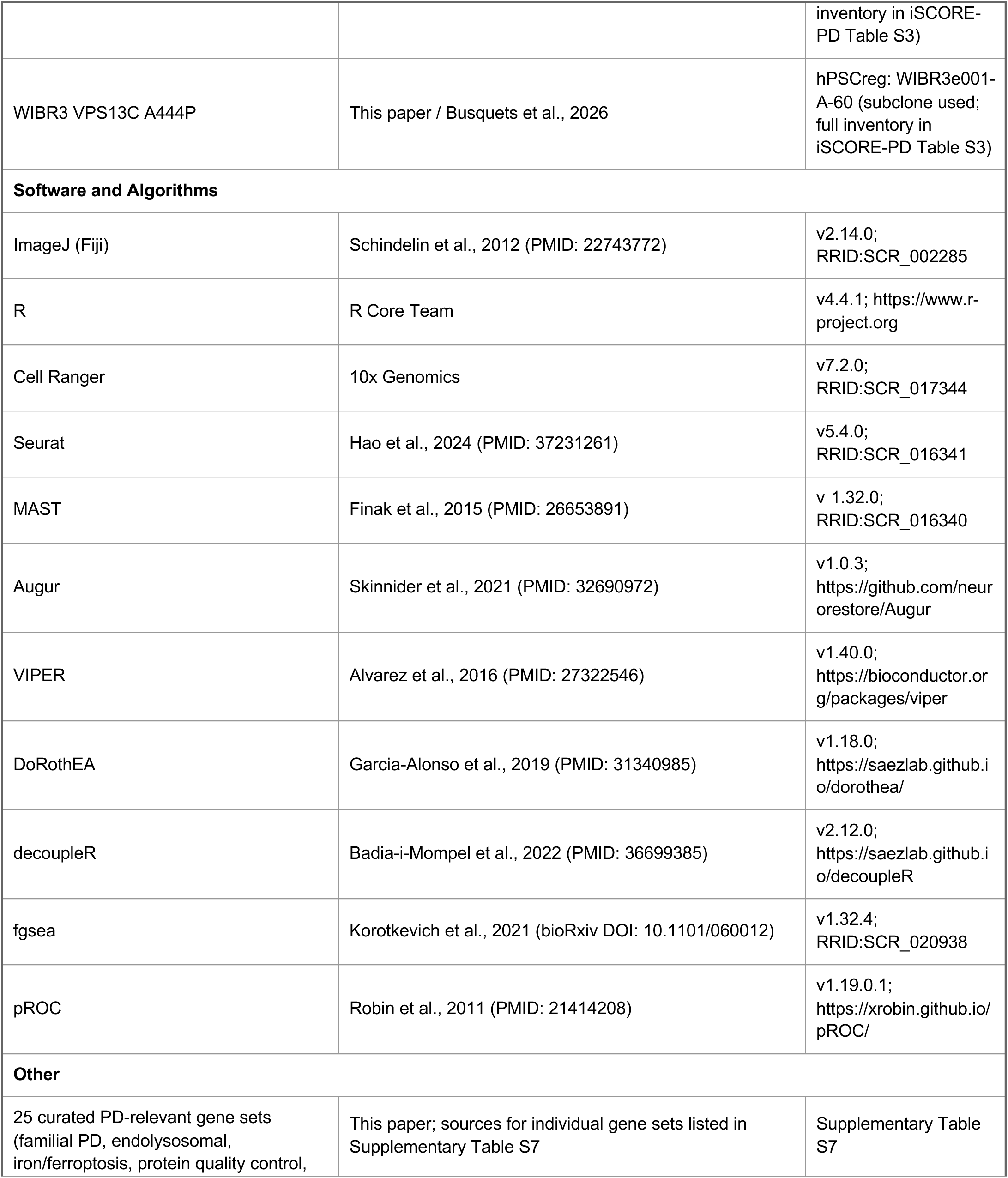
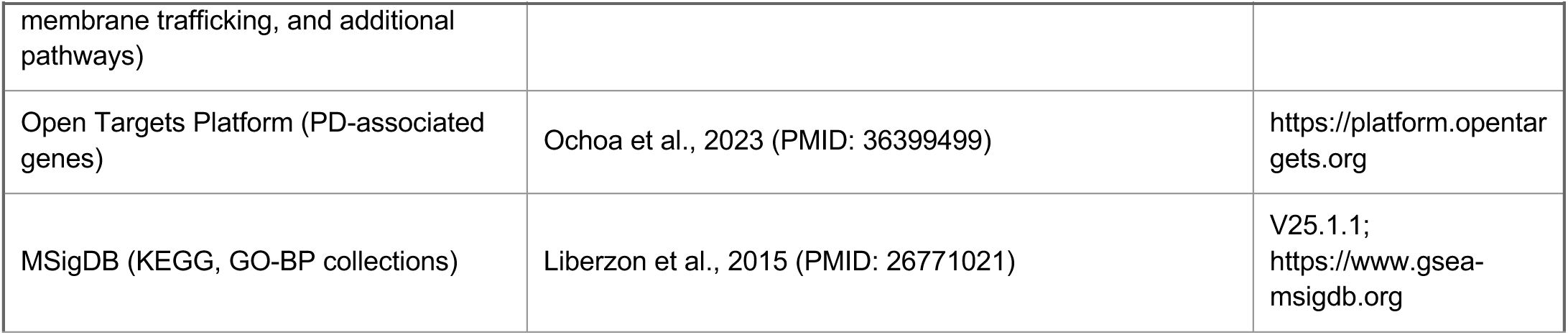
Clinical summary of patients.

### EXPERIMENTAL MODEL AND STUDY PARTICIPANT DETAILS

#### Human pluripotent stem cell lines

All experiments were performed using the WIBR3 hESC line (female; NIH approval number NIH hESC-10-0079) and WIBR3 CRISPR engineered familial PD lines from the iSCORE-PD collection^13^WIBR3 cells were maintained on feeder-free conditions on vitronectin or laminin-coated plates in mTESRplus medium (STEMCELL Technologies) at 37°c with 5% CO_2_, and 3% O_2_. Cells were passaged using accutase at approximately 70-80% confluence. All cell lines were regularly tested for mycoplasma contamination and confirmed negative and karyotype analysis was performed on parental and edited lines to confirm genomic integrity. The complete quality control framework, including whole-genome sequencing of each line, has been described previously^13^. Fourteen PD-associated mutations across 11 genes were introduced into the WIBR3 parental line using TALEN/CRISPR/Cas9/prime editing. *ATP13A2* p.H369Tfs*27, *DJ-1/PARK7* Ex1-5del, *DNAJC6*c.801-2 A>G/FS, *FBXO7* R498X/FS, *GBA* IVS2+1/FS (Heterozygous), *GBA* IVS2+1 (Homozygous), *LRRK2* G2019S, *PINK1* Q129X/FS, *PRKN* Ex3del, *SNCA* A53T, *SNCA* A30P, *SYNJ1* R258X/FS, *VPS13C* W395C, *VPS13C* A444P. Detailed protocols are described previously in Busquets O et al., 2026.

#### Midbrain dopamine neuron differentiation

Midbrain dopamine (DA) neurons were differentiated from human pluripotent stem cells using a modified floor-plate protocol^14^. Briefly, neural induction was initiated with dual SMAD inhibition using LDN193189 (BMP inhibitor) and SB431542 (TGF-B/Activin inhibitor) in N2B27 base medium (days 0-11). Ventral midbrain floor plate identity was specified by addition of the Sonic hedgehog (SHH) and WNT activator CHIR99021. DA neuron specification and maturation were promoted from day 11 onwards using neurotrophic and trophic factors including BDNF, GDNF, TGFb3, ascorbic acid, and dibutyryl-cAMP. Cells were replated at day 16 and day 25 at high density onto ploy-L-ornithine (PLO) and laminin-coated plates and maintained in maturation medium with media changes every two days. Cells were harvested at day 35-45 for single cell RNA sequencing and differentiation quality was confirmed by immunofluorescence for TH, FOXA2 (Supplementary Fig. S4). Differentiation experiments were organized into seven batches (B1-B7), each containing 2-4 mutant genotypes alongside isogenic Edited WT (EWT) controls processed in parallel. The genotype batch information tabled in Table S1. For specific details of the protocols, consult the published materials on protocols.io: https://doi.org/10.17504/protocols.io.3byI4q8yovo5/v1.

#### Immunofluorescence

Cells were fixed in 4% paraformaldehyde for 15 minutes at room temperature (RT), permeabilized with 0.3 % Triton X-100, and blocked in 3% BSA or 10% NGS for 1 hr at RT. Primary antibodies (see Key resource table) were incubated overnight (O/N) at 4C. Secondary antibodies conjugated to Alexa flour 488 and 594 were used for 1 hr at RT. Nuclei were counterstained with DAPI. Images were acquired on a fluorescence microscope. Details on the protocol used can be found in protocols.io: https://doi.org/10.7504/protocols.io.yxmvm3146I3p/v1.

#### Single-cell RNA sequencing and 10X genomics cell-plex multiplexing

Differentiated dopamine neurons (day35-45) were dissociated into single-cell suspensions using accutase for 15-30 min at 37°c, filtered through 40-µm cell strainers and assessed for viability Only samples with >80% viability was processed. Samples were labeled with 10X genomics 3’ CellPlex oligos (CMOs) for multiplexing, according to the 10x recommendations protocol and pooled prior to loading pooled samples were loaded onto a 10x Genomics Chromium controller targeting 30,000 cells per lane using Chromium Single Cell 3’ Gene Expression v3.1 chemistry. Gene expression and CellPlex libraries were prepared according to the manufacturer’s protocol and sequenced on Illumina NextSeq, NovaSeq 6000, or NovaSeq X platforms targeting 35,000-50,000 reads per cell. Sequencing was performed at the QB3 Genomics core facility, University of California, Berkeley.

#### Single-cell data preprocessing and quality control

Raw sequencing reads were aligned to the GRCh38 human reference genome and gene expression matrices were generated using Cell Ranger v7.2, 10X genomics. Quality control filtering was performed using Seurat v5 in R version 4.4. Cells were retained if they met the following criteria, nCount > 1000, nFeature >200, <15% mitochondrial reads. After the quality control filtering 209,000 cells were retained across all genotypes and batches.

#### Normalization, Integration, and clustering

Gene expression data were processed in Seurat. Counts were log-normalized using Seurat’s *NormalizeData()*, with a scale factor of 10,000 and the top 2000 variable features per sample were identified using *FindVariableFeatures()* with variance stabilizing transformation. Cells were filtered based on the quality control metrics described above, and S-phase and G2/M-phase scores were computed for each cell using the standard Seurat cell-cycle gene lists. Each sample was also normalized using SCTranform (Hafemeister and Satija, 2019). SCT-normalized layers were used only for sample integration, whereas downstream analyses including DE testing were performed on the log-normalized RNA assay. For integration, 3000 features were selected with *SelectIntegrationFeatures()* and prepared using *PrepSCTIntegration().* PCA was performed on the merged SCT assay. Batch integration was performed using canonical correlation analysis through Seurat v5 *IntegrateLayers()* with SCT normalization, generating a CCA-integrated reduction. Shared nearest graph construction, Louvain clustering and UMAP visualization were performed on the integrated reduction using the first 30 dimensions. Clusters were identified using at resolution 0.5.

#### Cell-type annotation

Cell-types were annotated based on expression of canonical marker genes drawn from published midbrain dopaminergic lineage references^16,69,70,10,71^. Seven cell populations were identified: DA1, DA2, Early Neurons, DAPro1, DAPro2, DAPro3 and ProProgen. Marker genes, supporting references, and the rationale for each annotation are provided in supplemental data Table S2.

#### Differential expression analysis

Differential expression (DE) testing was performed for each mutation against batch matched EWT controls within each cell-type using MAST framework^23^ as implemented in Seurat’s *FindMarkers()* function. We Paired mutants with EWT controls differentiated alongside them controlled for biological variation arising from differentiation batch and performed DE analysis. For LRRK2 G2019S, which is processed in both B2 and B3, DE analysis was performed separately in each batch and results were merged by retaining only genes showing concordant direction in both batches. Genes were considered DE at FDR<0.05 and log2FC cut off at 0.25. A more stringent threshold of log2FC cutoff of 0.5 is used in figure 6 enrichment analysis.

#### Cell-type prioritization using Augur

Cell-type vulnerability to genetic perturbation was quantified using Augur^20^, a machine learning framework that trains random forest classifiers on multivariate gene expression to distinguish mutant cells from control within each cell-type. For each of the 14 mutations, Augur was applied independently to compare mutant against batch matched EWT cells, yielding per cell-type specific area under the receiver operating characteristic curve (AUC) score. AUC values approaching 1.0 indicate strong transcriptional perturbation (high classifier separability), while values near 0.5 indicate minimal change. This AUC is conceptually distinct from single-gene and module-score AUC metrics described below in that it summarizes multivariable transcriptional separability rather than the discriminative power of a single feature. Augur was run with default parameters (subsample size=20 cells per cell type per condition). Results were visualized as per-genotype bar plots ranked by AUC scores and as UMAP overlays in which cells were colored by their cell-type level AUC for the corresponding mutation.

#### Directional single-gene AUC-ROC classification

To identify individual genes whose expression distinguishes mutant from control cells, a single-gene AUC-ROC classification framework was applied. For each gene within a curated gene set, the normalized expression value was used as a univariable classifier of mutation versus control identity. AUC was computed for each gene-genotype-cell-type combination using the *pROC::roc()*^72^ in R with direction =”<”. Under this convention AUC > 0.5 indicates upregulation in the mutant (higher expression predicts mutant status), AUC <0.5 indicates downregulation (inverse classifier), and AUC=0.5 indicates no discriminative power. This framework was applied to 353 genes across 25 curated PD relevant gene sets spanning familial PD genes, Endolysosomal degradation, iron ferroptosis, protein quality control, membrane trafficking, and additional pathways (Supplementary Table S7). For each gene, directional consistency was defined as the fraction of genotype-cell type comparisons in which the gene was dysregulated in the same direction across mutations, with upregulated (AUC>0.5) and downregulated (AUC<0.5) calls treated as opposing directions and AUC=0.5 ties excluded as neutral. Genes with negligible effect sizes (SD of AUC <0.01 and mean distance from 0.5<0.02) were removed before ranking.

#### Pathway module scoring and classification

Pathway-level perturbation was assessed using Seurat’s *AddModuleScore()*^31,73,74^, which computes a composite expression score for each gene set as the average expression of its member genes minus the mean expression of background control genes matched in aggregate expression level. Module scores were computed for each of the 25 curated gene sets. For each genotype-cell type combination, AUC-ROC analysis was applied to the module scores to test whether each gene set distinguished mutant from batch-matched EWT cells, the AUC was signed to reflect the direction of the mean difference between mutant and EWT distributions, such that AUC >0.5 indicates pathway enrichment (upregulation) in the mutant and AUC <0.5 indicates pathway depletion (downregulation).Statistical significance was assessed by Wilcoxon rank-sum test on the score distributions, with Benjamini-Hochberg correction across all gene set genotype-cell type comparisons. . This pathway-level AUC provides a complementary measure to single-gene analysis, capturing coordinated pathway-level perturbation that may be missed by individual gene tests. Correlation between Augur-derived cell-type perturbation scores and pathway module AUC scores was computed across the 14 genotypes using Pearson correlation. This correlation tests whether mutations causing greater overall transcriptional perturbation (high Augur AUC) show systematically higher or lower pathway module scores, identifying biological programs that scale with perturbation magnitude.

#### Transcription Factor activity inference and analysis

Transcription factor (TF) activity was inferred at single-cell resolution using the VIPER^22^ with TF-target gene regulons from the DoRothEA database^21^. Regulons at confidence levels A, B, and C were included, encompassing 298 TFs with sufficient target gene representation. VIPER computes a TF activity normalized enrichment score (NES) for each cell by assessing whether the TF’s known target genes are coordinately upregulated or downregulated relative to other genes, thereby inferring TF activity from target gene expression patterns. NES values were computed on the log-normalized RNA expression matrix and stored as a separate TF activity assay in the Seurat object for downstream analysis. Variance in TF activity was quantified using one-way ANOVA on per-cell NES values. Effect sizes were calculated as eta-squared (n^2^=SS_between/SS total) for one-way ANOVA models testing cell-type effects within each genotype and genotype effects within each cell type. Per-TF n^2^ distributions for each mutant were compared with EWT using one-sided Wilcoxon rank-sum test. Differential TF activity was tested within each cell type by comparing each mutant with its batch-matched EWT control using a Wilcoxon rank-sum test on per-cell NES values, implemented with *FindMarkers()* on the TF activity assay. Significant TF activity changes were defined as BH-adjusted p<0.05 and absolute log2FC of atleast 0.5. To identify TFs with recurrent activity changes across DA neuron populations, significant TFs were classified as showing increased or decreased activity in DA1, DA2, and Early Neurons. Within each cell type, a TF was considered directionally consistent if it had atleast four significant genotype-level hits and more than 50% of those hits were in the same direction, increased or decreased activity. TFs meeting this criterion in atleast two DA neuron cell types and accumulating atleast 10 significant hits across DA1, DA2 and Early Neurons were ranked using the composite score:

Rank score = (n cell types +1) * mean consistency* mean effect size* total hits +1) Where mean consistently and mean effect size were averaged across all three DA neuron cell types. The top 25 ranked TFs were selected as the core recurrently dysregulated TF activity set. Differential TF activity results were reported in supplementary Fig. S7 and S10.

#### PD GWAS gene enrichment analysis

PD associated genes were obtained from the Open Targets Platform^33^ (MONDO_0005180), filtered for GWAS credible-set evidence and an overall association score >0.1yielding 267 GWAS implicated genes. For each genotype-cell type combination, one-sided Fisher’s exact tests were used to assess whether DE genes (FDR adjusted p-value < 0.5, log2FC cutoff 0.25) were enriched for GWAS genes relative to the universe of all genes tested (detected) in that comparison. This condition-specific universe ensures that enrichment is not inflated by genes not expressed in that cell-type. Cell-type level enrichment was summarized as the median odds ratio across genotypes and tested against null (OR=1) using a one-sided Wilcoxon signed-rank test. To assess lineage-level enrichment, Pan-DA (DA1, DA2, and Early Neurons combined), and Pan-DAPro (DAPro1, DAPro2, DAPro3 combined) gene sets were generated by taking the union of DE genes and detected genes across constituent cell types, followed by the Fisher’s exact test. P values were corrected using the Benjamini-Hochberg method within each analysis.

#### Convergence of PD GWAS-overlapping DE gene analysis

For each genotype and cell type, PD GWAS-overlapping DE genes were defined as significantly DE genes, (log2FC 0.25 and FDR <0.05) that interested a PD GWAS gene panel from Open Targets Platform, globalScore >0.1, n=267. Convergence was assessed both across DA neurons collectively DA1, DA2, and Early Neurons, and within each DA neuronal cell type separately. For the pan-DA analysis, genotype-level gene sets were generated by taking the union of PD GWAS-overlapping DE genes across DA1, DA2, and Early Neurons; genes significant in multiple cell types were retained only when the consistently up or down regulated. For each genotype pair, convergence was quantified using the Jaccard index, overlap significance by one-sided hypergeometric test using the 267 PD GWAS genes as the background, directional concordance. Directional concordance is defined as the fraction of overlapping genes that were upregulated or downregulated in both genotypes. Pairs with fewer than two overlapping genes were excluded. Hypergeometric P values were Benjamini-Hochberg corrected within each analysis, pan-DA or individual cell type, and genotype pairs were classified as convergent when FDR <0.05 and directional concordance atleast 60%. To minimize the batch driven convergence, only cross-batch genotype pairs were retained. For LRRK2 G2019S, profiled in B2 and B3, batch-specific DE results were merged before convergence analysis; genes significant in both batches were averaged when they were consistently upregulated or down regulated and excluded when discordant, while genes significant in only one batch are retained. For batch filtering, LRRK2 G2019S was treated as matched to both B2 and B3, and comparisons to genotypes from either batch were excluded. Pan-DA and cell type-specific convergence results are reported in supplemental figure S26. For each cross-batch mutation pair, we computed the number of overlapping genes, then Jaccard index (intersection/union), followed by directional concordance percentage of overlapping genes showing the same direction of change.

#### MJFF Targets to Therapies (T2T) gene set analysis

High-priority PD target genes were defined from the MJFF Targets to Therapies (T2T) initiative by retaining the 59 genes annotated as “Selected as a top 59 Target”. These genes were intersected with DE genes identified for each genotype by cell-type comparison. Directional concordance was assessed using the T2T “PD differential expression direction” annotation, excluding targets annotated as inconsistent or not detected. Observed direction was defined by the log2FC (up or down), and concordance was calculated as the percentage of DE direction-annotated T2T genes matching the expected PD associated dysregulation.

#### Gene set enrichment and Over representation analysis

Gene set enrichment analysis (GSEA) was performed using the fgsea^75^ with minSize=15 and maxSize=500 in R. Genes were ranked by signed significance metric: Rank= -log10(padj+1x10^-10^) x sign (log2FC) computed from MAST differential expression output. For SNCA A53T and GBA homozygous, where p-value based ranking yielded sparse results, log2FC was utilized as the ranking metric. Enrichment was tested against two MSigDB v2023.2.Hs collections: Gene Ontology BP and KEGG Legacy. To refine for neuronal model a curated lists of terms were excluded prior to enrichment. For the LRRK2 G2019S genotype, results from two independent batches (B2, B3) were integrated by averaging ranking metrics per gene, thereby prioritizing concordant effects. Pathways with FDR <0.05 were considered significant, with Normalized Enrichment Scores (NES) used to compared enrichment magnitude across genotypes.

#### Over-representation analysis (ORA)

Over-representation analysis was performed using *enrichGO* from clusterProfiler^76^(v4.14.6) against the Gene Ontology Biological Process (GO-BP) ontology. For genotype and cell type, ORA was conducted separately on significantly upregulated and downregulated genes (FDR <0.05; log2FC 0.25 threshold). The background universe for each test was strictly defined as the total set of genes tested in the corresponding differential expression analysis. To maintain consistency with GSEA, the curated set of terms were excluded. For the genotype LRRK2 G2019S, input gene sets were restricted to those exhibiting concordant upregulated or downregulated significance across both independent batches (B2 and B3), the background for this comparison was defined as the intersection of all genes detected in both batches.

#### Neurodevelopmental and psychiatric disorder gene enrichment analysis

Overlap between PD mutation DE genes and neurodevelopment/psychiatric disorder risk genes was tested using curated gene sets from published resources and consortia. SFRAI Gene category 1 ASD gene sets (SFRAI database), SCZ and BP gene sets (DisGENET) and a curated cross disorder panel assembled from high relevant genes reported by ASD, schizophrenia, bipolar disorder, and ADHD studies.

#### Neuropsychiatric gene loss-burden analysis

Across PD genotypes, loss burden was calculated for neuropsychiatric gene sets, including ASD SFARI category 1, DisGeNET SCZ and BP and curated cross-disorder gene sets. Significantly DE genes (FDR <0.05, log2FC cutoff 0.5) used for scoring. Scores were calculated separately in neuronal populations (DA1, DA2, and Early Neurons) and Progenitors (DAPro1, DAPro2, DAPro3). Within each genotype and cell-population group, genes detected in multiple cell types were collapsed to a single value, defined as the mean absolute log2FC across downregulated cell types. Loss was calculated as the sum of these values for each genotype and disease gene set. For LRRK2/G2019S, genes were retained only when downregulated in both batches.

#### Quantification and statistical analysis

Statistical analyses were performed in R version 4.4. Sample sizes (number of cells, genotypes, batches) and the statistical test applied are reported in the figure legends and corresponding methods subsections. Statistical methods used included the MAST hurdle model with likelihood ratio testing (for single-cell differential expression), adaptive multilevel permutation testing (for gene set enrichment analysis), the Wilcoxon rank-sum test (for differential transcription factor activity, applied per cell), the one-sided Fisher’s exact test (for gene-set over representation and PD GWAS gene enrichment), the one-sided hypergeometric test (for cross-batch pairwise convergence), Pearson correlation (for cross-genotype scaling between Augur perturbation and pathway module AUC scores), and one-way ANOVA with n^2^ effect-size quantification (for variance decomposition of TF activity). All AUC-based metrics (Augur cell-type perturbation, single-gene classification, and pathway module classification) were derived from receiver operating characteristic analyses, with AUC>0.5 indicating depletion. Multiple testing was corrected using the Benjamini-Hochberg method within each defined analysis scope, and adjusted p-values <0.05 were considered statistically significant unless otherwise indicated. All sample sizes, statistical methods, and statistical parameters, including p-values, are detailed in the respective figure legends. Due to the nature of the study, experiments were not randomized and investigators were not blinded during data acquisition or analysis. No statistical methods were used to predetermine sample size, cell numbers per genotype-cell type combination reflect cell yields after quality control filtering of the scRNA-seq data.

## Supplemental Data Tables

**Table S1.** Dataset composition, subclone metadata, and per genotype quality control metrics. Related to Figure 1 and STAR Methods.

Per-genotype summary of subclones, mutations, differentiation batches, GEX and paired CMO library identifiers (including GEO submission naming), total post-quality control cell counts per-cell-type cell yields, and quality control metrics for the iSCORE-PD scRNA-seq atlas.

**Table S2.** Cell cluster annotations and supporting marker genes. Related to Figure 1. Annotation rationale, top marker genes, and supporting literature references for seven cell populations identified in the integrated iSCORE-PD scRNA-seq atlas.

**Table S3.** Differential transcription factor activity across familial PD mutations. Related to Figure 2 and Figure S7 and S23.

DoRothEA/VIPER-inferred TF activity comparing each mutant genotype to its batch-matched EWT across cell types.

**Table S4.** Transcription factor activity variance decomposition. Related to Figure 2.

Per-TF percent variance by cell-type identity (within each genotype) and by genotype (within each cell type) from one-way ANOVA analysis, with Wilcoxon rank-sum p-values comparing each mutant to EWT.

**Table S5.** Differential expression results across familial PD mutations. Related to Figure 3 and Figures S8-S20.

MAST results per cell type for each genotype vs batch-match EWT, one sheet per genotype.

**Table S6.** Gene Set Enrichment Analysis results across familial PD mutations. Related to Figure 3 and Figures S8-S20.

Significant KEGG and GO Biological Process pathways (BH-adjusted p < 0.05) per genotype x cell type.

**Table S7.** Curated PD-relevant gene modules used for module scoring and pathway-level scoring analyses. Related to Figure 4 and Figures S21, S24.

The 25 PD-relevant gene sets (353 genes) grouped by functional category, with member genes and supporting literature references for each module.

**Table S8.** Gene-level AUC-ROC analysis of consistently dysregulated genes across familial PD mutations. Related to Figure 4 and Figure 22.

Per-gene directional AUC values across 14 mutations x 3 neuronal cell types from 25 curated PD-relevant gene modules.

**Table S9.** Transcription factor consistency ranking across DA neuron populations. Related to Figure 2 and Figure S23.

Per-TF consistency rankings across DA1, DA2 and Early Neurons for TFs qualifying as recurrently dysregulated.

**Table S10.** Pathway module score activity across familial PD mutations. Related to Figure 4. Per-genotype x cell-type module score statistics for the 25 curated PD-relevant gene modules.

**Table S11.** PD GWAS gene list from the Open Targets Platform. Related to Figure 5.

The PD-associated genes (MONDO_0005180) with overall association score >0.1 and GWAS credible-set evidence score per gene.

**Table S12.** PD-GWAS gene enrichment analysis results against differentially expressed genes across mutations. Related to Figure 5.

**Table S13.** PD GWAS genes overlap and concordance results. Related to Figures S26-S27. Pairwise cross-batch convergence of PD genotypes with PD GWAS overlap statistics.

**Table S14.** Neuropsychiatric gene enrichment analysis among differentially expressed genes. Related to Figure 6 and Figure S28-S29.

## Supplementary Materials

**Figure S1:**
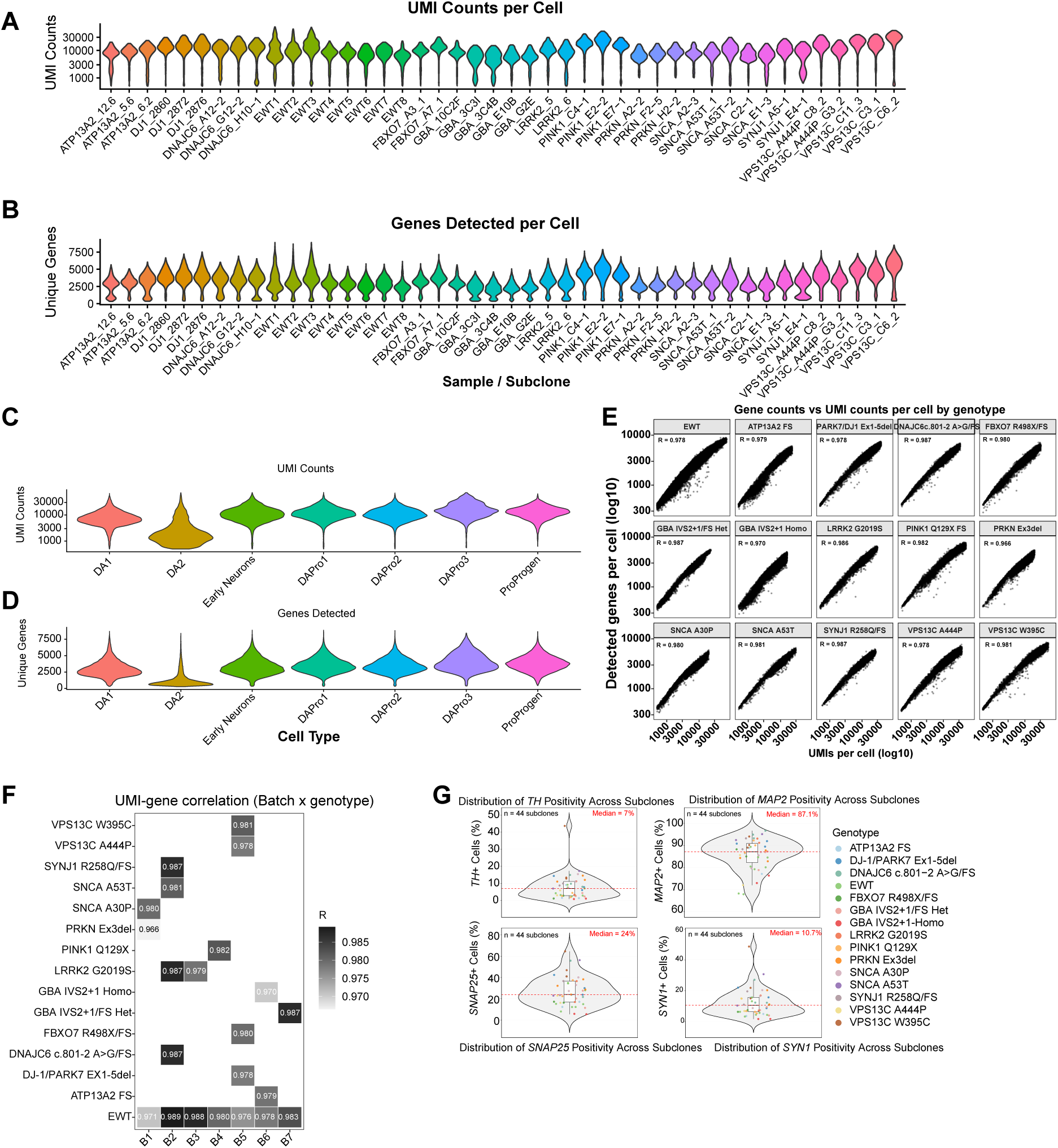
Per-subclone sequencing depth and gene detection quality metrics across iSCORE-PD subclones. A-B) Violin plots displaying single-cell quality metrics for each individual subclone in the iSCORE-PD sc-Atlas after quality filtering. **A)** Total UMI counts per cell displayed on a log10 scale (y-axis, range ∼1000-30,000). Each violin represents the density estimate of all cells within a given subclone. The majority of subclones display compact, symmetric diamond-shaped distributions with the bulk of cells concentrated between ∼3000-10,000 UMIs, indicating consistent sequencing depth across the dataset. **B)** Number of unique genes detected per cell. Most subclones show distributions centered between ∼2000-4000 genes per cell. Subclones are labeled by their mutation identity and clone ID (example: ATP13A2_12.6, SNCA_A53T_1), spanning all 14 familial PD mutations and isogenic engineered wildtype controls (EWT1-EWT8). The broad consistency of violin shape, width, and median position across all subclones confirms uniform library preparation and sequencing depth. **C-D)** Violin plots displaying single-cell quality metrics stratified by annotated cell type across the full integrated dataset. **C)** Total UMI counts per cell displayed on a log10 scale (y-axis, ∼1000-30,000). **D)** Number of unique genes detected per cell. Seven annotated populations are shown: dopamine neuron sub-populations DA1, DA2, Early Neurons, and dopamine progenitor populations DAPro1, DAPro2, DAPro3 and proliferating cell type ProProgen. DA2 neurons are a notable exception, displaying a lower transcriptional complexity compared to all other populations, with a median of ∼1000 detected genes per cell and bottom concentrated violin shape. Importantly, DA2 cells passed all quality control threshold, and this distribution is reproducible across all subclones and batches, ruling out a technical or sample-specific origin. The reduced gene detection in DA2 likely reflects a genuine biological property of this population, potentially representing a transcriptionally quiescent or transitional dopamine state. **E)** UMI-gene count correlation confirms uniform library complexity across genotypes and batches. Scatter plots showing the relationship between total UMI counts per cell (x-axis) and the number of detected genes per cell (y-axis) for each genotype. Each panel represents one genotype, the isogenic engineered wild type control (EWT) and all 14 familial PD mutations. Pearson correlation coefficients (R) are in the top-right corner of each panel. R values range from 0.926 to 0.961 across all genotypes, indicating a consistently strong linear relationship between sequencing depth and transcriptome complexity in all lines. **F)** Heatmap of Pearson R values for the UMI-gene correlations for each batch genotype combination. EWT controls are present across all seven batches (B1-B7). Each familial PD mutation was processed in a single designated batch, reflecting the experimental design of the iSCORE-PD platform. The uniform high R values across all batch/genotype cells confirm that the library is consistent regardless of genotype or batch, and that no specific batch or mutation introduces systematic sequencing artifacts. **G)** Single-cell transcriptomic quantification of neuronal and dopamine marker expression across iSCORE-PD subclones. Violin plots showing the percentage of marker-positive cells per subclone across all 44 subclones (n=44). Each dot represents one subclone, colored by genotype. Embedded boxplots show median and interquartile; the dashed line indicates the median across all subclones. Four markers shown: TH (tyrosine hydroxylase; median = 7%), MAP2 (microtubule-associated protein 2; median =87.1%), SNAP25 (synaptosomal-associated protein 25; median=24%) SYN1 (synapsin-1; median =10.7%).

**Figure S2:**
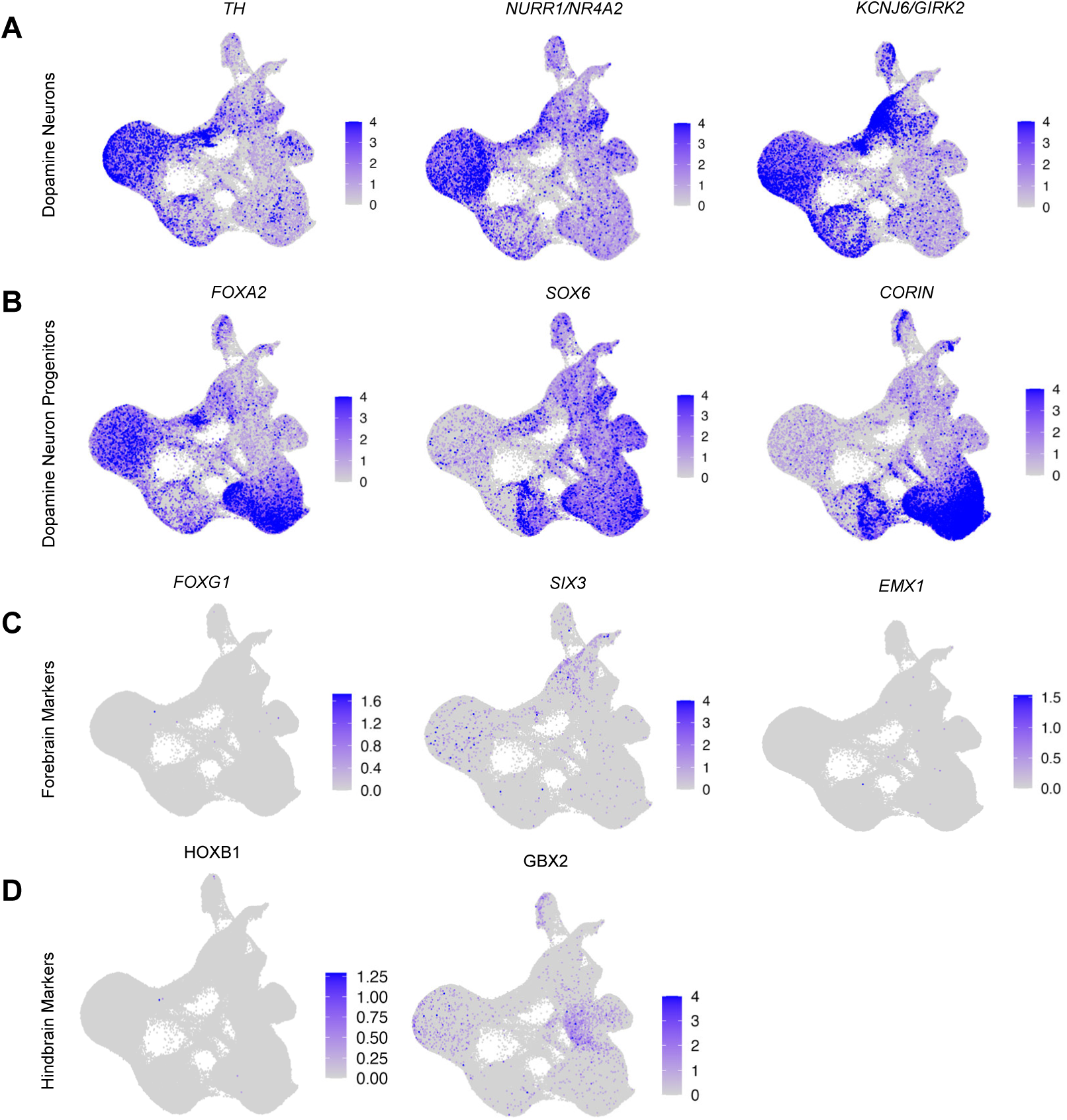
Expression of dopamine neuron and progenitor marker genes on the integrated iSCORE-PD scAtlas UMAP. A-B) Feature plots showing SCT-normalized expression of six canonical midbrain marker genes. **A)** Mature dopamine neuron markers; TH (tyrosine hydroxylase), enriched in the DA1 cluster with additional expression in DA2 and Early Neurons; KCNJ6 (GIRK2), a potassium channel; NR4A2 (NURR1), a transcription factor required for dopamine neuron identity and maintenance, broadly expressed across DA1, DA2, Early Neuron clusters. **B)** Midbrain floor plate and dopamine progenitor markers: FOXA2, a floor plate transcription factor expressed across both mature and progenitor populations, including DA1; SOX6, enriched in DA2; CORIN enriched in DAPro populations. **C-D)** Feature plots showing SCT-normalized expression of five canonical marker genes of forebrain and hindbrain. **C)** forebrain markers; FOXG1, a telencephalic transcription factor; SIX3, an anterior forebrain specification factor; EMX1, a dorsal telencephalon marker, all showing negligible expression across all clusters, confirming absence of forebrain identity. **D)** hindbrain markers; HOXB1, a rhombomere identity gene; GBX2, a hindbrain patterning factor, both showing minimal expression, confirming appropriate midbrain regional specification of the differentiated cells.

**Figure S3:**
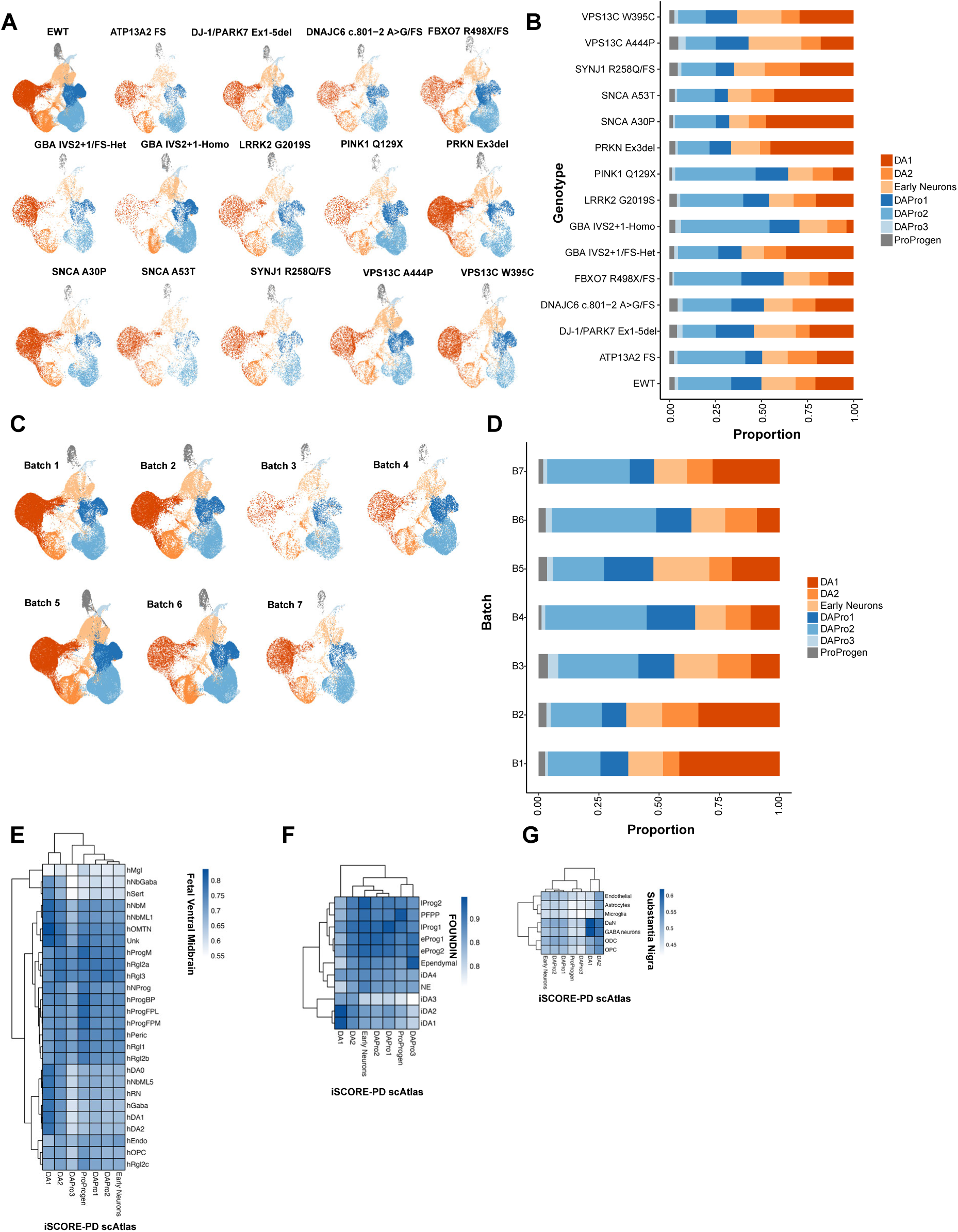
Cell type composition of iSCORE-PD dopamine cell populations across individual genotypes and Batches. A-C) UMAP projections showing the transcriptional landscape of CCA-integrated iSCORE-PD cells, displaying separately for each genotype and Batch. Each UMAP contains only cells from the indicated genotype, projected onto shared CCA-corrected UMAP embedding across all subclones. **B-D)** Stacked bar plots illustrating the relative distribution of the seven cell populations across 14 PD mutations and EWT controls (n=8). Each bar represents an individual genotype(B) and batch (D), demonstrating the high degree of consistency in the differentiation platform across mutations. Colors correspond to the annotated UMAP in A): bright orange, DA1; medium orange, DA2; light orange, Early Neurons; bright blue, DAPro1; medium blue, DAPro2; light blue, DAPro3; and grey, ProProgen. **E-G)** Correlation between iSCORE-PD scAtlas cell types and primary human and iPSC-derived reference datasets. Heatmaps showing pairwise Spearman correlation coefficients between pseudo-bulk average gene expression profiles of cell types and three independent reference datasets. **E)** Twenty-six annotated cell types from the human fetal ventral midbrain (La Manno et al., 2016; Spearman R= 0.55-0.85). **F)** Eleven annotated cell types from the FOUNDIN-PD, an iPSC-derived dopamine neuron dataset (Spearman R= 0.78-0.92); **G)** Seven annotated cell types from adult human substantia nigra (Agarwal et al., Spearman R=0.43-0.63). For all comparisons, correlations were computed using SCT-normalized expression averaged per cell type across all common genes detected in both datasets, whereas the FOUNDIN-PD where RNA-normalized expression was used. Rows and columns in each heatmap are hierarchically clustered; color scales from white (lower R) to deep blue (higher R). Across all three reference datasets, iSCORE-PD DA1 and DA2 neurons show the highest correlation with their respective dopamine counterparts (fetal hDA1/hDA2, iPSC-derived iDA1/iDA2, and adult DaN), while progenitor populations (DAPro1-3) align with developmentally progenitor cell types. Together, these cross-dataset comparisons validate the biological fidelity of iSCORE-PD cell type annotations across fetal, iPSC and adult human reference contexts.

**Figure S4:**
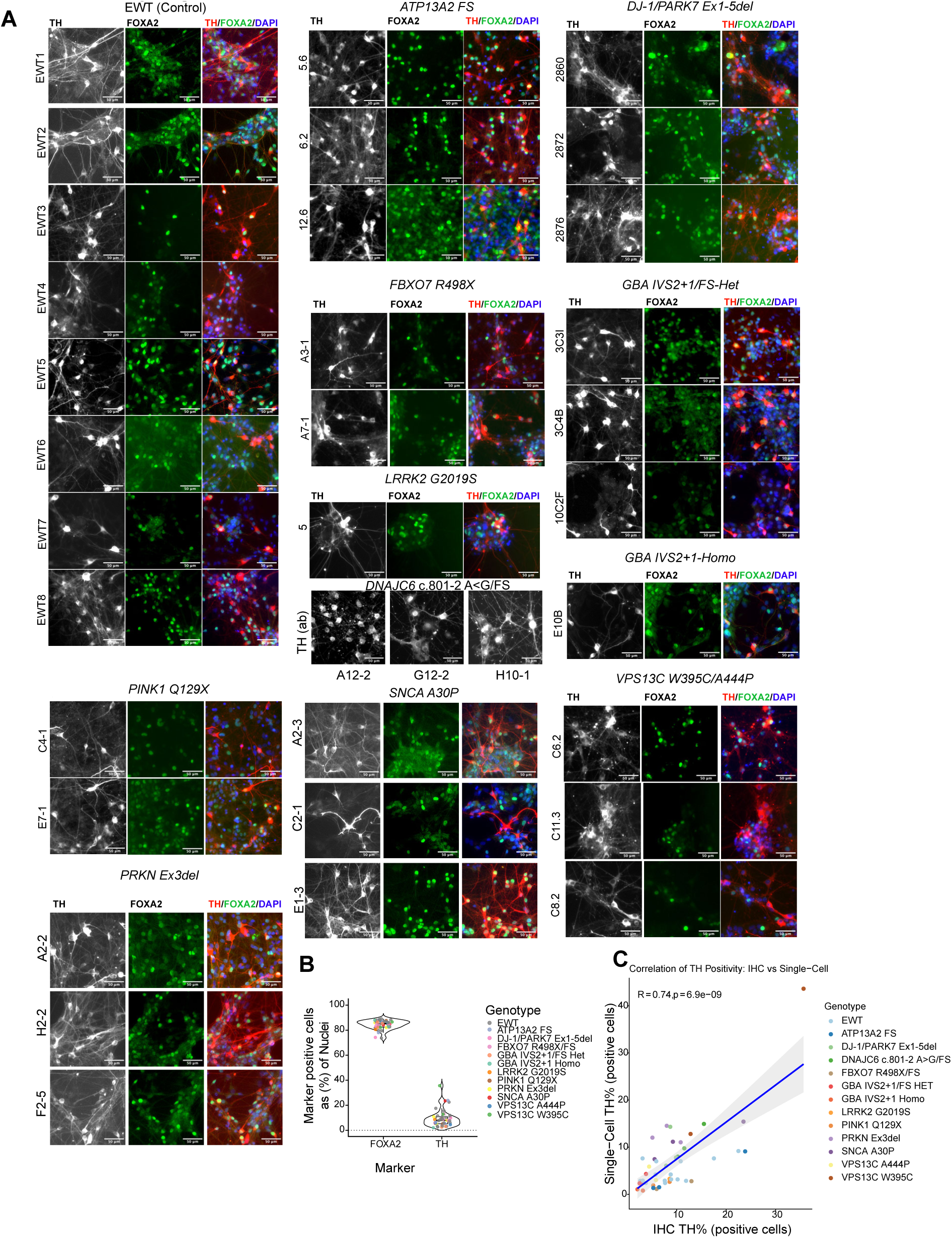
Immunofluorescence images of TH and FOXA2 staining in dopamine differentiation cultures. **A)** Fluorescence images showing TH and FOXA2 at day 35-45 of differentiation. DAPI (blue) marks all nuclei. Images are shown for all genotypes at least 1-2 subclones per genotype (SNCA A53T and SYNJ1 excluded due to suboptimal staining quality). Scale bar=50 µm. **B)** Violin plots showing the percentage of marker positive cells as proportion of total DAPI-counted nuclei, quantified using ImageJ analysis across all iSCORE-PD subclones. Each dot represents one subclone, colored by genotype. Embedded boxplots show the median (center line) and interquartile range (box), with whiskers extending to 1.5x IQR. TH (tyrosine hydroxylase), a rate-limiting enzyme in dopamine biosynthesis and canonical marker of dopamine neurons, was detected in the median of ∼8% of DAPI-positive cells across all genotypes. FOXA2 (Forkhead box A2), a floor plate and midbrain dopamine progenitor transcription factor that persists in mature DA neurons, was detected in a median of ∼85% of DAPI-positive cells, confirming high-efficiency midbrain floor plate induction. **C)** Concordance between immunofluorescence and single-cell transcriptomic quantification of TH-positive cells across iSCORE-PD subclones. Scatter plot showing the Pearson correlation between TH+ cell percentage quantified by immunofluorescence (IHC; x-axis) and TH-positive cell percentage from single-cell RNA-seq (y-axis), per subclone. Each dot represents one subclone, colored by genotype. The regression line (blue) with 95% confidence interval (grey shading) was fit by linear regression.

**Figure S5:**
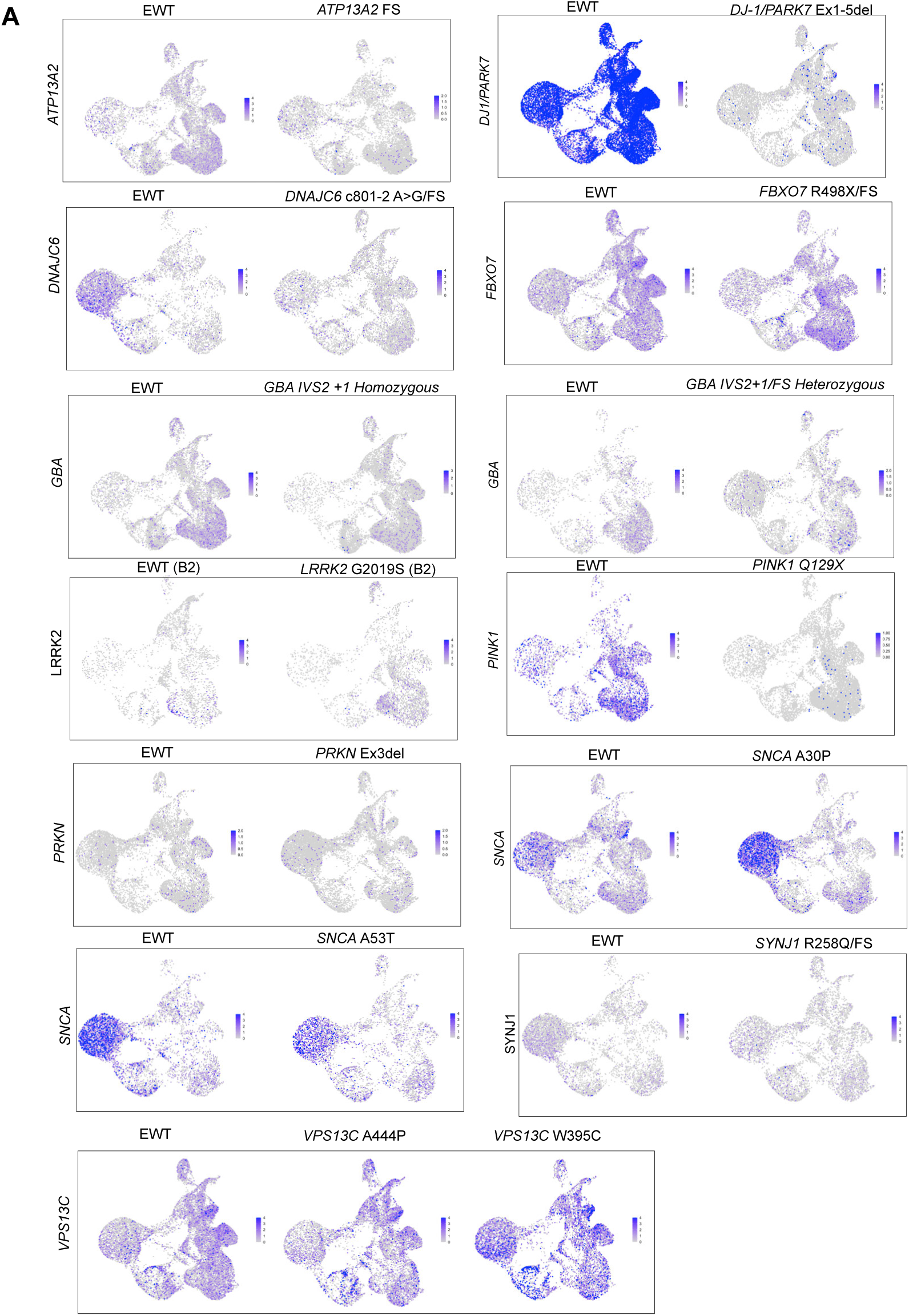
Expression of familial PD-associated genes in mutant and batch-matched EWT cells. Feature plots showing SCT-normalized expression of each familial PD associated gene in its corresponding mutation genotype alongside the EWT control from the same sequencing batch. Panel displays gene in alphabetical order (*ATP13A2, DJ-1/PARK7, DNAJC6, FBXO7, GBA, LRRK2, PINK1, PRKN, SNCA, SYNJ1, VPS13C*), each mutation genotype panel paired with its batch-matched EWT panel. Color intensity shows expression from gray (0, undetected) to deep blue (>=4). Batch-matched controls are used as the reference rather than a global EWT aggregate, to account for batch-batch variation in sequencing depth and cell type composition.

**Figure S6:**
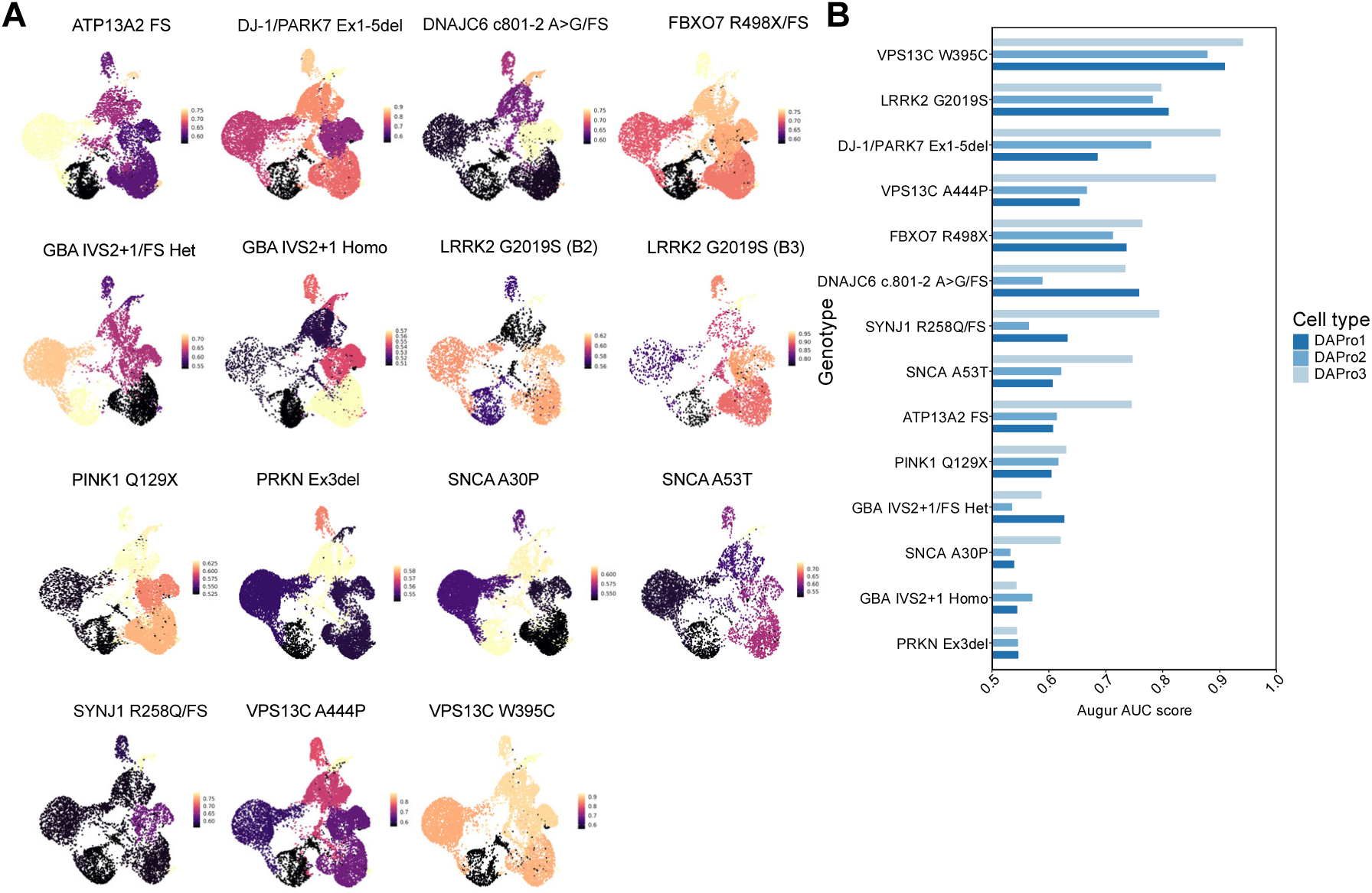
Cell-type resolved Augur perturbation scores overlaid on UMAP, displayed per genotype and batch. **A)** Feature plots showing Augur AUC scores for each mutant genotype on the CCA-integrated UMAP embedding, restricted to mutation cells only. Each panel represents one mutation-batch pair. LRRK2/G2019S, which was differentiated and sequenced across two batches (B2 and B3), is shown as two independent UMAPs. Augur AUC scores are calculated per cell type, visualized using a continuous magma color scale (low AUC=dark; high AUC= bright/yellow), reflecting the degree to which each cell type is transcriptionally distinguishable from its batch-matched EWT control. AUC=0.5 indicates no perturbation (random classifier performance); AUC=1.0 indicates strong transcriptional perturbation. UMAPs are arranged by genotype. **B)** Augur perturbation scores for dopamine progenitor populations across all familial PD mutations. Bar plot showing Augur scores (computed as described in S6A) for the three dopamine progenitor cell types (DAPro1, DAPro2, DAPro3) across all 14 familial PD mutations. For LRRK2 G2019S, which was processed in two batches, a single representative AUC score was calculated by taking the average across batches.

**Figure S7:**
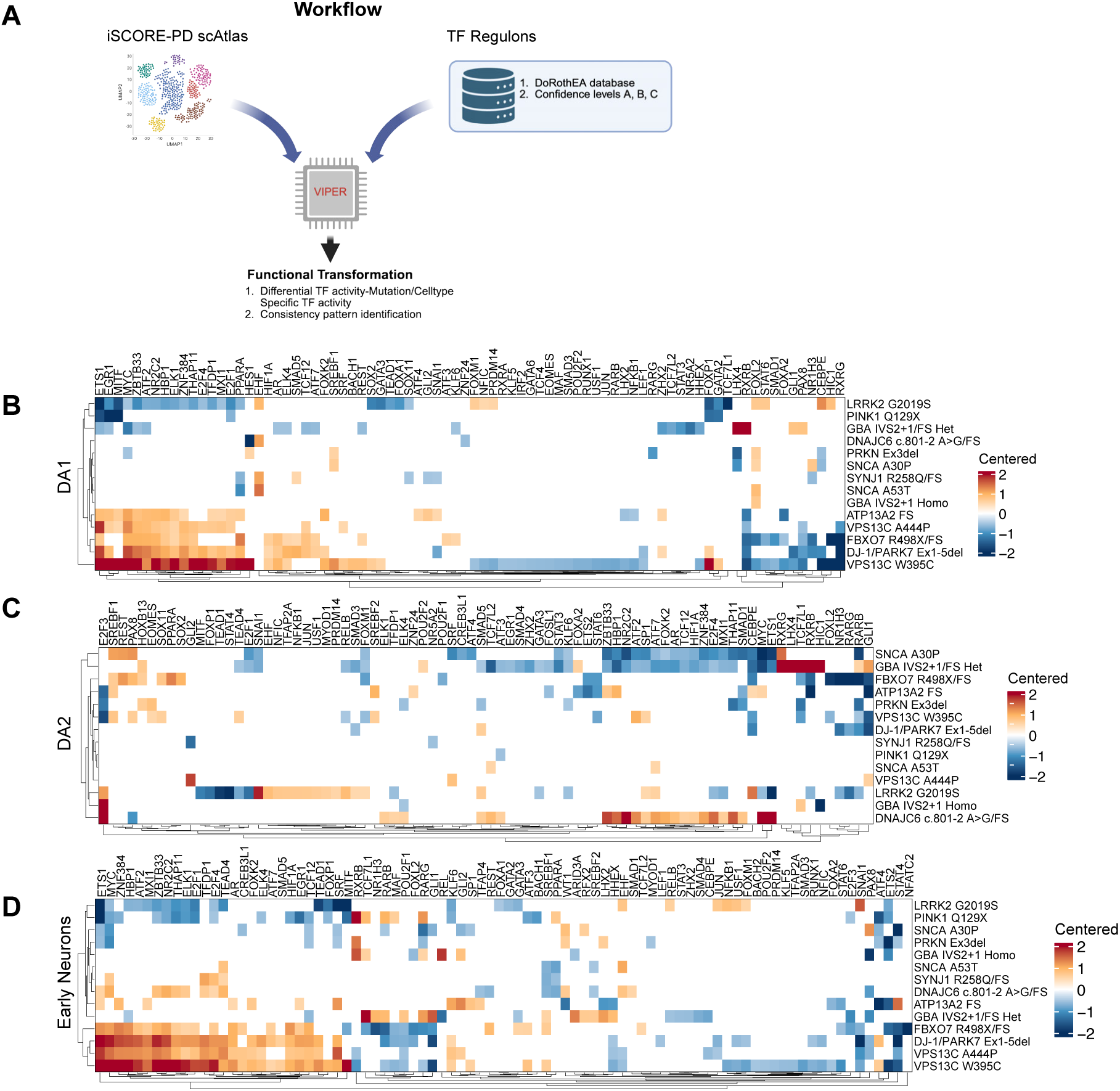
Systematic mapping of transcription factor activity identifies regulatory heterogeneity in PD neurons. **A)** Schematic representation of TF activity workflow. **B-D)** Differential TF activity landscape in DA1, DA2, Early Neurons. Heatmap displaying centered TF activity of dysregulated TFs (rows) across PD mutations (columns). TF activity was compared between each mutation and batch-matched EWT control. Centered log2FC was computed by subtracting the mean log2FC across all 298 TFs within each comparison, isolating TF-specific effects from global directional shifts. TFs were selected as the union of the top 10 upregulated and top 10 downregulated TFs per genotype (filtered at centered log2FC cutoff 0.5. Rows and columns are hierarchically clustered (Euclidean distance, complete linkage).

**Figure S21:**
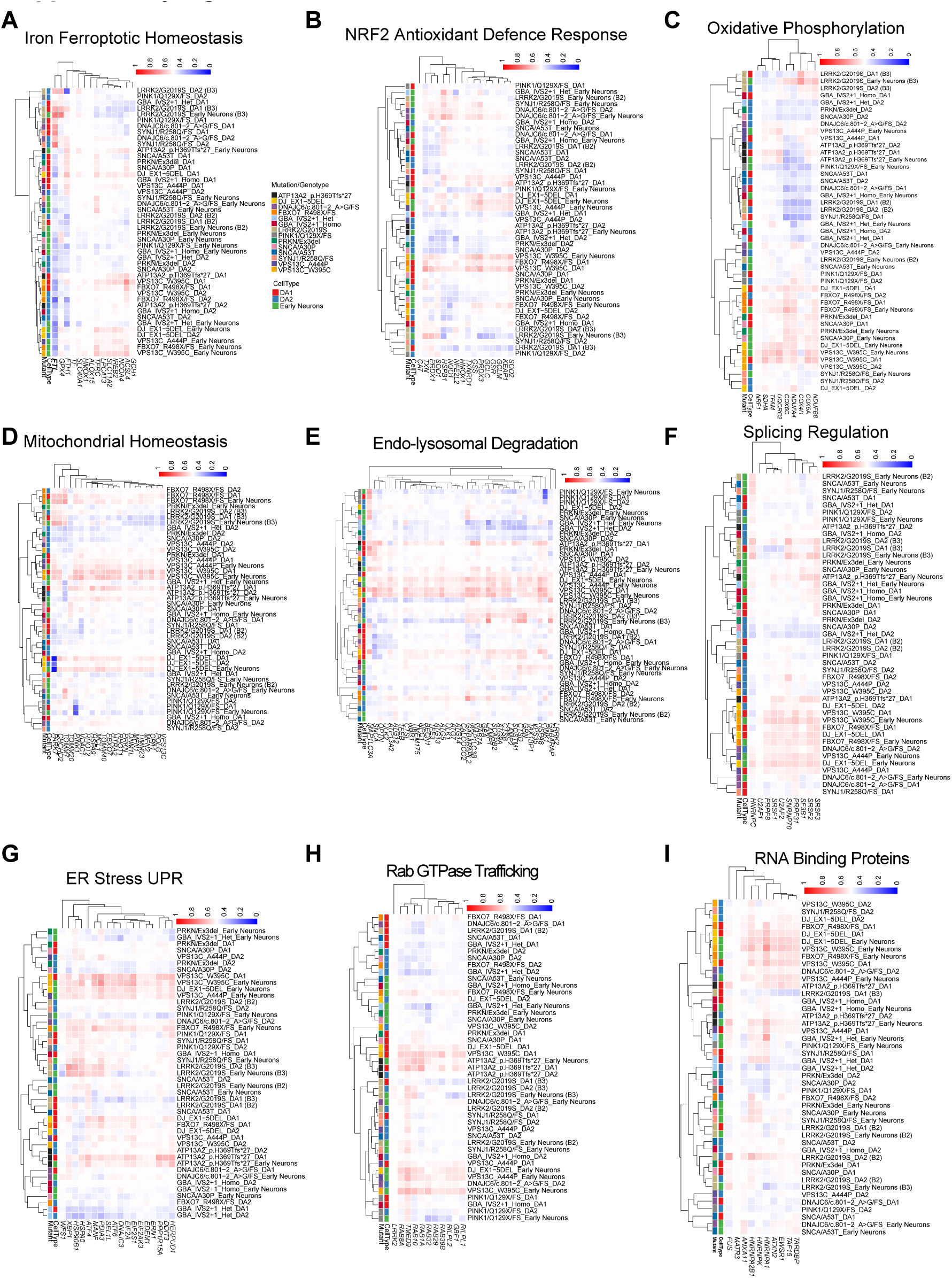
Heatmaps of AUC values across selected PD relevant gene sets. Heatmaps show AUC values for selected curated gene sets (full data in Table S10) across DA1, DA2, and Early Neurons. For each batch-matched comparison, mutation cells were compared against EWT cells, and AUC was calculated from single-cell log-normalized RNA expression to quantify directional separation between mutation and control cells. Rows represent individual mutation-cell type comparisons, and columns represent genes within each indicated gene set. AUC values >0.5 indicate relatively higher expression in mutation cells, whereas values <0.5 indicate relatively lower expression compared to EWT controls. Rows and columns were hierarchically clustered using correlation-based distance.

**Figure S22:**
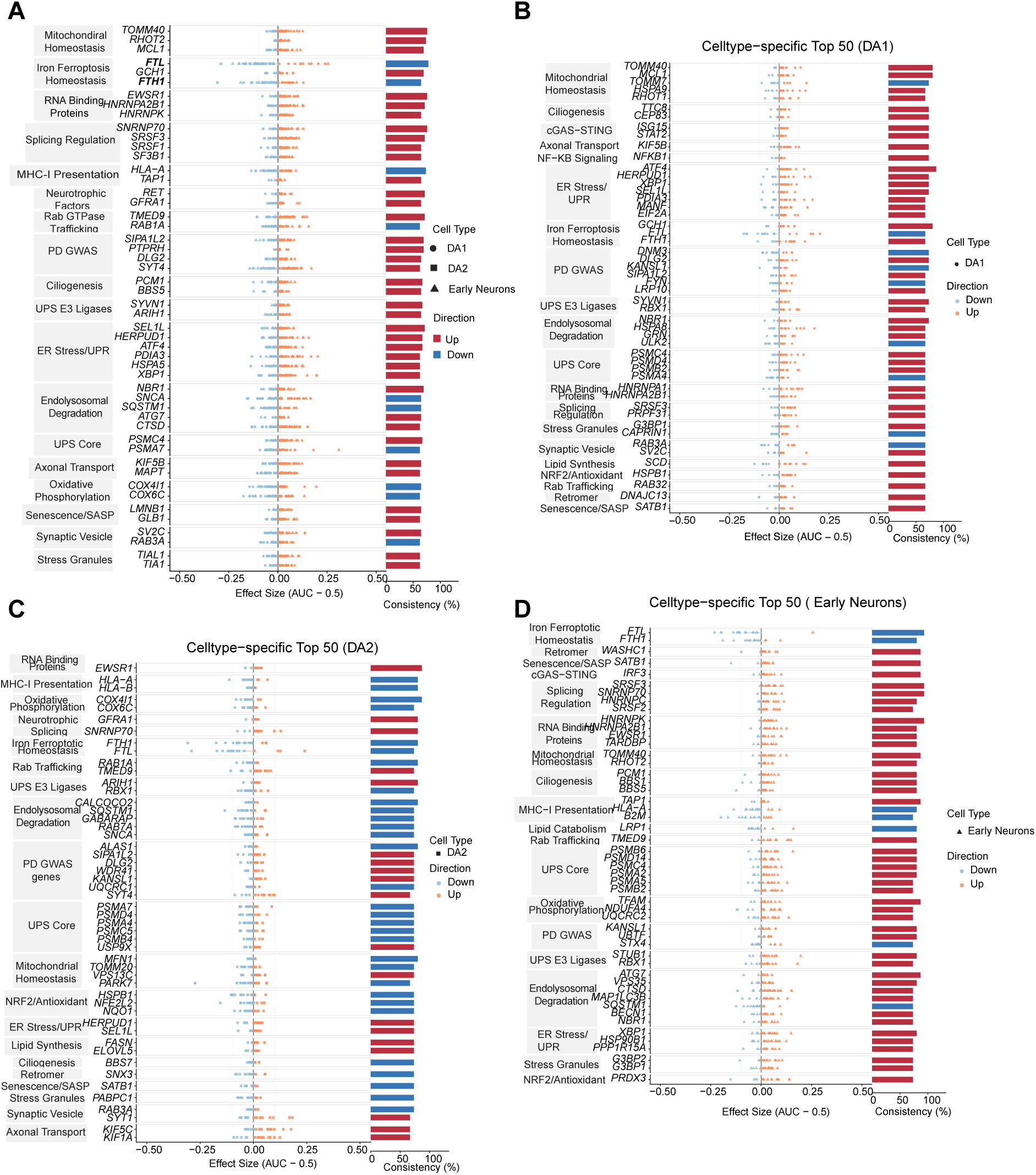
Top 50 consistently dysregulated genes across familial PD mutations, ranked pooled across cell types and per cell type. **A)** Individual AUC values for each genotype-cell type comparison are shown for the most consistently dysregulated genes across all three dopamine neurons populations combined (DA1, DA2, Early Neurons). Points are colored by direction (red: AUC>0.5, upregulation; blue: AUC<0.5, downregulation) and shaped by cell type. Grey horizontal bars indicate the mean effect size per gene (mean centered AUC). Right bars represent directional consistency, defined as the fraction of genotype-cell type comparisons dysregulated in the predominant direction (up to n=42 comparisons per gene; 14 familial PD mutations x 3 cell types). **B-D)** Cell-type-specific top 50 genes, same visualization as in (A), but genes are independently selected and ranked per cell type (DA1, DA2 and Early Neurons shown separately), displaying the 50 most consistently dysregulated genes specific to each population (up to n=14 comparisons per gene,14 familial PD mutations within a single cell type). Consistency bars reflect per-cell type directionality for each gene.

**Figure S23:**
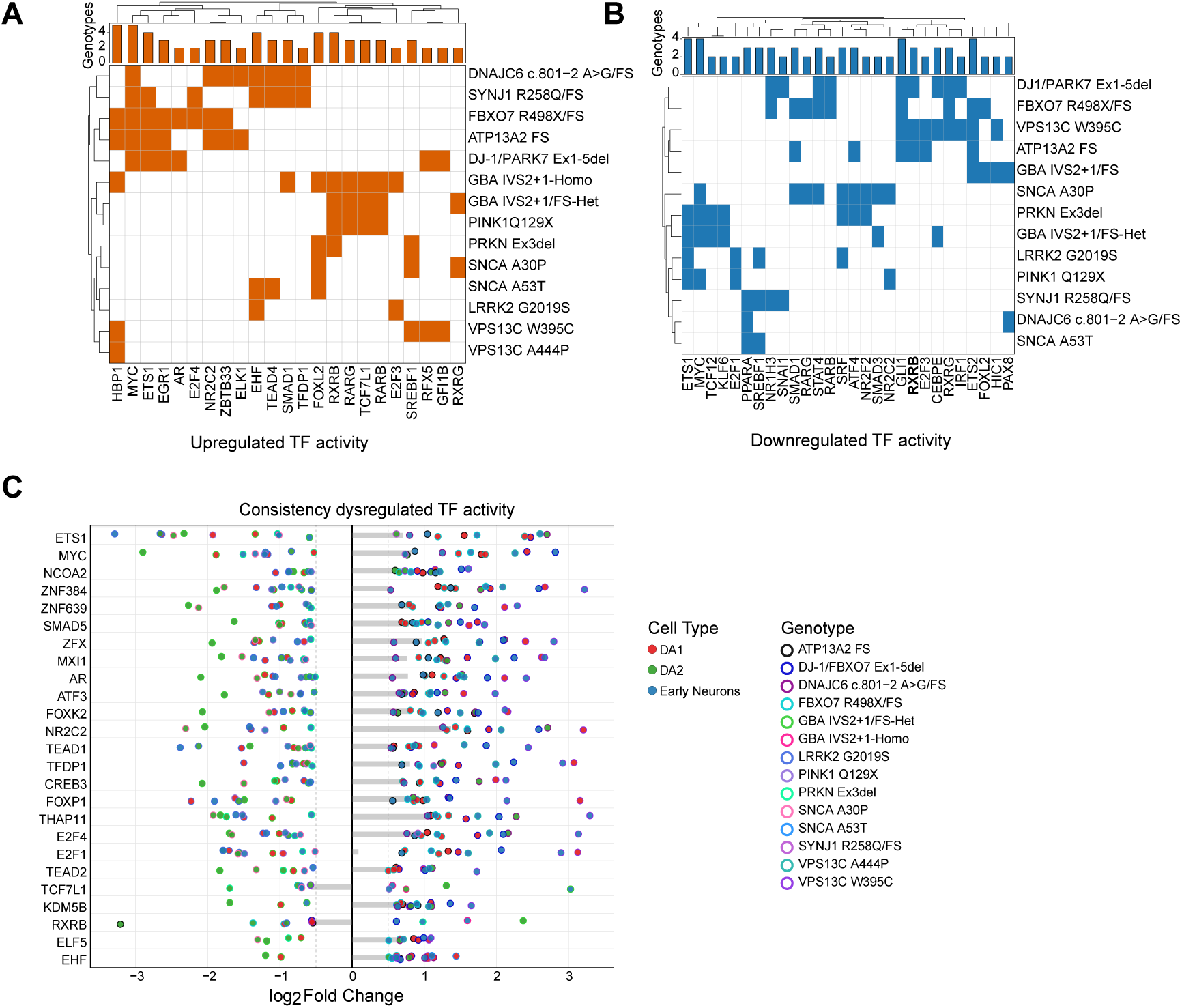
Common TFs dysregulated in neuronal populations. A-B) Heatmaps showing top dysregulated TFs overlap across familial PD genotypes. Heatmaps showing which TFs are among the top 10 most dysregulated per genotype, displayed separately for upregulated (left, orange) and downregulated (right, blue) TFs. For each genotype, the top 10 upregulated and top 10 downregulated TFs were identified by differential TF activity, pooled across DA1, DA2, and Early Neurons; each TF is counted once per genotype regardless of the number of cell types in which it was dysregulated. Columns represent TFs filtered to those present in at least two genotypes; rows represent genotypes. Both rows and columns are hierarchically clustered (Euclidean distance, complete linkage). **C)** Top 25 consistently dysregulated TFs across neuronal populations. Horizontal dot plot showing log2FC (x axis) for the 25 most consistently dysregulated TFs (y-axis) across DA1, DA2, Early Neurons. Each dot represents a significant TF-mutation combination (cutoff log2FC 0.5); dot fill indicates cell type (red=DA1, green = DA2, blue= Early Neurons), and dot border indicates genotype. TFs were ranked by a composite score integrating directional consistency, effect size, number of cell types affected, and total significant combinations, requiring consistency in at least two cell types and 10 significant combinations across all cell types and mutations (each TF-mutation-cell type combination counted once).

**Figure S24:**
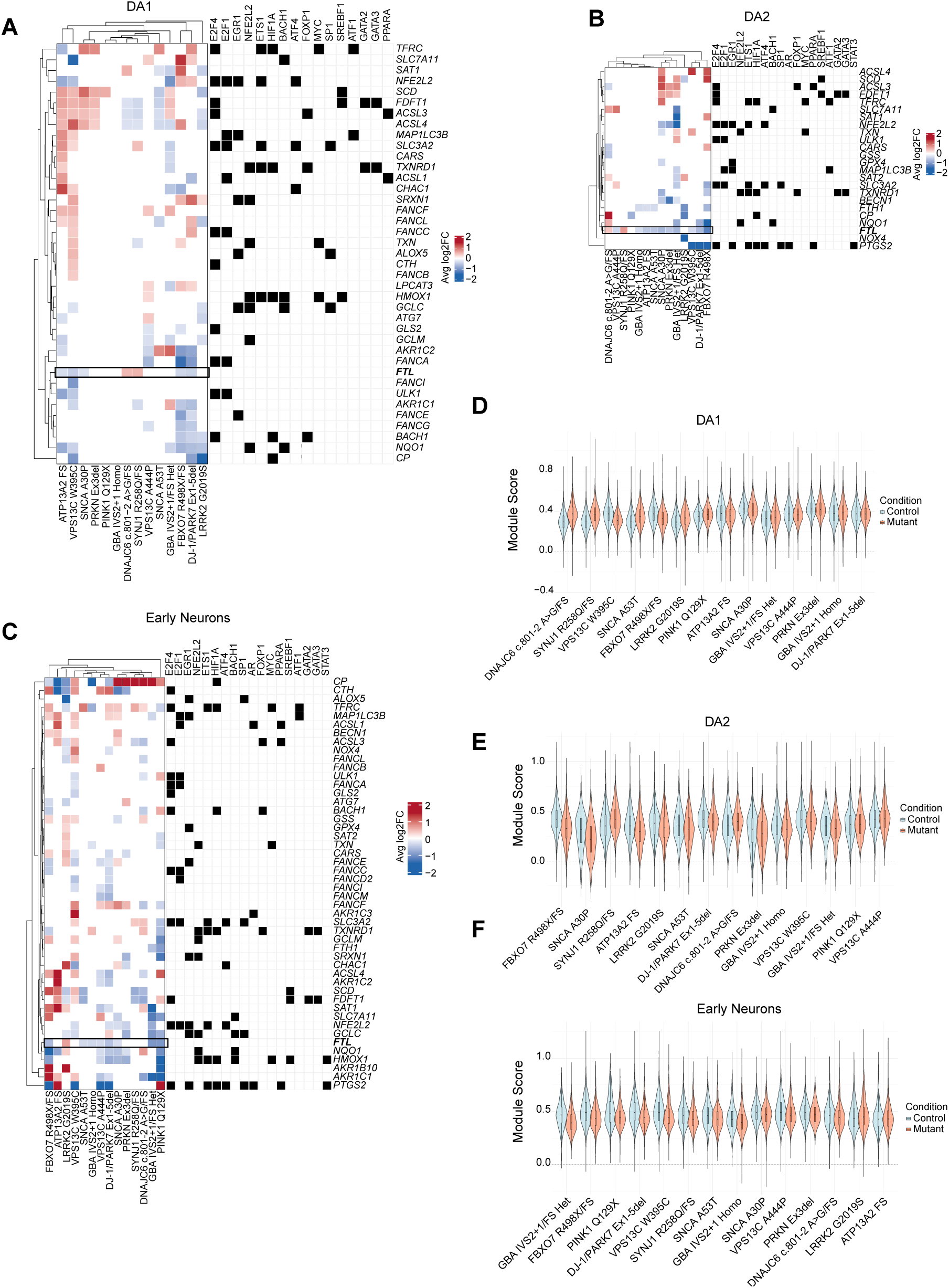
Ferroptosis gene set dysregulation and module-level activity across familial PD mutations in neuronal populations. A-C) Heatmaps showing differentially expressed genes from the WikiPathways ferroptosis gene set (rows) across familial PD mutations (columns) in (A) DA1, (B) DA2, and (C) Early Neuron cell types. Each cell shows avg log2FC for that gene in mutant against batch-matched EWT. Only DE genes (FDR<0.05) in at least mutation) intersecting the WikiPathways ferroptosis gene set are shown. Rows and columns are hierarchically clustered. The right-side annotation indicates DoRothEA regulon membership; filled squares mark genes that are predicted targets of the indicated TFs, with the TFs drawn from the top dysregulated TFs identified in the TF activity analysis (Fig S23). **D-F)** Violin plots showing per-cell ferroptosis module scores (D) DA1, (E) DA2, and (F) Early Neuron cell types across the familial PD mutations. Module scores were computed with Seurat’s *AddModuleScore()* function using a curated ferroptosis gene set (independent from the WikiPathways gene set used in A-C; see Table S10 for full data on other gene sets). Batch-matched EWT cells (blue is control) are plotted alongside mutant cells (orang is mutant) for each mutation, with genotypes ordered along the x-axis.

**Figure S25:**
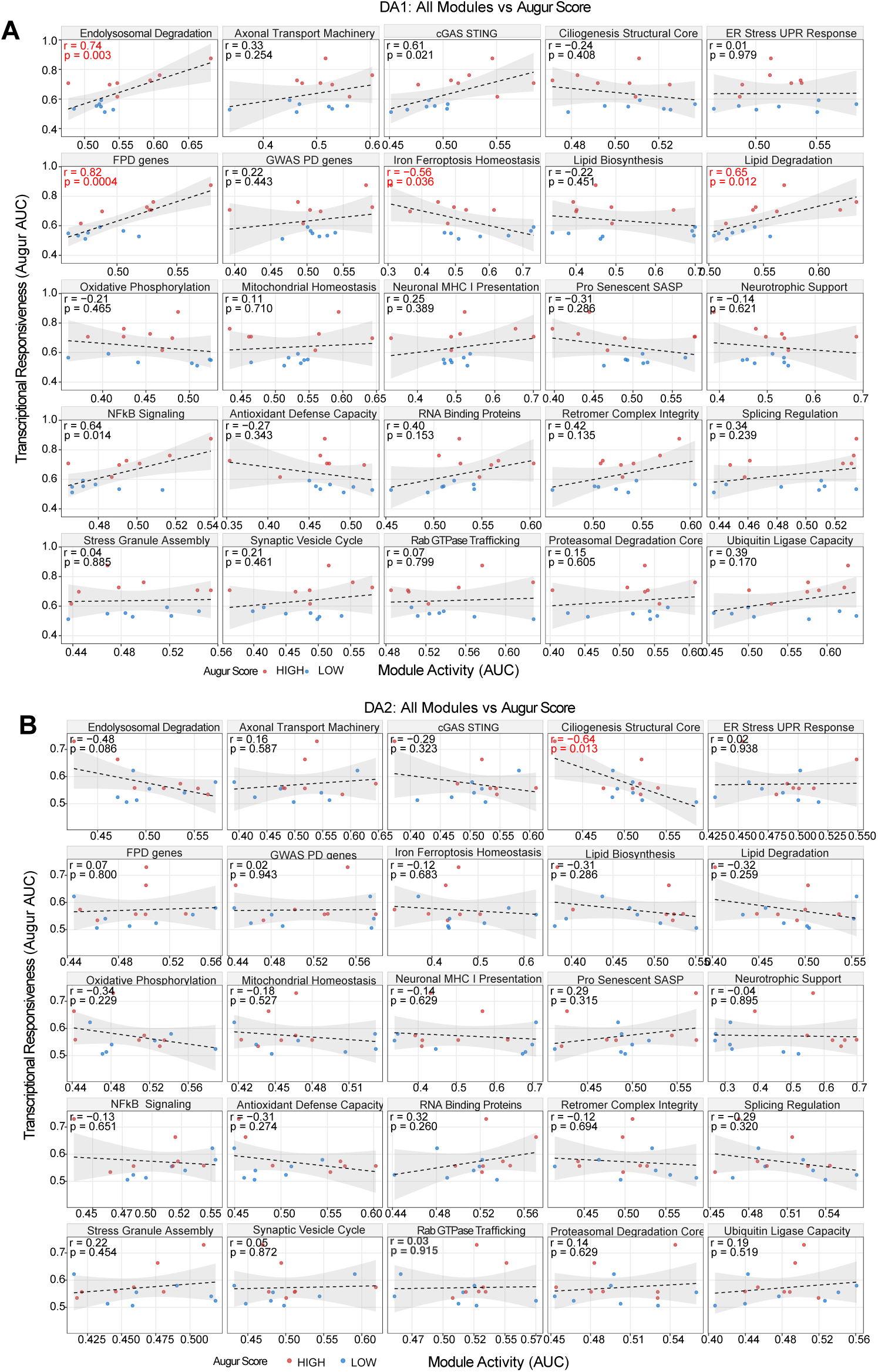
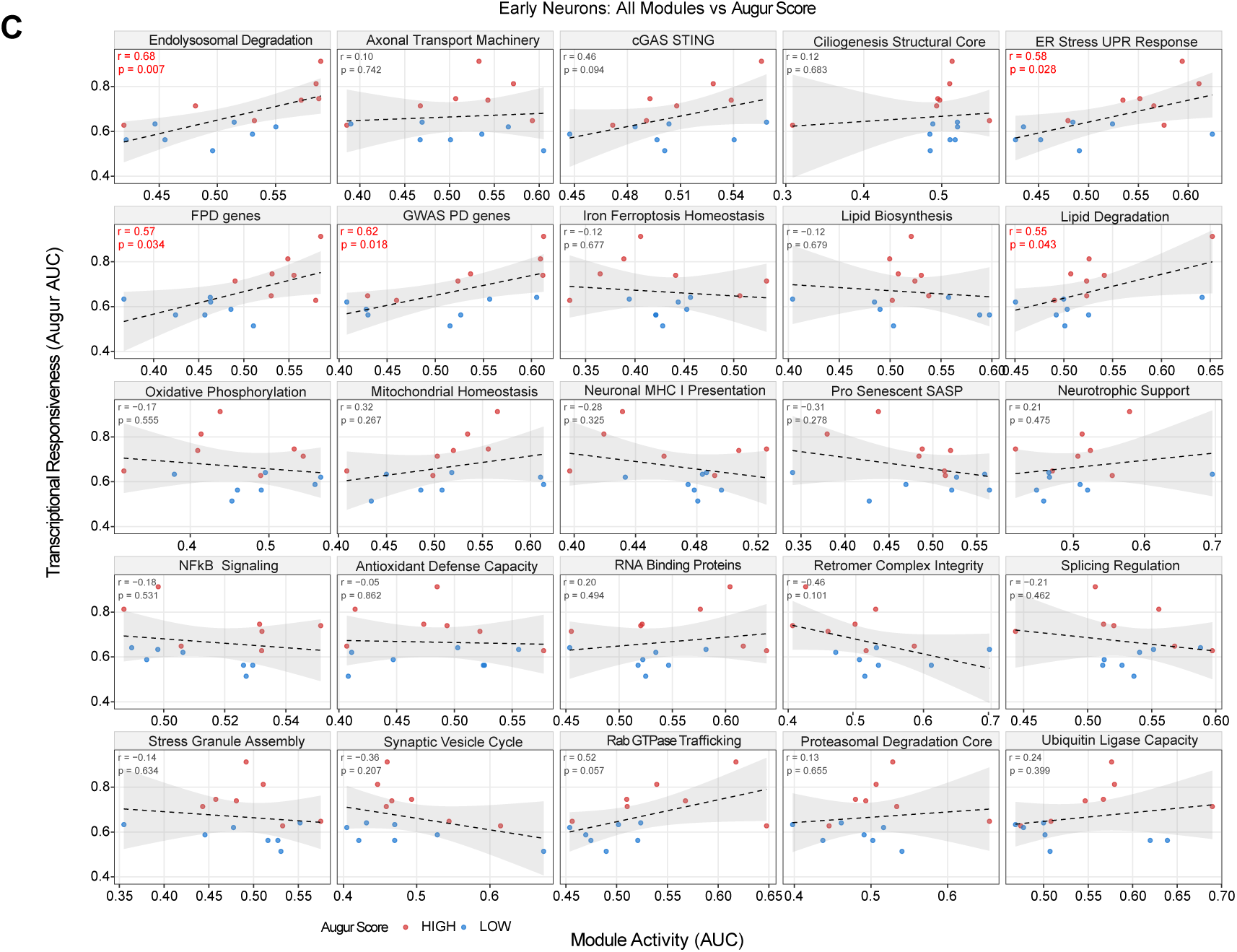
Cell-type specific correlation between PD-relevant module dysregulation and genotype vulnerability across familial PD mutations. A-C) Scatter plots showing the relationship between pathway module activity and cellular transcriptional responsiveness for each dopamine neuron population (DA1, DA2, Early Neurons; shown as separate panels). Within each cell-type, sub-panels represent one of the 25 PD relevant modules. Each dot represents a genotype (14 familial PD mutations), colored by transcriptional perturbation (red: HIGH; blue: LOW). Vulnerability classification was determined by calculating the mean Augur AUC for each genotype across DA1, DA2, and Early Neurons combined, then applying a median split; genotypes above the median were classified as HIGH vulnerability and those below as LOW vulnerability. The x-axis shows module activity, defined as the mean directional AUC score for that module within the given cell type and genotype. The y-axis shows cell-type specific vulnerability as measured by Augur AUC. A dashed linear regression line with 95% confidence interval is shown. Pearson correlation coefficient (r) and p value are annotated within each sub-panel. Sub-panels with significant correlations (p<0.05) are highlighted in red.

**Figure S26:**
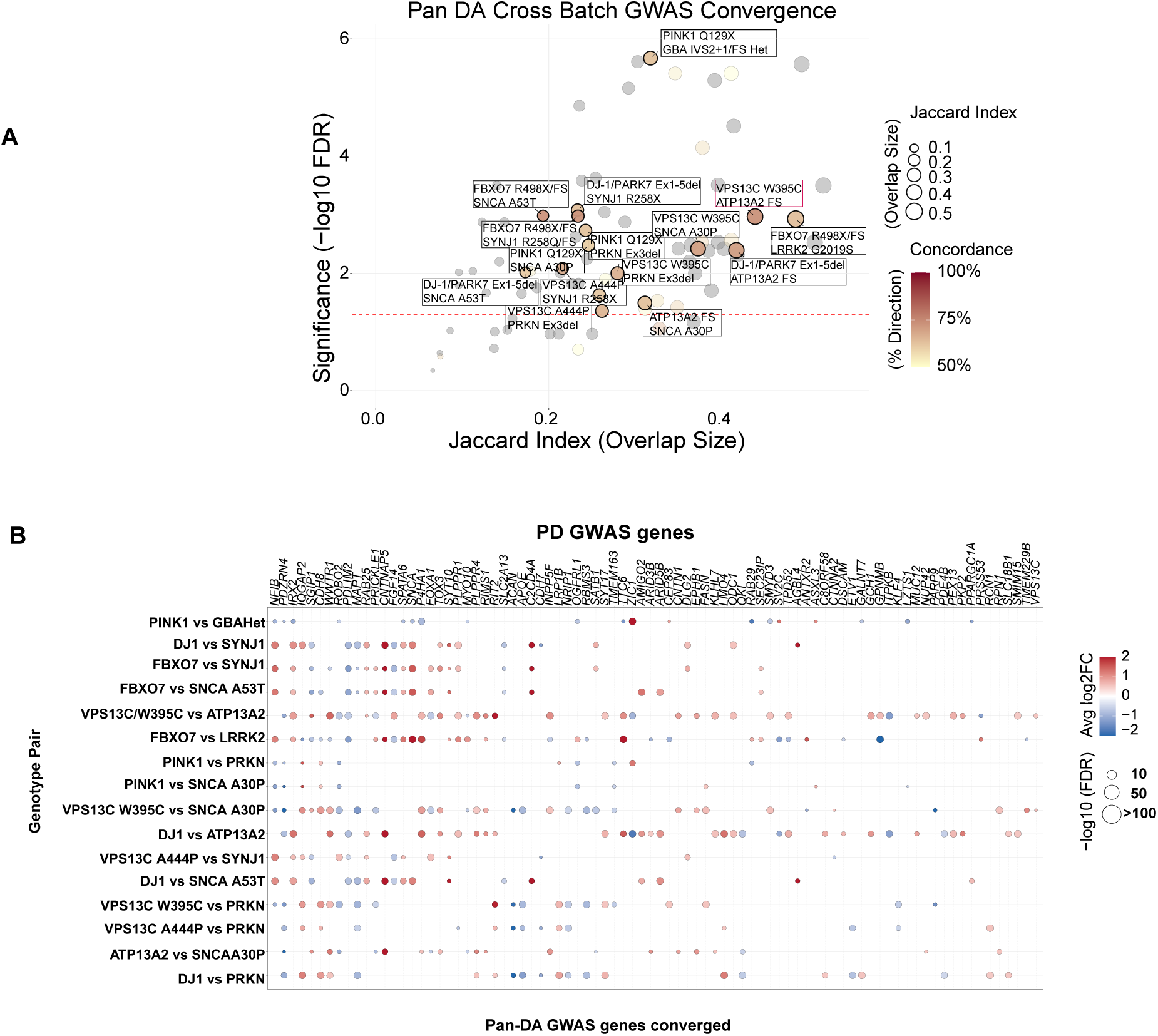
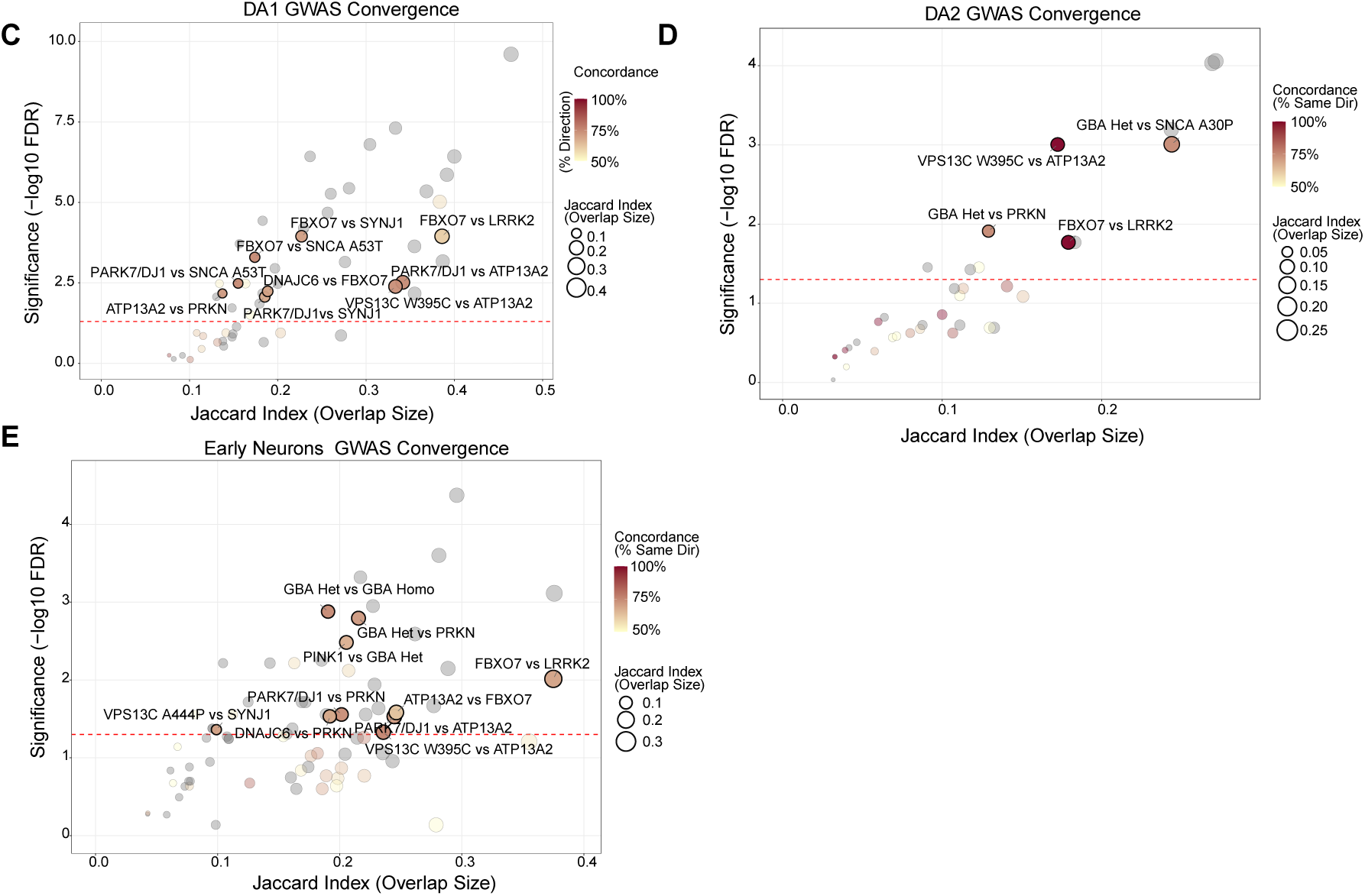
GWAS-nominated gene convergence across cross-batch mutation pairs. **A)** Cross-batch pairwise convergence of familial PD mutations on shared GWAS risk genes. Scatter plot showing pairwise convergence of GWAS-overlapping DE gene sets between all cross-batch genotype pairs (Pan-DA analysis). X-axis is Jaccard index quantification of shared GWAS-DE gene perturbation between two genotypes. Dot color indicates the proportion of shared GWAS-DE genes dysregulated in the same direction by both mutations, ranging from yellow (50%, no directional preference) to dark red (100%, fully concordant). Dot size indicates Jaccard Index. Pairs were classified as convergent (black-colored borders) with FDR<0.05 and concordance atleast 60%. Significant convergent pairs were labelled. Genes appearing in multiple cell types were retained only if directional effects were concordant across cell types within each genotype. For each genotype, genes appearing in multiple cell types (DA1, DA2, Early Neurons) required directional concordance. **B)** Dot plot displaying individual GWAS genes (y-axis) shared across cross-batch significant convergent pairs (x-axis). From the Pan-Cellular convergence analysis, convergent pairs are identified. For each convergent pair, overlapping GWAS-DE genes were extracted and filtered to those with concordant directionality (both genotypes) and magnitude of average log2fc cutoff 0.5. Only genes associated with at least two pairs are shown. Dot fill color indicates the average log2FC across both genotypes in the pair (blue=downregulated; white=unchanged; red= upregulated). Dot size indicates the statistical significance. Genes are ordered by the number of pairs in which they appear. Pairs are ordered by overlap significance. Note: This plot limits the visualization to genes with robust effect sizes across both genotypes, with average threshold cutoff of log2FC 0.5 across the pairs; this filter selectively retains genes with consistently moderate to strong dysregulation in both mutations. **C-E)** Cell-type-specific GWAS gene convergence across cross-batch mutation pairs in DA1, DA2, and Early Neurons. Scatter plots showing pairwise GWAS-nominated gene convergence calculated separately for each dopamine neuron cell type: **C)** DA1, **D)** DA2, **E)** Early Neurons. Each point represents one cross-batch genotype pair. The x-axis shows the Jaccard index of the overlapping gene sets between two genotypes. The y-axis shows -log10 FDR from one-sided hypergeometric test. Point color shows directional concordance of overlapping genes, light yellow (50%, equivalent to chance) to dark red (100% same direction). Size indicates the Jaccard index. The dashed red line marks the FDR<0.05 significance threshold. Circles above this line with concordance >=60% are classified as significant pairs (circles with black outline) and labelled with genotype pair name (top 15 by significance shown).

**Supplementary Note S26:** Mutations differentiated within the same batch can show inflated transcriptional similarity due to shared technical variation in cell-state composition, which confound true biological convergence. As a result, we restricted analysis to mutation pairs profiled in different differentiation batches, where any observed overlap is independent of shared technical conditions and therefor reflects shared biological signal rather than artifacts. Beyond the VPS13C W395C and ATP13A2 pair noted in the main text, the cross-batch analysis revealed additional mechanistically informative overlaps. The strongest overall convergence linked PINK1 and heterozygous GBA (40 shared genes, 62% directional concordance), bridging mitochondrial quality control and lysosomal dysfunction. Within-related closely related PINK1 and PRKN, we have observed convergence (24 shared genes, 62% concordance) along the mitochondrial axis.

**Figure S27:**
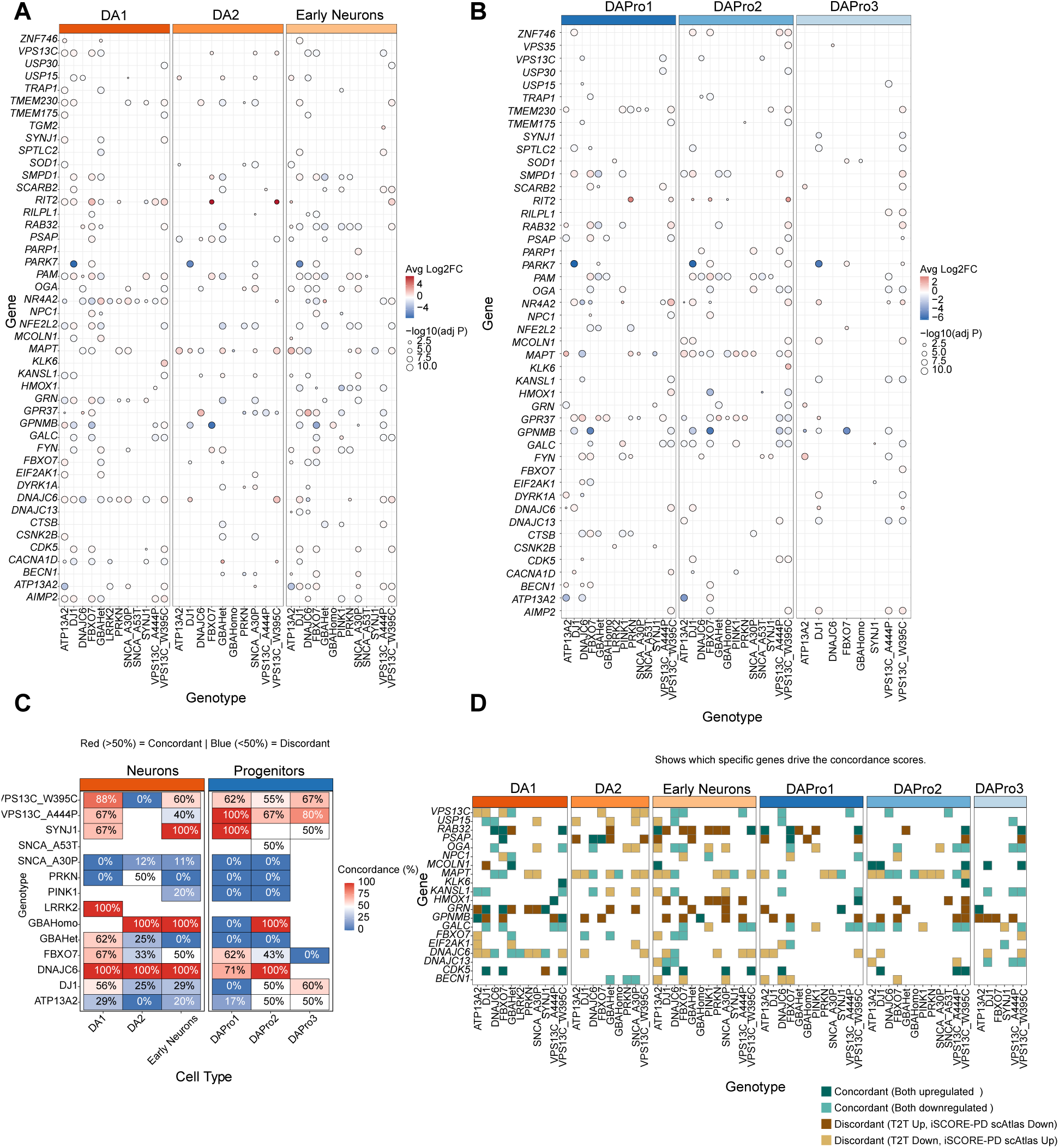
Integrative analysis of transcriptomic concordance with MJFF T2T Parkinson’s Disease Target Genes. A-B) Expression profiles of T2T target genes in neuronal and progenitor populations. Dot plot representing the differential expression of genes selected from the MJFF. Top 59 target list across 14 PD mutations and six cell populations. The size of each dot represents the statistical significance (adjusted p-value), and the color represents the magnitude and direction of the average fold change (red=upregulated, blue= downregulated). Data shown are significance (FDR <0.05) and log2FC threshold 0.25. **C)** Directional concordance with T2T disease state. Heatmap showing the percentage of overlapping differentially expressed genes that match the expected PD expression direction (Up or Down) as defined by the T2T list. Heatmap cells are colored by concordance percentage: red (>50%) indicated most genes match the expected disease state (concordant), while blue (<505) indicates the majority of genes show the opposite direction (discordant). Labels with 0% indicate a complete reversal of expected disease signals, while 100% indicates perfect directional alignment with the T2T reference. **D)** Directional state across cell populations. A detailed categorial heatmap illustrating the specific directional relationship between iSCORE-PD scAtlas observations and T2T expectations for each gene. Genes are categorized into four states.

**Figure S28:**
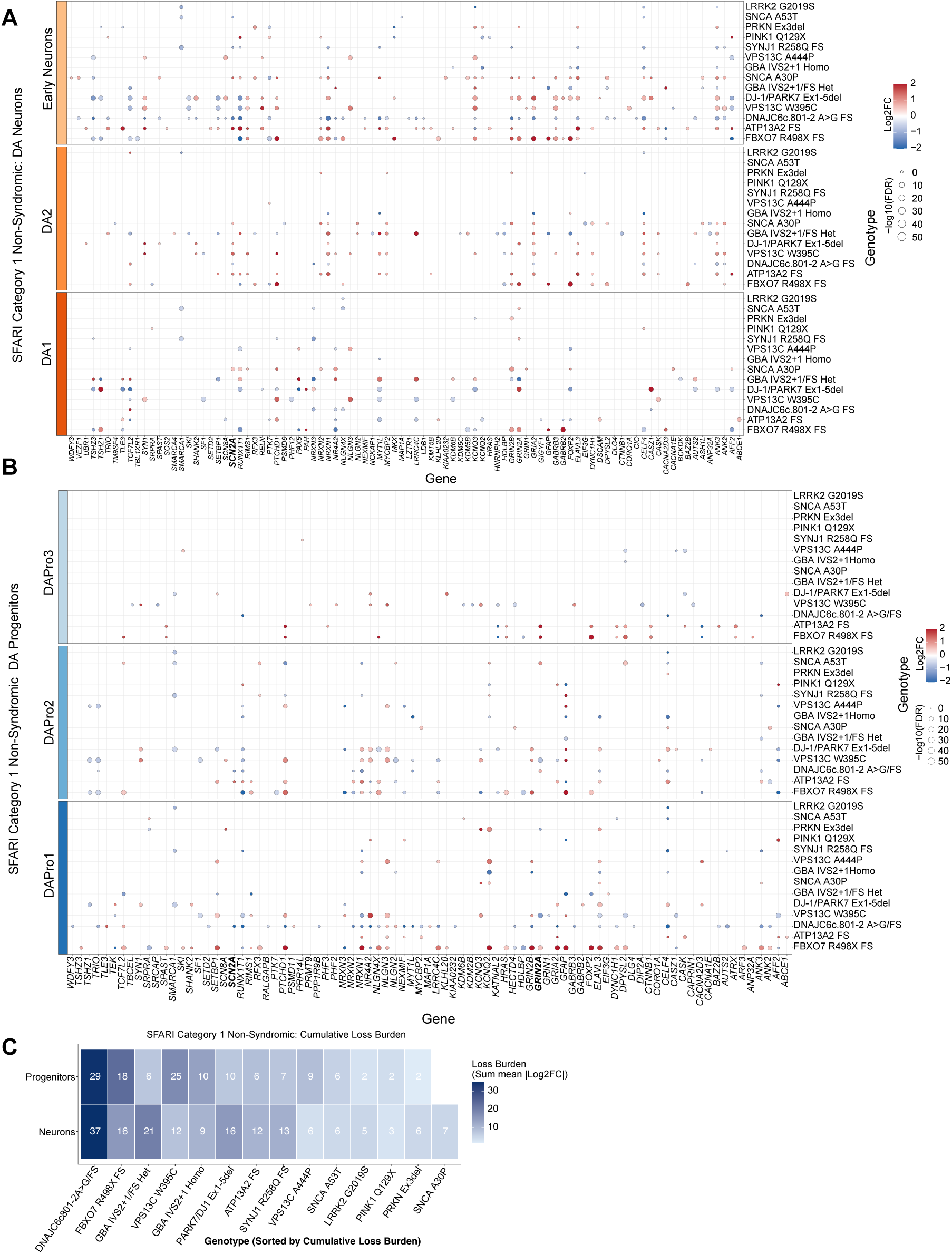
Dysregulation of SFARI non-syndromic ASD genes in iSCORE-PD dopamine populations. **A)** Dot plot showing differential expression of SFARI category 1 non-syndromic ASD genes (n=123 high-confidence non-syndromic ASD risk genes; gene-score =1, syndromic =0 in SFARI Gene database 2025 Q4) across all 14 familial PD mutations in dopamine neuron cell types (DA1, DA2, and Early Neurons). Each row is one gene, and the column represents a genotype, ordered by decreasing cumulative dysregulation magnitude (sum of log2FC). Log2FC threshold 0.5 and FDR cutoff <0.05 is used. **B)** As described in S27A, shown in dopamine progenitor populations (DAPro1, DAPro2, DAPro3). **C)** Heatmap summarizing the cumulative downregulation burden of non-syndromic ASD genes, for DA neurons and progenitors. Each tile represents one genotype-cell-type group combination. The color indicates loss-of-function burden as the sum of absolute log2FC values across all significantly downregulated genes. Numbers indicate the count of unique downregulated genes contributing to each tile. Note that gene count and burden sum correlate strongly across genotypes, as mutations affecting more ASD genes accumulate higher aggregate log2FC burden.

**Figure S29:**
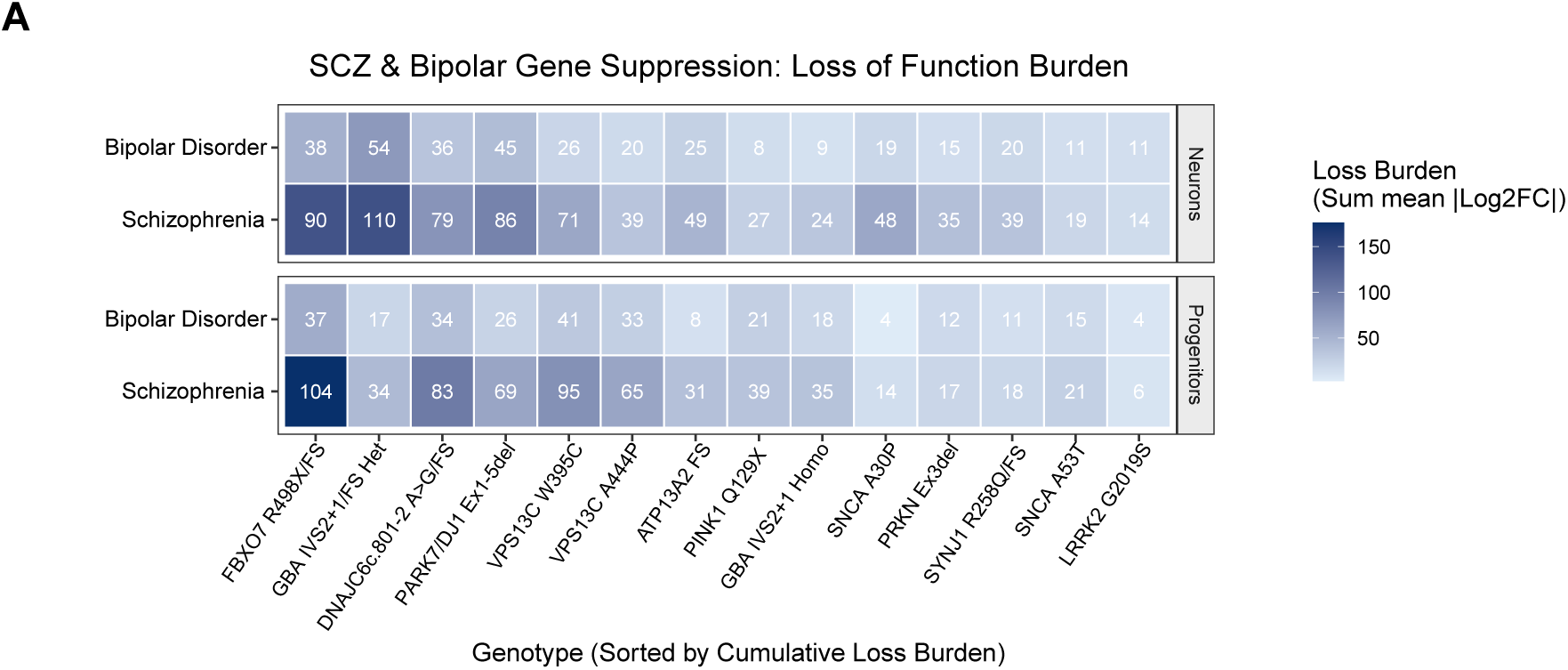
Schizophrenia and bipolar disorder gene suppression burden across iSCORE-PD genotypes. Heatmaps summarizing the cumulative dysregulation of DisGeNET high-confidence schizophrenia (SCZ; n= 497 genes, GDA score >=0.6) and bipolar (BD; n=204 genes, GDA score >=0.6) risk genes across all 14 familial PD mutations, separately for neuronal (DA1, DA2, Early Neurons) and progenitor (DAPro1, DAPro2, DAPro3) populations. Loss-of-function burden: tile color indicates the sum of mean absolute log2FC across downregulated disease risk genes (FDR <0.05, log2FC 0.5 cutoff). Annotated numbers indicate the count of dysregulated genes per genotype per category. Genotypes are ordered in decreasing order of total burden.

## Supplementary Materials

**Figure S8:**
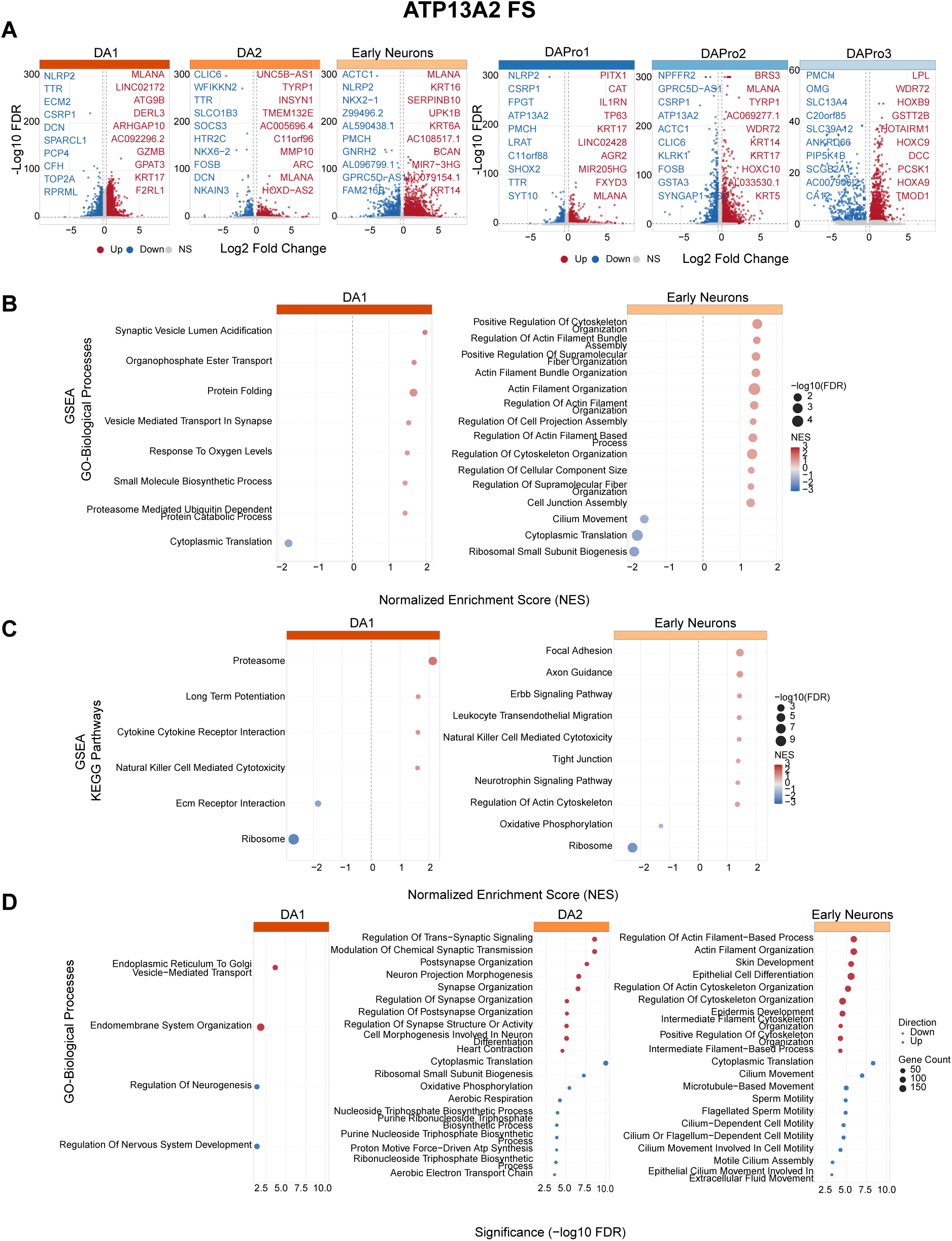
Differential expression and pathway enrichment analyses for each familial PD mutation across iSCORE-PD cell types. One supplementary figure is provided per mutation (S8: *ATP132* FS; S9: *DNAJC6* c.801-2 A>G/FS; S10: *FBXO7* R498X/FS; S11: *GBA* IVS2+1 heterozygous; S12: *GBA* IVS2+1 homozygous; S13: *LRRK2* G2019S; S14 *PINK1* Q129X; S15: *PRKN* Ex3 del; S16: *SNCA* A30P; S17: *SNCA* A53T; S18: *SYNJ1* R258Q/FS; S19: *VPS13C* A444P; S20: *VPS13C* W395C). Each figure contains four panels described below. All analyses compared each mutation genotype with the wild-type control (EWT) in the same batch. Differential expression was performed at the single-cell level using MAST. No latent variables were included, because each comparison was restricted to cells from the same batch, thereby controlling batch effects by design. EWT subclones within a given batch served as the reference population for each mutation. **A)** Differential gene expression. Volcano plots showing DE results for each annotated cell type, split into two groups: Neuronal populations and progenitor populations. X-axis shows average log2 Fold change; Y-axis shows -log10 (FDR-adjusted p-value). Upregulated genes are in red, downregulated in blue. The top 10 genes per direction per cell type ranked by log2FC are labeled. **B)** Gene Set Enrichment Analysis (GSEA)-GO biological processes: Dot plot showing GSEA results in DA neuron populations (DA1, Early Neurons; DA2 is omitted when no gene set passes the FDR <0.05 threshold). Genes were ranked using p-value and log2FC combined, capturing both effect direction and significance. **C)** GSEA, dot plots showing KEGG pathways: Same parameters used as described in B. **D)** Over-Representation Analysis (ORA), GO-BP: Dot plot showing ORA results for GO: BP terms. The top 15 terms per cell type ranked by -log10(FDR-adjusted p-value) are shown. Dot color indicates direction (red= upregulated genes, blue=downregulated genes); dot size represents the number of genes overlapping the GO term.

**Figure S9:**
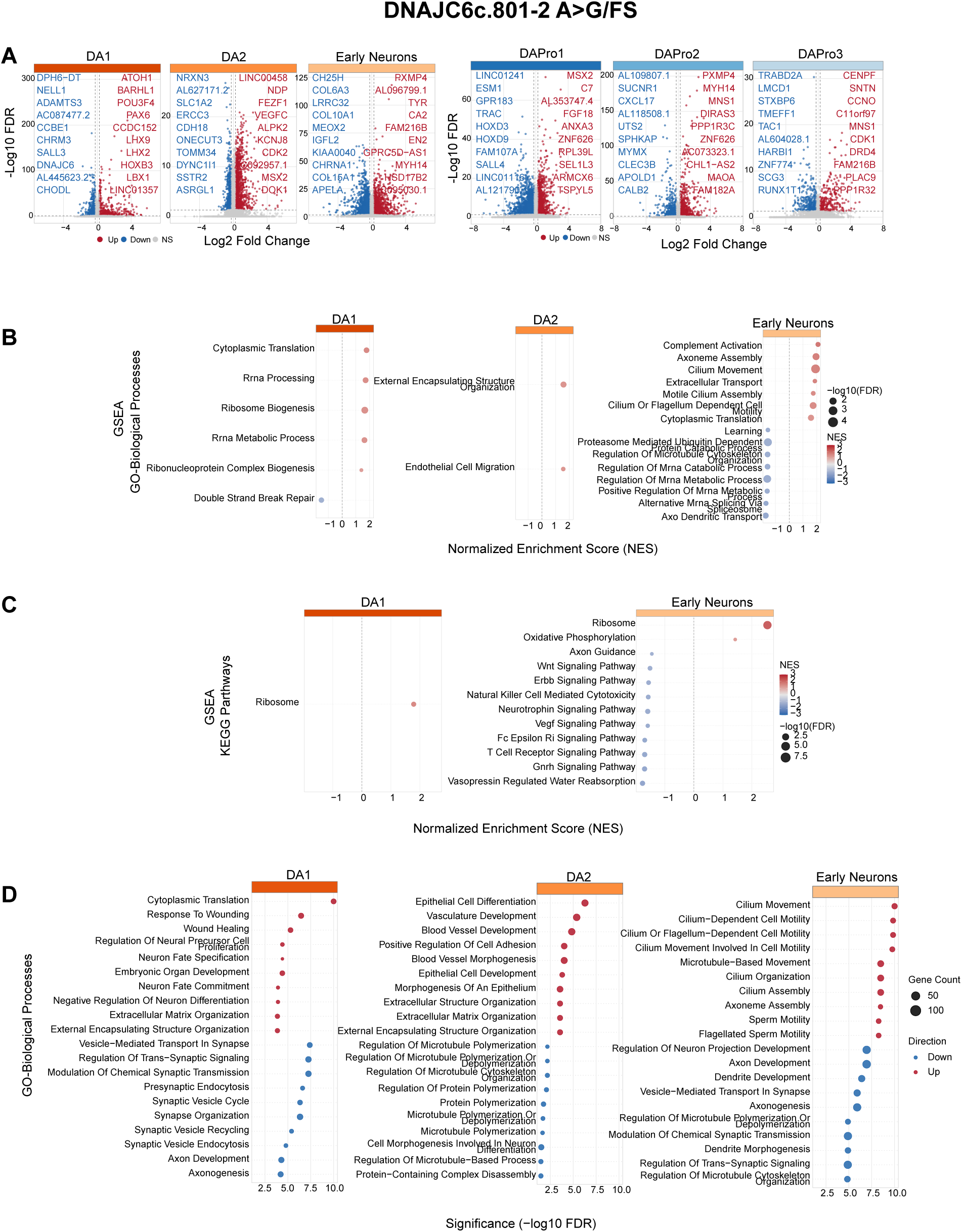
Differential expression and pathway enrichment analyses for each familial PD mutation across iSCORE-PD cell types. One supplementary figure is provided per mutation (S8: *ATP132* FS; S9: *DNAJC6* c.801-2 A>G/FS; S10: *FBXO7* R498X/FS; S11: *GBA* IVS2+1 heterozygous; S12: *GBA* IVS2+1 homozygous; S13: *LRRK2* G2019S; S14 *PINK1* Q129X; S15: *PRKN* Ex3 del; S16: *SNCA* A30P; S17: *SNCA* A53T; S18: *SYNJ1* R258Q/FS; S19: *VPS13C* A444P; S20: *VPS13C* W395C). Each figure contains four panels described below. All analyses compared each mutation genotype with the wild-type control (EWT) in the same batch. Differential expression was performed at the single-cell level using MAST. No latent variables were included, because each comparison was restricted to cells from the same batch, thereby controlling batch effects by design. EWT subclones within a given batch served as the reference population for each mutation. **A)** Differential gene expression. Volcano plots showing DE results for each annotated cell type, split into two groups: Neuronal populations and progenitor populations. X-axis shows average log2 Fold change; Y-axis shows -log10 (FDR-adjusted p-value). Upregulated genes are in red, downregulated in blue. The top 10 genes per direction per cell type ranked by log2FC are labeled. **B)** Gene Set Enrichment Analysis (GSEA)-GO biological processes: Dot plot showing GSEA results in DA neuron populations (DA1, Early Neurons; DA2 is omitted when no gene set passes the FDR <0.05 threshold). Genes were ranked using p-value and log2FC combined, capturing both effect direction and significance. **C)** GSEA, dot plots showing KEGG pathways: Same parameters used as described in B. **D)** Over-Representation Analysis (ORA), GO-BP: Dot plot showing ORA results for GO: BP terms. The top 15 terms per cell type ranked by -log10(FDR-adjusted p-value) are shown. Dot color indicates direction (red= upregulated genes, blue=downregulated genes); dot size represents the number of genes overlapping the GO term.

**Figure S10:**
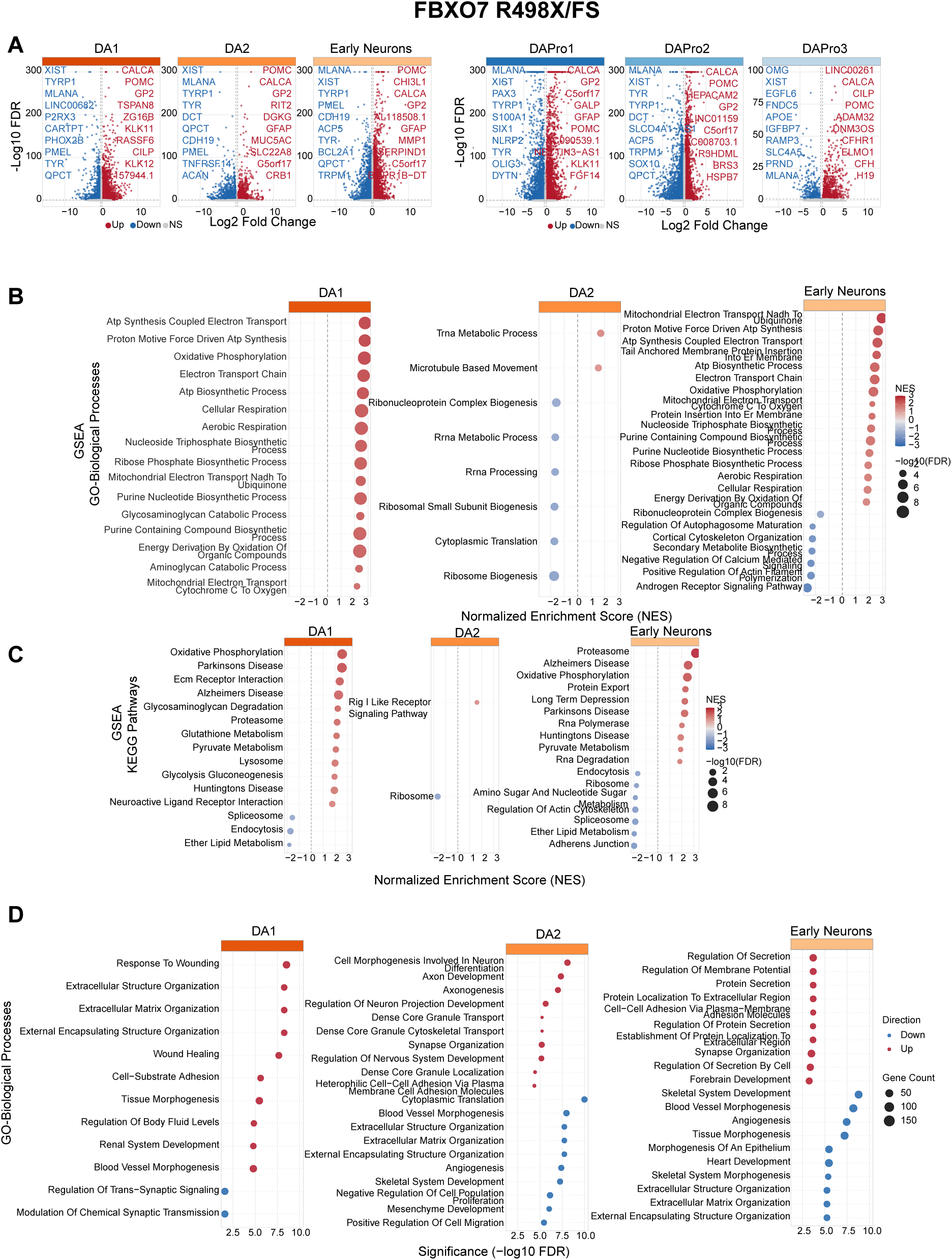
Differential expression and pathway enrichment analyses for each familial PD mutation across iSCORE-PD cell types. One supplementary figure is provided per mutation (S8: *ATP132* FS; S9: *DNAJC6* c.801-2 A>G/FS; S10: *FBXO7* R498X/FS; S11: *GBA* IVS2+1 heterozygous; S12: *GBA* IVS2+1 homozygous; S13: *LRRK2* G2019S; S14 *PINK1* Q129X; S15: *PRKN* Ex3 del; S16: *SNCA* A30P; S17: *SNCA* A53T; S18: *SYNJ1* R258Q/FS; S19: *VPS13C* A444P; S20: *VPS13C* W395C). Each figure contains four panels described below. All analyses compared each mutation genotype with the wild-type control (EWT) in the same batch. Differential expression was performed at the single-cell level using MAST. No latent variables were included, because each comparison was restricted to cells from the same batch, thereby controlling batch effects by design. EWT subclones within a given batch served as the reference population for each mutation. **A)** Differential gene expression. Volcano plots showing DE results for each annotated cell type, split into two groups: Neuronal populations and progenitor populations. X-axis shows average log2 Fold change; Y-axis shows -log10 (FDR-adjusted p-value). Upregulated genes are in red, downregulated in blue. The top 10 genes per direction per cell type ranked by log2FC are labeled. **B)** Gene Set Enrichment Analysis (GSEA)-GO biological processes: Dot plot showing GSEA results in DA neuron populations (DA1, Early Neurons; DA2 is omitted when no gene set passes the FDR <0.05 threshold). Genes were ranked using p-value and log2FC combined, capturing both effect direction and significance. **C)** GSEA, dot plots showing KEGG pathways: Same parameters used as described in B. **D)** Over-Representation Analysis (ORA), GO-BP: Dot plot showing ORA results for GO: BP terms. The top 15 terms per cell type ranked by -log10(FDR-adjusted p-value) are shown. Dot color indicates direction (red= upregulated genes, blue=downregulated genes); dot size represents the number of genes overlapping the GO term.

**Figure S11:**
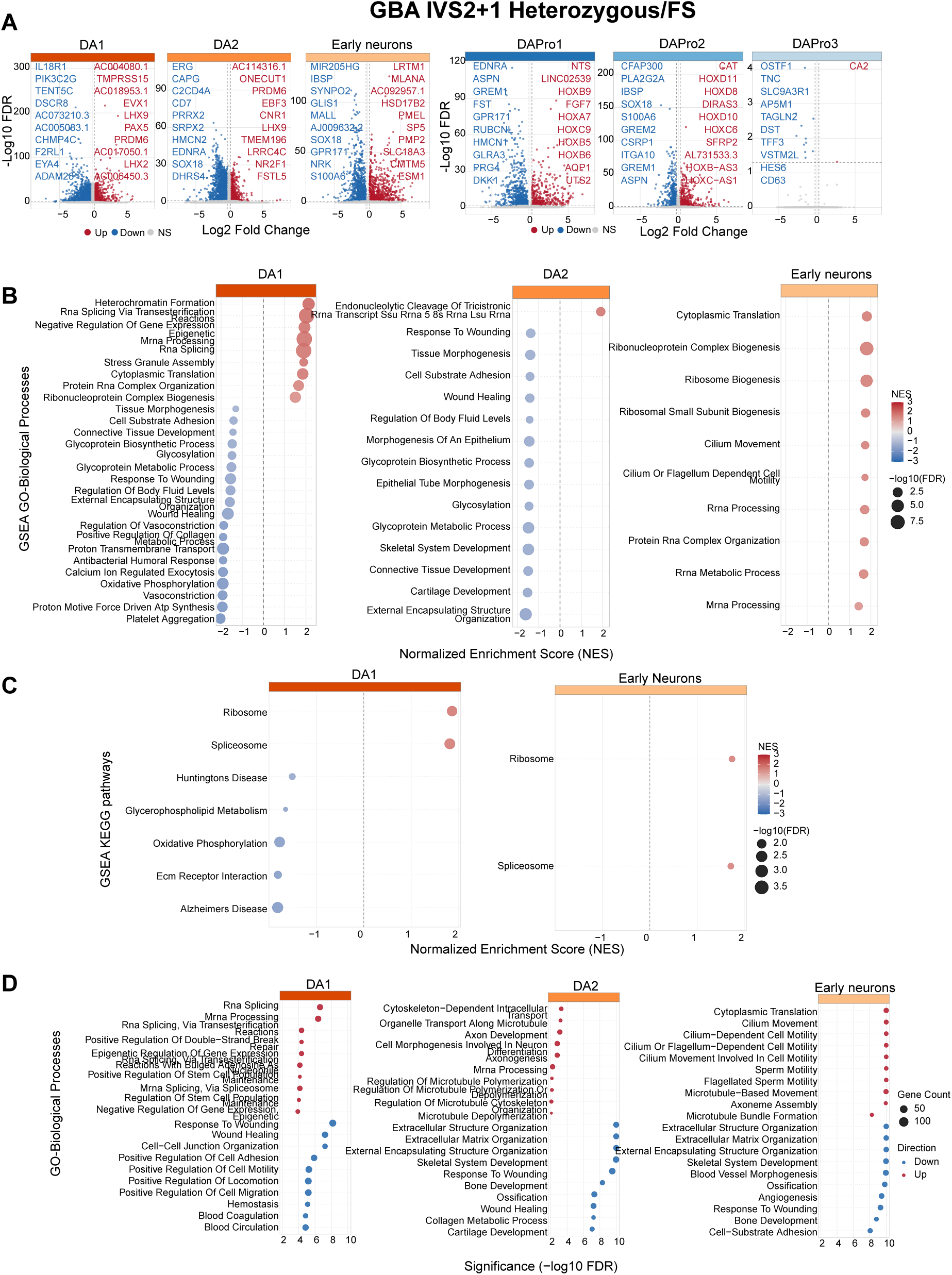
Differential expression and pathway enrichment analyses for each familial PD mutation across iSCORE-PD cell types. One supplementary figure is provided per mutation (S8: *ATP132* FS; S9: *DNAJC6* c.801-2 A>G/FS; S10: *FBXO7* R498X/FS; S11: *GBA* IVS2+1 heterozygous; S12: *GBA* IVS2+1 homozygous; S13: *LRRK2* G2019S; S14 *PINK1* Q129X; S15: *PRKN* Ex3 del; S16: *SNCA* A30P; S17: *SNCA* A53T; S18: *SYNJ1* R258Q/FS; S19: *VPS13C* A444P; S20: *VPS13C* W395C). Each figure contains four panels described below. All analyses compared each mutation genotype with the wild-type control (EWT) in the same batch. Differential expression was performed at the single-cell level using MAST. No latent variables were included, because each comparison was restricted to cells from the same batch, thereby controlling batch effects by design. EWT subclones within a given batch served as the reference population for each mutation. **A)** Differential gene expression. Volcano plots showing DE results for each annotated cell type, split into two groups: Neuronal populations and progenitor populations. X-axis shows average log2 Fold change; Y-axis shows -log10 (FDR-adjusted p-value). Upregulated genes are in red, downregulated in blue. The top 10 genes per direction per cell type ranked by log2FC are labeled. **B)** Gene Set Enrichment Analysis (GSEA)-GO biological processes: Dot plot showing GSEA results in DA neuron populations (DA1, Early Neurons; DA2 is omitted when no gene set passes the FDR <0.05 threshold). Genes were ranked using p-value and log2FC combined, capturing both effect direction and significance. **C)** GSEA, dot plots showing KEGG pathways: Same parameters used as described in B. **D)** Over-Representation Analysis (ORA), GO-BP: Dot plot showing ORA results for GO: BP terms. The top 15 terms per cell type ranked by -log10(FDR-adjusted p-value) are shown. Dot color indicates direction (red= upregulated genes, blue=downregulated genes); dot size represents the number of genes overlapping the GO term.

**Figure S12:**
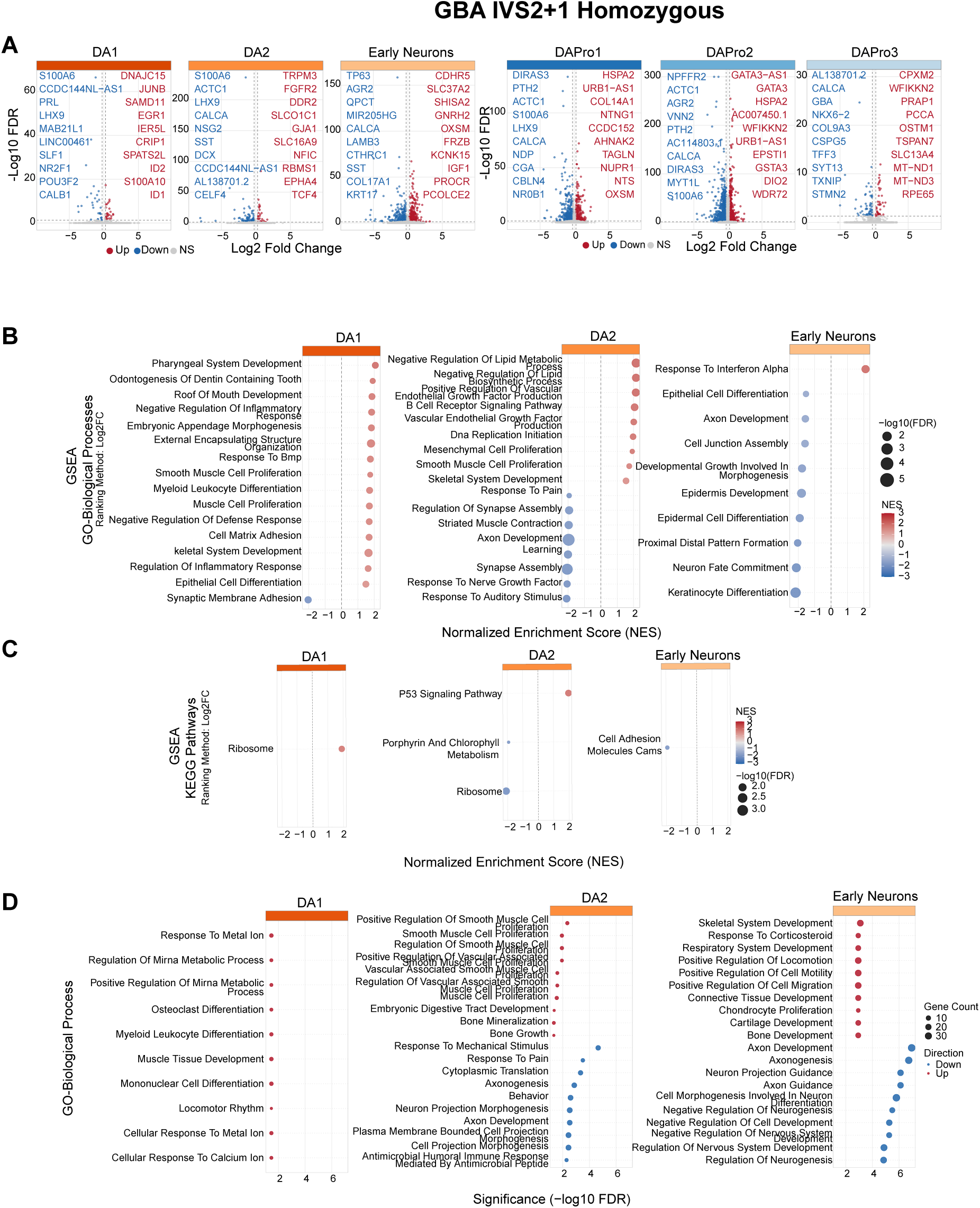
Differential expression and pathway enrichment analyses for each familial PD mutation across iSCORE-PD cell types. One supplementary figure is provided per mutation (S8: *ATP132* FS; S9: *DNAJC6* c.801-2 A>G/FS; S10: *FBXO7* R498X/FS; S11: *GBA* IVS2+1 heterozygous; S12: *GBA* IVS2+1 homozygous; S13: *LRRK2* G2019S; S14 *PINK1* Q129X; S15: *PRKN* Ex3 del; S16: *SNCA* A30P; S17: *SNCA* A53T; S18: *SYNJ1* R258Q/FS; S19: *VPS13C* A444P; S20: *VPS13C* W395C). Each figure contains four panels described below. All analyses compared each mutation genotype with the wild-type control (EWT) in the same batch. Differential expression was performed at the single-cell level using MAST. No latent variables were included, because each comparison was restricted to cells from the same batch, thereby controlling batch effects by design. EWT subclones within a given batch served as the reference population for each mutation. **A)** Differential gene expression. Volcano plots showing DE results for each annotated cell type, split into two groups: Neuronal populations and progenitor populations. X-axis shows average log2 Fold change; Y-axis shows -log10 (FDR-adjusted p-value). Upregulated genes are in red, downregulated in blue. The top 10 genes per direction per cell type ranked by log2FC are labeled. **B)** Gene Set Enrichment Analysis (GSEA)-GO biological processes: Dot plot showing GSEA results in DA neuron populations (DA1, Early Neurons; DA2 is omitted when no gene set passes the FDR <0.05 threshold). Genes were ranked using p-value and log2FC combined, capturing both effect direction and significance. **C)** GSEA, dot plots showing KEGG pathways: Same parameters used as described in B. **D)** Over-Representation Analysis (ORA), GO-BP: Dot plot showing ORA results for GO: BP terms. The top 15 terms per cell type ranked by -log10(FDR-adjusted p-value) are shown. Dot color indicates direction (red= upregulated genes, blue=downregulated genes); dot size represents the number of genes overlapping the GO term.

**Figure S13:**
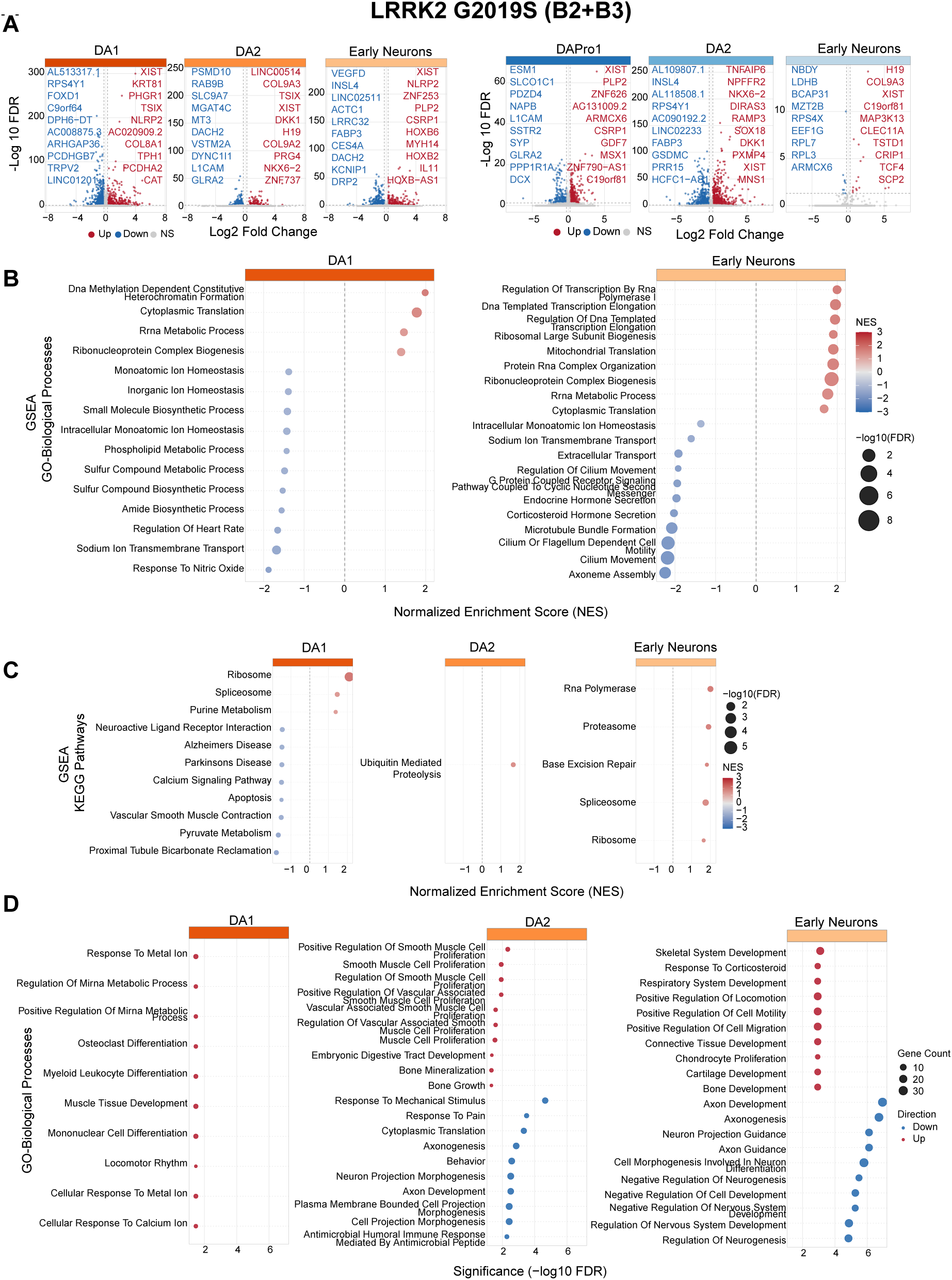
Differential expression and pathway enrichment analyses for each familial PD mutation across iSCORE-PD cell types. One supplementary figure is provided per mutation (S8: *ATP132* FS; S9: *DNAJC6* c.801-2 A>G/FS; S10: *FBXO7* R498X/FS; S11: *GBA* IVS2+1 heterozygous; S12: *GBA* IVS2+1 homozygous; S13: *LRRK2* G2019S; S14 *PINK1* Q129X; S15: *PRKN* Ex3 del; S16: *SNCA* A30P; S17: *SNCA* A53T; S18: *SYNJ1* R258Q/FS; S19: *VPS13C* A444P; S20: *VPS13C* W395C). Each figure contains four panels described below. All analyses compared each mutation genotype with the wild-type control (EWT) in the same batch. Differential expression was performed at the single-cell level using MAST. No latent variables were included, because each comparison was restricted to cells from the same batch, thereby controlling batch effects by design. EWT subclones within a given batch served as the reference population for each mutation. **A)** Differential gene expression. Volcano plots showing DE results for each annotated cell type, split into two groups: Neuronal populations and progenitor populations. X-axis shows average log2 Fold change; Y-axis shows -log10 (FDR-adjusted p-value). Upregulated genes are in red, downregulated in blue. The top 10 genes per direction per cell type ranked by log2FC are labeled. **B)** Gene Set Enrichment Analysis (GSEA)-GO biological processes: Dot plot showing GSEA results in DA neuron populations (DA1, Early Neurons; DA2 is omitted when no gene set passes the FDR <0.05 threshold). Genes were ranked using p-value and log2FC combined, capturing both effect direction and significance. **C)** GSEA, dot plots showing KEGG pathways: Same parameters used as described in B. **D)** Over-Representation Analysis (ORA), GO-BP: Dot plot showing ORA results for GO: BP terms. The top 15 terms per cell type ranked by -log10(FDR-adjusted p-value) are shown. Dot color indicates direction (red= upregulated genes, blue=downregulated genes); dot size represents the number of genes overlapping the GO term.

**Figure S14:**
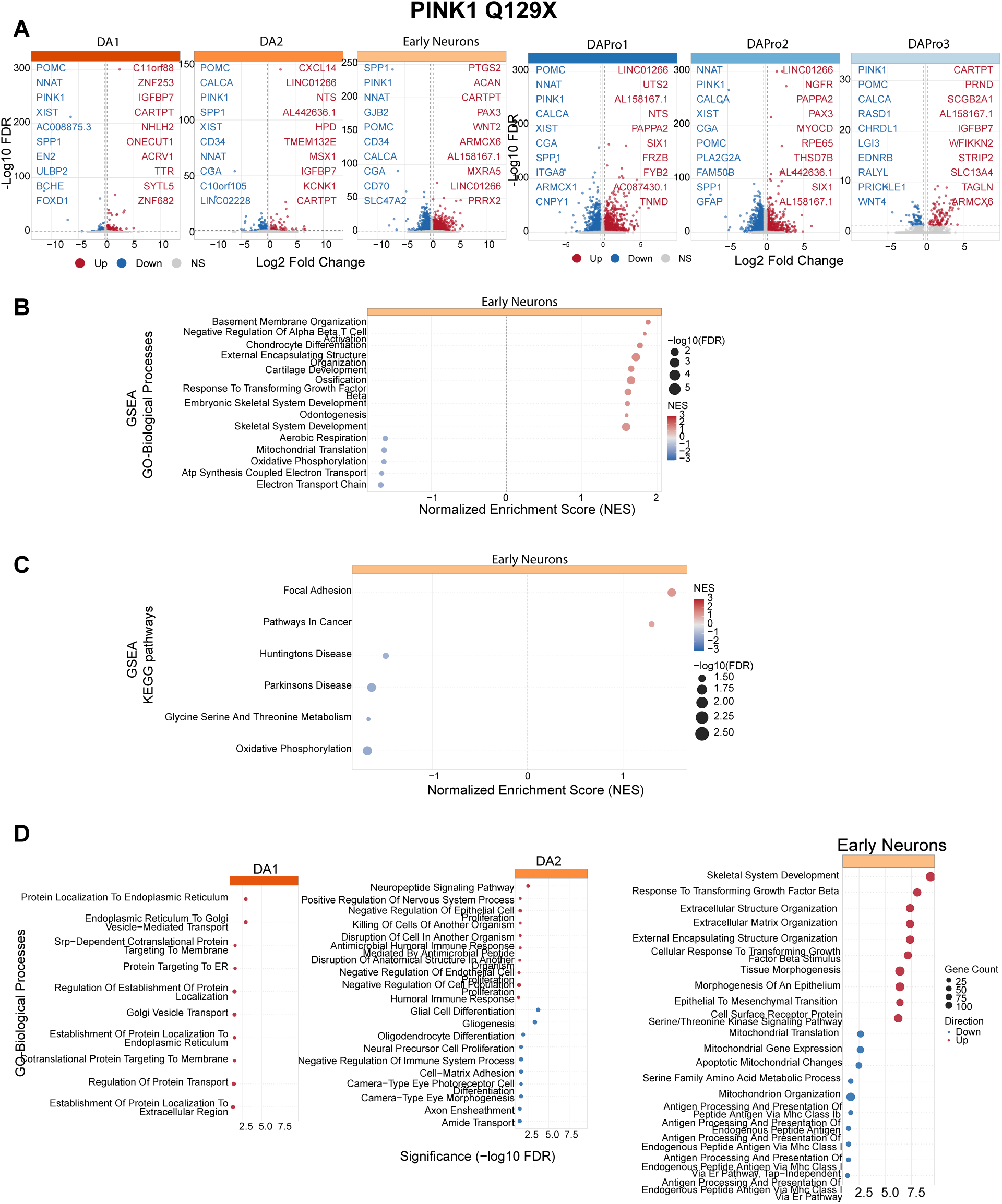
Differential expression and pathway enrichment analyses for each familial PD mutation across iSCORE-PD cell types. One supplementary figure is provided per mutation (S8: *ATP132* FS; S9: *DNAJC6* c.801-2 A>G/FS; S10: *FBXO7* R498X/FS; S11: *GBA* IVS2+1 heterozygous; S12: *GBA* IVS2+1 homozygous; S13: *LRRK2* G2019S; S14 *PINK1* Q129X; S15: *PRKN* Ex3 del; S16: *SNCA* A30P; S17: *SNCA* A53T; S18: *SYNJ1* R258Q/FS; S19: *VPS13C* A444P; S20: *VPS13C* W395C). Each figure contains four panels described below. All analyses compared each mutation genotype with the wild-type control (EWT) in the same batch. Differential expression was performed at the single-cell level using MAST. No latent variables were included, because each comparison was restricted to cells from the same batch, thereby controlling batch effects by design. EWT subclones within a given batch served as the reference population for each mutation. **A)** Differential gene expression. Volcano plots showing DE results for each annotated cell type, split into two groups: Neuronal populations and progenitor populations. X-axis shows average log2 Fold change; Y-axis shows -log10 (FDR-adjusted p-value). Upregulated genes are in red, downregulated in blue. The top 10 genes per direction per cell type ranked by log2FC are labeled. **B)** Gene Set Enrichment Analysis (GSEA)-GO biological processes: Dot plot showing GSEA results in DA neuron populations (DA1, Early Neurons; DA2 is omitted when no gene set passes the FDR <0.05 threshold). Genes were ranked using p-value and log2FC combined, capturing both effect direction and significance. **C)** GSEA, dot plots showing KEGG pathways: Same parameters used as described in B. **D)** Over-Representation Analysis (ORA), GO-BP: Dot plot showing ORA results for GO: BP terms. The top 15 terms per cell type ranked by -log10(FDR-adjusted p-value) are shown. Dot color indicates direction (red= upregulated genes, blue=downregulated genes); dot size represents the number of genes overlapping the GO term.

**Figure S15:**
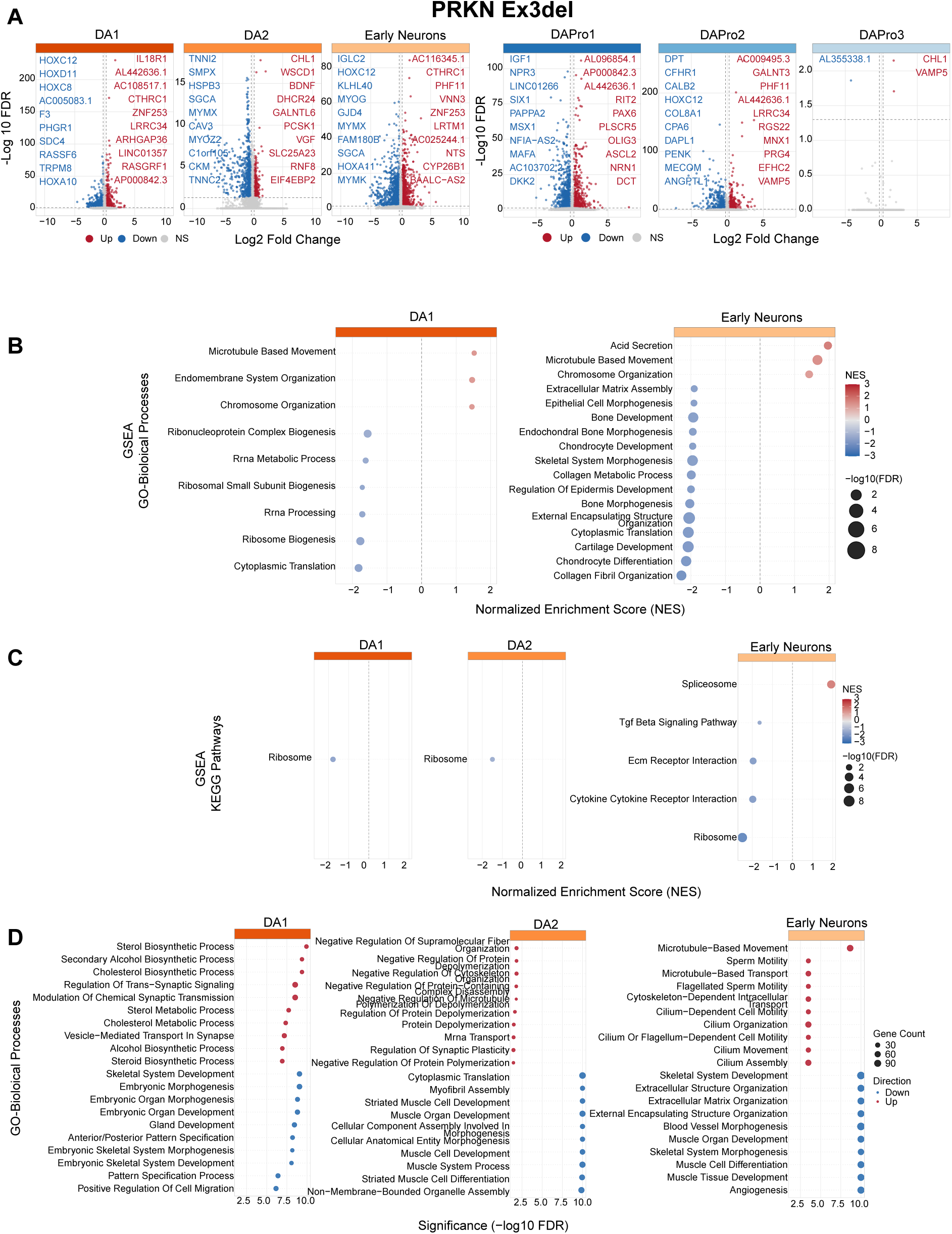
Differential expression and pathway enrichment analyses for each familial PD mutation across iSCORE-PD cell types. One supplementary figure is provided per mutation (S8: *ATP132* FS; S9: *DNAJC6* c.801-2 A>G/FS; S10: *FBXO7* R498X/FS; S11: *GBA* IVS2+1 heterozygous; S12: *GBA* IVS2+1 homozygous; S13: *LRRK2* G2019S; S14 *PINK1* Q129X; S15: *PRKN* Ex3 del; S16: *SNCA* A30P; S17: *SNCA* A53T; S18: *SYNJ1* R258Q/FS; S19: *VPS13C* A444P; S20: *VPS13C* W395C). Each figure contains four panels described below. All analyses compared each mutation genotype with the wild-type control (EWT) in the same batch. Differential expression was performed at the single-cell level using MAST. No latent variables were included, because each comparison was restricted to cells from the same batch, thereby controlling batch effects by design. EWT subclones within a given batch served as the reference population for each mutation. **A)** Differential gene expression. Volcano plots showing DE results for each annotated cell type, split into two groups: Neuronal populations and progenitor populations. X-axis shows average log2 Fold change; Y-axis shows -log10 (FDR-adjusted p-value). Upregulated genes are in red, downregulated in blue. The top 10 genes per direction per cell type ranked by log2FC are labeled. **B)** Gene Set Enrichment Analysis (GSEA)-GO biological processes: Dot plot showing GSEA results in DA neuron populations (DA1, Early Neurons; DA2 is omitted when no gene set passes the FDR <0.05 threshold). Genes were ranked using p-value and log2FC combined, capturing both effect direction and significance. **C)** GSEA, dot plots showing KEGG pathways: Same parameters used as described in B. **D)** Over-Representation Analysis (ORA), GO-BP: Dot plot showing ORA results for GO: BP terms. The top 15 terms per cell type ranked by -log10(FDR-adjusted p-value) are shown. Dot color indicates direction (red= upregulated genes, blue=downregulated genes); dot size represents the number of genes overlapping the GO term.

**Figure S16:**
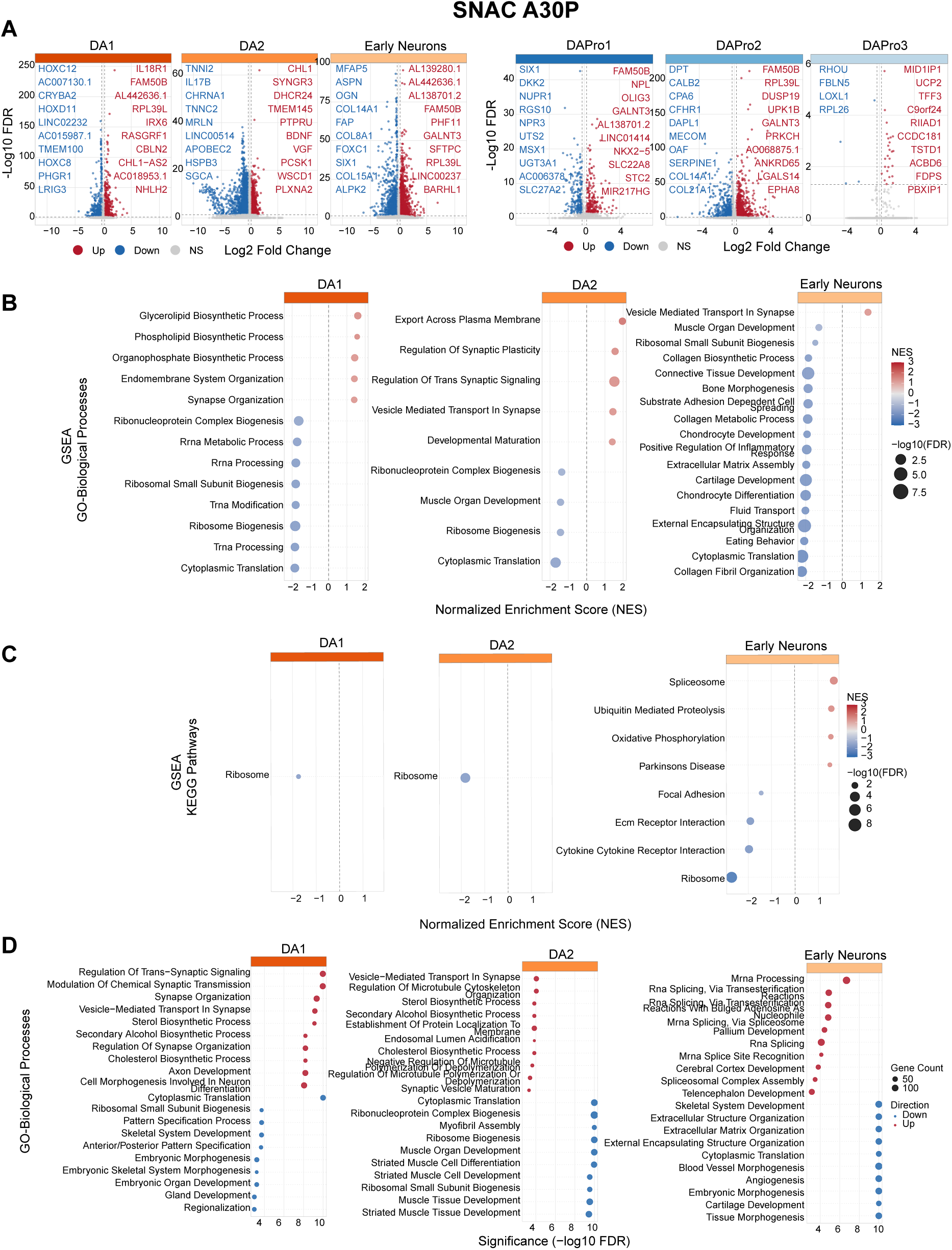
Differential expression and pathway enrichment analyses for each familial PD mutation across iSCORE-PD cell types. One supplementary figure is provided per mutation (S8: *ATP132* FS; S9: *DNAJC6* c.801-2 A>G/FS; S10: *FBXO7* R498X/FS; S11: *GBA* IVS2+1 heterozygous; S12: *GBA* IVS2+1 homozygous; S13: *LRRK2* G2019S; S14 *PINK1* Q129X; S15: *PRKN* Ex3 del; S16: *SNCA* A30P; S17: *SNCA* A53T; S18: *SYNJ1* R258Q/FS; S19: *VPS13C* A444P; S20: *VPS13C* W395C). Each figure contains four panels described below. All analyses compared each mutation genotype with the wild-type control (EWT) in the same batch. Differential expression was performed at the single-cell level using MAST. No latent variables were included, because each comparison was restricted to cells from the same batch, thereby controlling batch effects by design. EWT subclones within a given batch served as the reference population for each mutation. **A)** Differential gene expression. Volcano plots showing DE results for each annotated cell type, split into two groups: Neuronal populations and progenitor populations. X-axis shows average log2 Fold change; Y-axis shows -log10 (FDR-adjusted p-value). Upregulated genes are in red, downregulated in blue. The top 10 genes per direction per cell type ranked by log2FC are labeled. **B)** Gene Set Enrichment Analysis (GSEA)-GO biological processes: Dot plot showing GSEA results in DA neuron populations (DA1, Early Neurons; DA2 is omitted when no gene set passes the FDR <0.05 threshold). Genes were ranked using p-value and log2FC combined, capturing both effect direction and significance. **C)** GSEA, dot plots showing KEGG pathways: Same parameters used as described in B. **D)** Over-Representation Analysis (ORA), GO-BP: Dot plot showing ORA results for GO: BP terms. The top 15 terms per cell type ranked by -log10(FDR-adjusted p-value) are shown. Dot color indicates direction (red= upregulated genes, blue=downregulated genes); dot size represents the number of genes overlapping the GO term.

**Figure S17:**
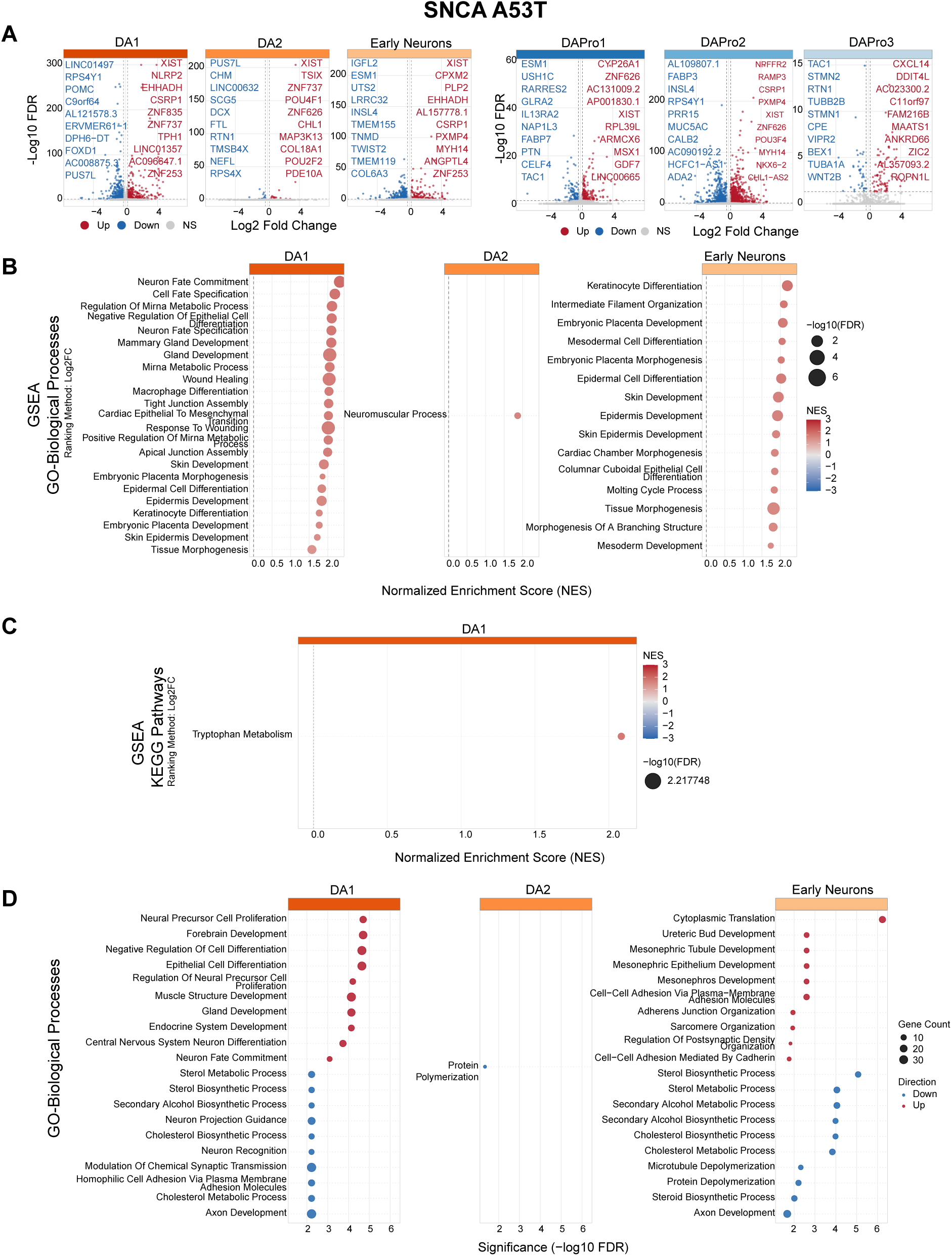
Differential expression and pathway enrichment analyses for each familial PD mutation across iSCORE-PD cell types. One supplementary figure is provided per mutation (S8: *ATP132* FS; S9: *DNAJC6* c.801-2 A>G/FS; S10: *FBXO7* R498X/FS; S11: *GBA* IVS2+1 heterozygous; S12: *GBA* IVS2+1 homozygous; S13: *LRRK2* G2019S; S14 *PINK1* Q129X; S15: *PRKN* Ex3 del; S16: *SNCA* A30P; S17: *SNCA* A53T; S18: *SYNJ1* R258Q/FS; S19: *VPS13C* A444P; S20: *VPS13C* W395C). Each figure contains four panels described below. All analyses compared each mutation genotype with the wild-type control (EWT) in the same batch. Differential expression was performed at the single-cell level using MAST. No latent variables were included, because each comparison was restricted to cells from the same batch, thereby controlling batch effects by design. EWT subclones within a given batch served as the reference population for each mutation. **A)** Differential gene expression. Volcano plots showing DE results for each annotated cell type, split into two groups: Neuronal populations and progenitor populations. X-axis shows average log2 Fold change; Y-axis shows -log10 (FDR-adjusted p-value). Upregulated genes are in red, downregulated in blue. The top 10 genes per direction per cell type ranked by log2FC are labeled. **B)** Gene Set Enrichment Analysis (GSEA)-GO biological processes: Dot plot showing GSEA results in DA neuron populations (DA1, Early Neurons; DA2 is omitted when no gene set passes the FDR <0.05 threshold). Genes were ranked using p-value and log2FC combined, capturing both effect direction and significance. **C)** GSEA, dot plots showing KEGG pathways: Same parameters used as described in B. **D)** Over-Representation Analysis (ORA), GO-BP: Dot plot showing ORA results for GO: BP terms. The top 15 terms per cell type ranked by -log10(FDR-adjusted p-value) are shown. Dot color indicates direction (red= upregulated genes, blue=downregulated genes); dot size represents the number of genes overlapping the GO term.

**Figure S18:**
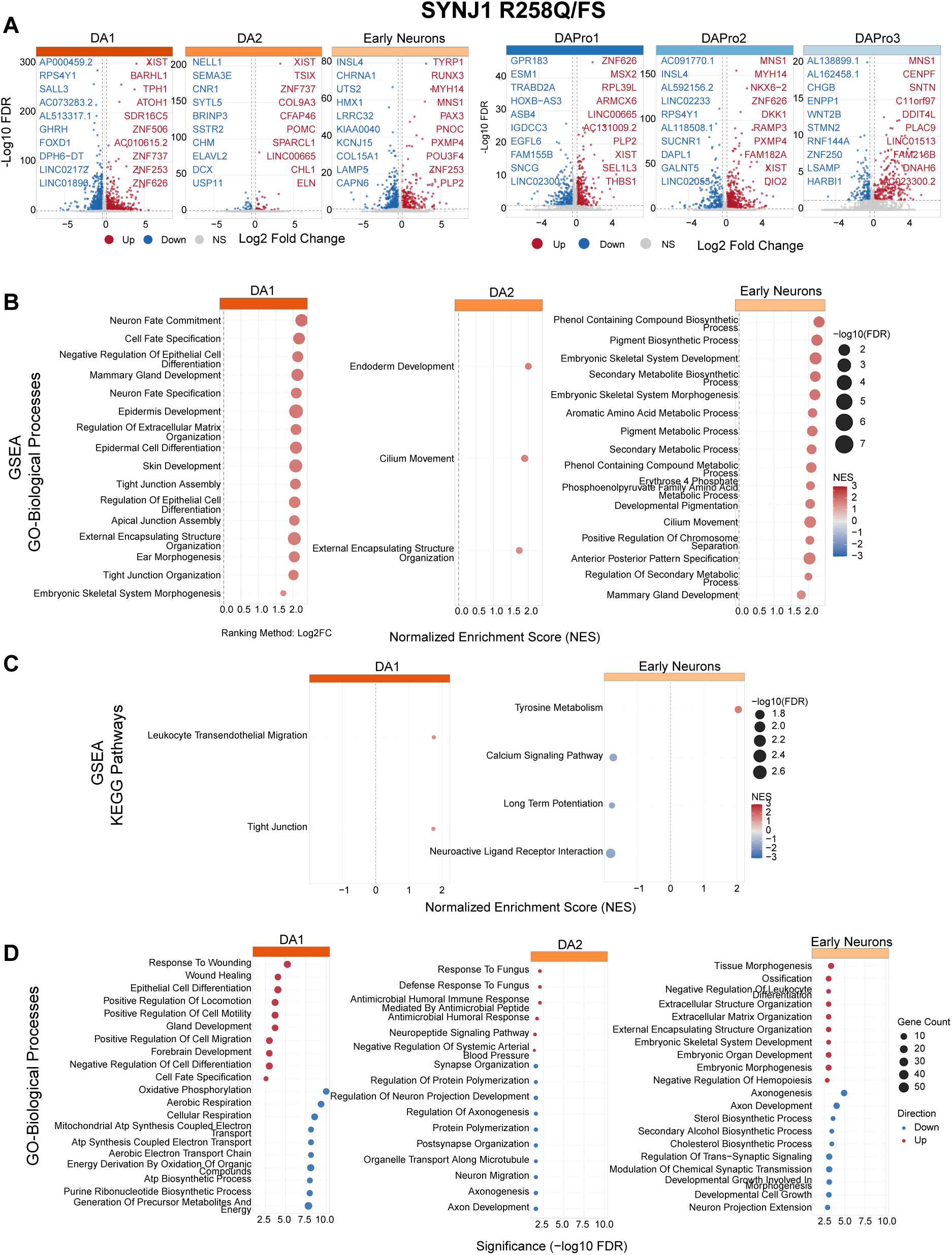
Differential expression and pathway enrichment analyses for each familial PD mutation across iSCORE-PD cell types. One supplementary figure is provided per mutation (S8: *ATP132* FS; S9: *DNAJC6* c.801-2 A>G/FS; S10: *FBXO7* R498X/FS; S11: *GBA* IVS2+1 heterozygous; S12: *GBA* IVS2+1 homozygous; S13: *LRRK2* G2019S; S14 *PINK1* Q129X; S15: *PRKN* Ex3 del; S16: *SNCA* A30P; S17: *SNCA* A53T; S18: *SYNJ1* R258Q/FS; S19: *VPS13C* A444P; S20: *VPS13C* W395C). Each figure contains four panels described below. All analyses compared each mutation genotype with the wild-type control (EWT) in the same batch. Differential expression was performed at the single-cell level using MAST. No latent variables were included, because each comparison was restricted to cells from the same batch, thereby controlling batch effects by design. EWT subclones within a given batch served as the reference population for each mutation. **A)** Differential gene expression. Volcano plots showing DE results for each annotated cell type, split into two groups: Neuronal populations and progenitor populations. X-axis shows average log2 Fold change; Y-axis shows -log10 (FDR-adjusted p-value). Upregulated genes are in red, downregulated in blue. The top 10 genes per direction per cell type ranked by log2FC are labeled. **B)** Gene Set Enrichment Analysis (GSEA)-GO biological processes: Dot plot showing GSEA results in DA neuron populations (DA1, Early Neurons; DA2 is omitted when no gene set passes the FDR <0.05 threshold). Genes were ranked using p-value and log2FC combined, capturing both effect direction and significance. **C)** GSEA, dot plots showing KEGG pathways: Same parameters used as described in B. **D)** Over-Representation Analysis (ORA), GO-BP: Dot plot showing ORA results for GO: BP terms. The top 15 terms per cell type ranked by -log10(FDR-adjusted p-value) are shown. Dot color indicates direction (red= upregulated genes, blue=downregulated genes); dot size represents the number of genes overlapping the GO term.

**Figure S19:**
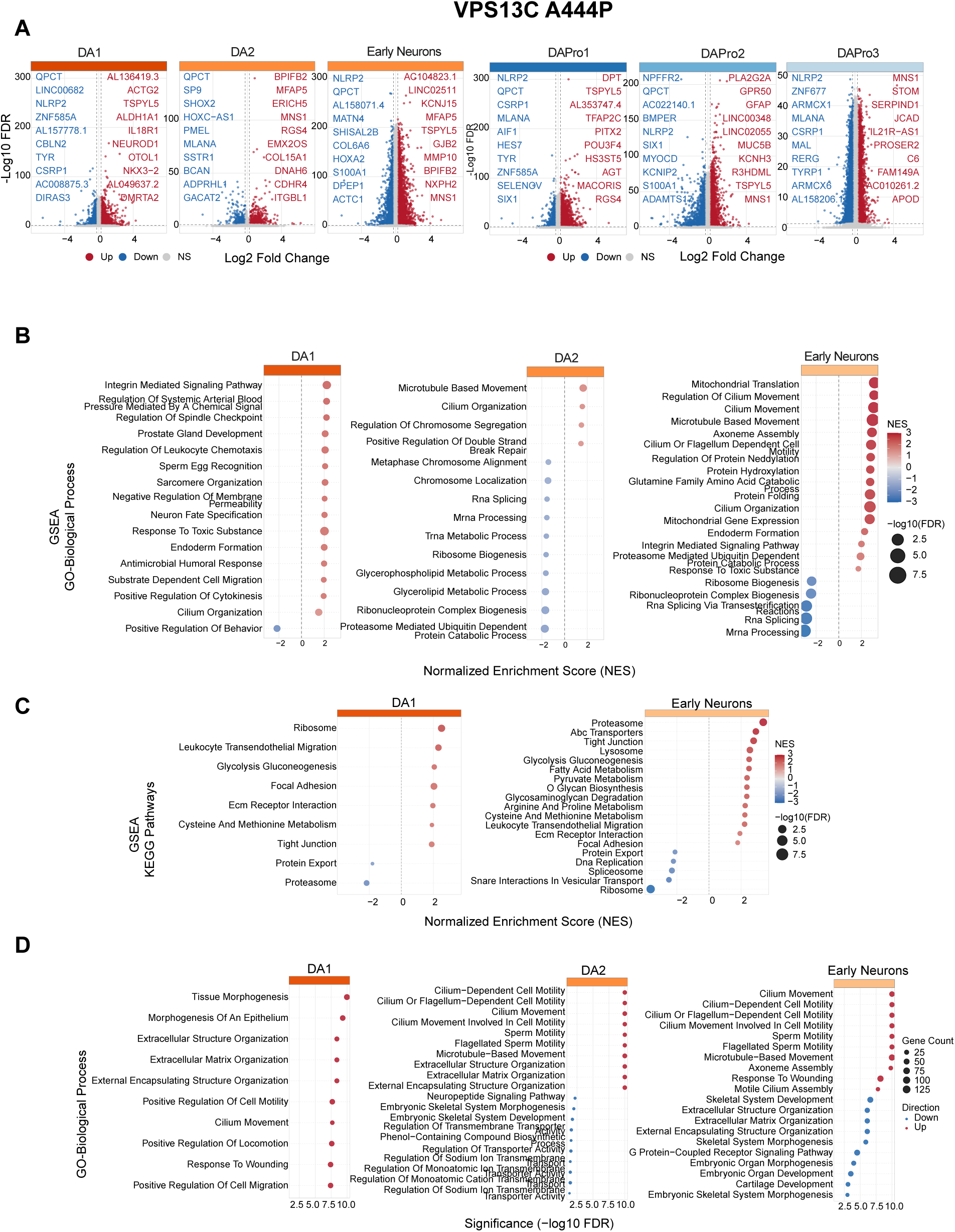
Differential expression and pathway enrichment analyses for each familial PD mutation across iSCORE-PD cell types. One supplementary figure is provided per mutation (S8: *ATP132* FS; S9: *DNAJC6* c.801-2 A>G/FS; S10: *FBXO7* R498X/FS; S11: *GBA* IVS2+1 heterozygous; S12: *GBA* IVS2+1 homozygous; S13: *LRRK2* G2019S; S14 *PINK1* Q129X; S15: *PRKN* Ex3 del; S16: *SNCA* A30P; S17: *SNCA* A53T; S18: *SYNJ1* R258Q/FS; S19: *VPS13C* A444P; S20: *VPS13C* W395C). Each figure contains four panels described below. All analyses compared each mutation genotype with the wild-type control (EWT) in the same batch. Differential expression was performed at the single-cell level using MAST. No latent variables were included, because each comparison was restricted to cells from the same batch, thereby controlling batch effects by design. EWT subclones within a given batch served as the reference population for each mutation. **A)** Differential gene expression. Volcano plots showing DE results for each annotated cell type, split into two groups: Neuronal populations and progenitor populations. X-axis shows average log2 Fold change; Y-axis shows -log10 (FDR-adjusted p-value). Upregulated genes are in red, downregulated in blue. The top 10 genes per direction per cell type ranked by log2FC are labeled. **B)** Gene Set Enrichment Analysis (GSEA)-GO biological processes: Dot plot showing GSEA results in DA neuron populations (DA1, Early Neurons; DA2 is omitted when no gene set passes the FDR <0.05 threshold). Genes were ranked using p-value and log2FC combined, capturing both effect direction and significance. **C)** GSEA, dot plots showing KEGG pathways: Same parameters used as described in B. **D)** Over-Representation Analysis (ORA), GO-BP: Dot plot showing ORA results for GO: BP terms. The top 15 terms per cell type ranked by -log10(FDR-adjusted p-value) are shown. Dot color indicates direction (red= upregulated genes, blue=downregulated genes); dot size represents the number of genes overlapping the GO term.

**Figure S20:**
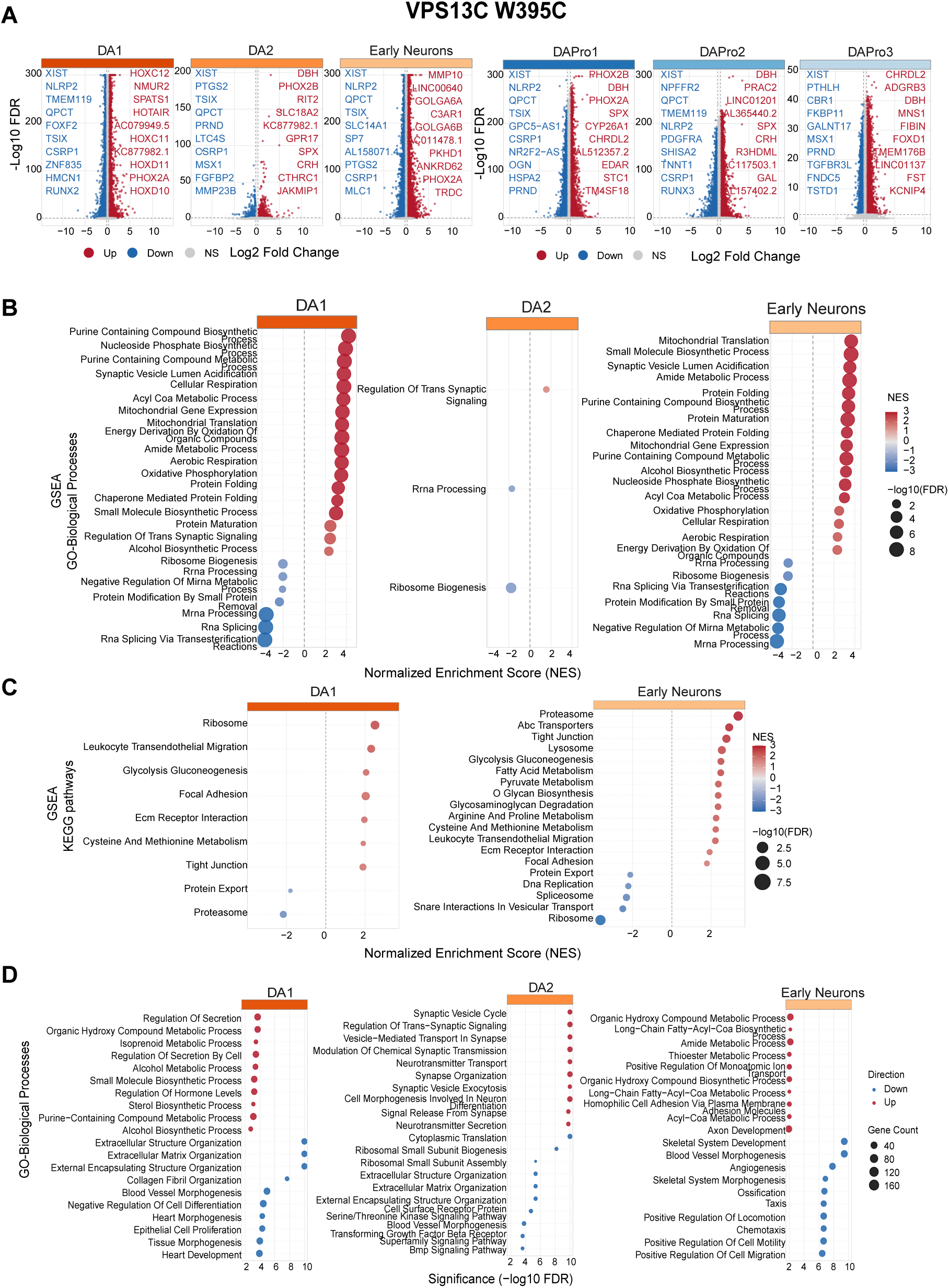
Differential expression and pathway enrichment analyses for each familial PD mutation across iSCORE-PD cell types. One supplementary figure is provided per mutation (S8: *ATP132* FS; S9: *DNAJC6* c.801-2 A>G/FS; S10: *FBXO7* R498X/FS; S11: *GBA* IVS2+1 heterozygous; S12: *GBA* IVS2+1 homozygous; S13: *LRRK2* G2019S; S14 *PINK1* Q129X; S15: *PRKN* Ex3 del; S16: *SNCA* A30P; S17: *SNCA* A53T; S18: *SYNJ1* R258Q/FS; S19: *VPS13C* A444P; S20: *VPS13C* W395C). Each figure contains four panels described below. All analyses compared each mutation genotype with the wild-type control (EWT) in the same batch. Differential expression was performed at the single-cell level using MAST. No latent variables were included, because each comparison was restricted to cells from the same batch, thereby controlling batch effects by design. EWT subclones within a given batch served as the reference population for each mutation. **A)** Differential gene expression. Volcano plots showing DE results for each annotated cell type, split into two groups: Neuronal populations and progenitor populations. X-axis shows average log2 Fold change; Y-axis shows -log10 (FDR-adjusted p-value). Upregulated genes are in red, downregulated in blue. The top 10 genes per direction per cell type ranked by log2FC are labeled. **B)** Gene Set Enrichment Analysis (GSEA)-GO biological processes: Dot plot showing GSEA results in DA neuron populations (DA1, Early Neurons; DA2 is omitted when no gene set passes the FDR <0.05 threshold). Genes were ranked using p-value and log2FC combined, capturing both effect direction and significance. **C)** GSEA, dot plots showing KEGG pathways: Same parameters used as described in B. **D)** Over-Representation Analysis (ORA), GO-BP: Dot plot showing ORA results for GO: BP terms. The top 15 terms per cell type ranked by -log10(FDR-adjusted p-value) are shown. Dot color indicates direction (red= upregulated genes, blue=downregulated genes); dot size represents the number of genes overlapping the GO term.

